# Compartmentalized Biosynthesis of Mycophenolic Acid

**DOI:** 10.1101/524025

**Authors:** Wei Zhang, Lei Du, Zepeng Qu, Xingwang Zhang, Fengwei Li, Zhong Li, Feifei Qi, Xiao Wang, Yuanyuan Jiang, Ping Men, Jingran Sun, Shaona Cao, Ce Geng, Fengxia Qi, Xiaobo Wan, Changning Liu, Shengying Li

## Abstract

Mycophenolic acid (MPA) from filamentous fungi is the first natural product antibiotic in human history and a first-line immunosuppressive drug for organ transplantations and autoimmune diseases. However, its biosynthetic mechanisms have remained a long-standing mystery. Here, we elucidate the MPA biosynthetic pathway that features both compartmentalized enzymatic steps and unique cooperation between biosynthetic and *β*-oxidation catabolism machineries based on targeted gene inactivation, feeding experiments in heterologous expression hosts, enzyme functional characterization and kinetic analysis, and microscopic observation of protein subcellular localization. Besides identification of the oxygenase MpaB’ as the long-sought key enzyme responsible for the oxidative cleavage of sesquiterpene side chain, we reveal the intriguing pattern of compartmentalization for the MPA biosynthetic enzymes, including the cytosolic polyketide synthase MpaC’ and *O*-methyltransferase MpaG’, the Golgi apparatus-associated prenyltransferase MpaA’, the endoplasmic reticulum-bound oxygenase MpaB’ and P450-hydrolase fusion enzyme MpaDE’, and the peroxisomal acyl-CoA hydrolase MpaH’. The whole pathway is elegantly co-mediated by these compartmentalized enzymes, together with the peroxisomal *β*-oxidation machinery. Beyond characterizing the remaining outstanding steps of the MPA biosynthetic pathway, our study highlights the importance of considering subcellular contexts and the broader cellular metabolism in natural product biosynthesis.

**Significance Statement:** Here we elucidate the full biosynthetic pathway of the fungal natural product mycophenolic acid (MPA), which represents an unsolved mystery for decades. Besides the intriguing enzymatic mechanisms, we reveal that the MPA biosynthetic enzymes are elegantly compartmentalized; and the subcellular localization of the acyl-CoA hydrolase MpaH’ in peroxisomes is required for the unique cooperation between biosynthetic and *β*-oxidation catabolism machineries. This work highlights the importance of a cell biology perspective for understanding the unexplored organelle-associated essential catalytic mechanisms in natural product biosynthesis of fungi and other higher organisms. The insights provided by our work will also benefit future efforts for both industrial strain improvement and novel drug development.

## Main Text

Mycophenolic acid (MPA), which was discovered from *Penicillium brevicompactum* in 1893 (1), is the first natural product antibiotic in human history. Today, its different active forms (*e.g.*, CellCept^®^ by Roche and Myfortic^®^ by Novartis) have annual sales over $1 billion, owing to their wide use as first-line immunosuppressive drugs to control immunologic rejection during organ transplantations and to treat autoimmune diseases (2, 3). Mechanistically, MPA inhibits inosine-5’-monophosphate dehydrogenase; this enzyme catalyzes a known pathway-regulating step of guanine synthesis, which is essential for lymphocyte proliferation (4). MPA is a tetraketide–terpenoid (TKTP) compound; this family comprises various chemical structures with a wide spectrum of biological activities (*SI Appendix*, Fig. S1), and TKTPs are the largest class of meroterpenoids produced by filamentous fungi (5). Despite both MPA’s status as the first natural product antibiotic and the growing number of studies reporting the characterization of fungal TKTP biosynthetic pathways (5–8), a full understanding of MPA biosynthesis has remained elusive for more than a century. This knowledge gap is especially conspicuous when one considers that the industrial fermentation of MPA has been established for decades and its structure is not particularly complex, with a full synthesis having been demonstrated by 1969 (9).

The first insights into MPA biosynthesis, which were gained more than four decades ago from culture feeding studies using synthetic radioactive isotope labeling precursors, revealed its skeleton is derived from 5-methylorsellinic acid (5-MOA) and farnesyl pyrophosphate (FPP), and > a putative oxidative cleavage of the sesquiterpene (C_15_) side chain (10–12). The *C*-methyl group at C6 and the *O*-methyl group at C5 were proposed to originate from *S*-adenosyl-L-methionine (SAM) (10, 13). However, the genetic and enzymological bases for MPA biosynthesis remained obscure until the recent independent discoveries of three analogous MPA biosynthetic gene clusters (*SI Appendix*, Fig. S2) (14–16). Upon identification of these clusters, a sub-set of the MPA biosynthetic pathway steps have been revealed through the functional characterization of three biosynthetic enzymes: MpaC (14, 17) and MpaDE (18) from *P. brevicompactum* IBT23078 as well as MpaG' from *P. brevicompactum* NRRL864 (*Pb*_864_) (15) (Fig. 1).

**Fig. 1.**
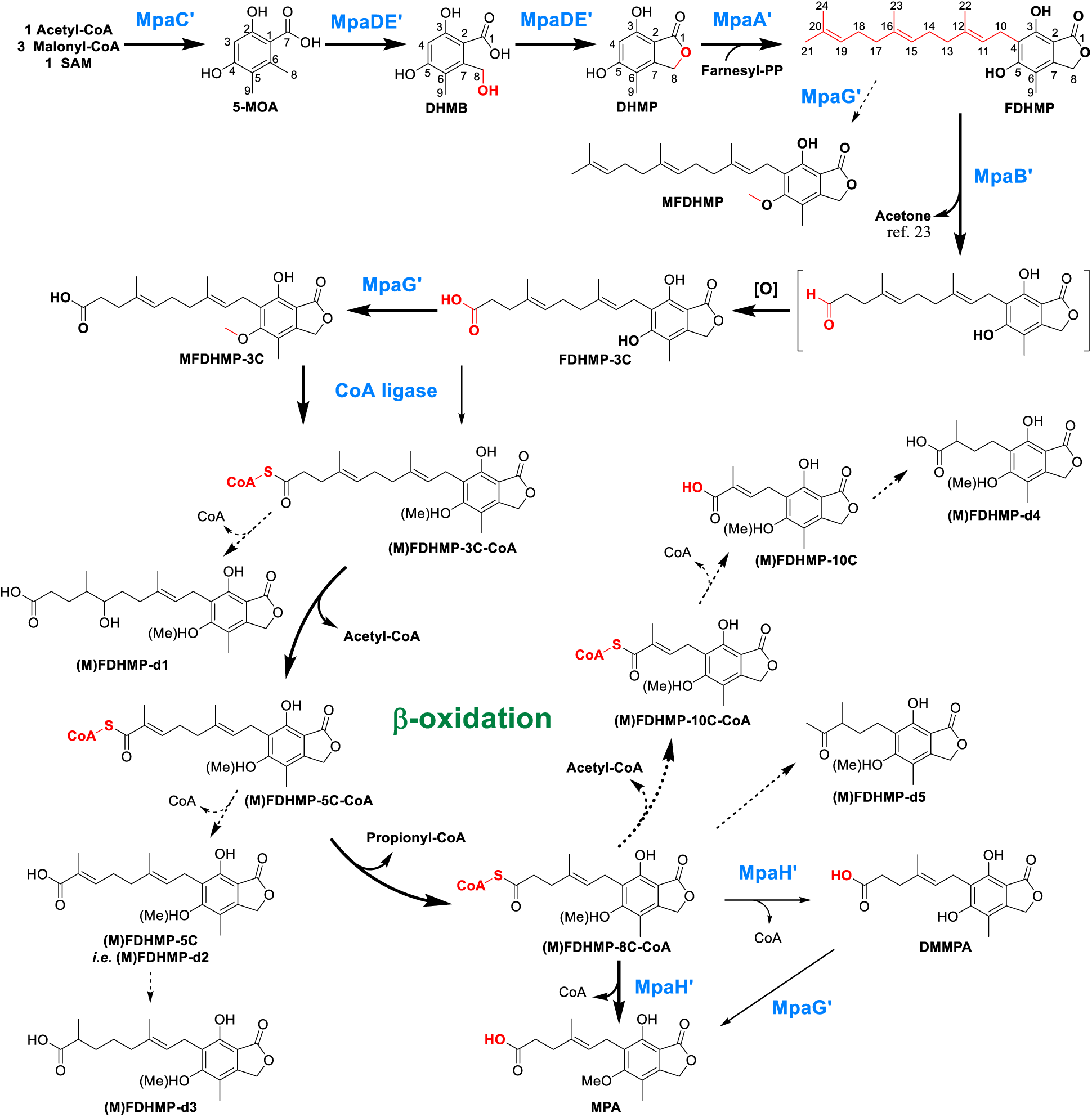
The MPA biosynthetic pathway. Bold, plain, and dashed arrows indicate the major, minor, and shunt pathways, respectively. The newly installed functional groups are colored in red.

Using examples from the *mpa’* gene cluster of *Pb*_864_ (*SI Appendix*, Table S1) to illustrate the present state of knowledge about MPA biosynthesis (Fig. 1): it is known that the MpaC' enzyme is a polyketide synthase (PKS) that catalyzes the formation of its 5-MOA product from one acetyl-CoA molecule, three malonyl-CoA units, and one SAM molecule. The fascinating MpaDE' enzyme comprises a cytochrome P450 domain (MpaD') fused to a hydrolase domain (MpaE') and catalyzes both the formation of 3,5-dihydroxy-7-(hydroxymethyl)-6-methylbenzoic acid (DHMB) via the C4 hydroxylation activity of MpaD’ and the subsequent intramolecular dehydration by MpaE’ to produce 3,5-dihydroxy-6-methylphthalide (DHMP). The following biosynthetic steps lack experimental confirmation, but it has been proposed that DHMP is next farnesylated by the prenyltransferase MpaA' to yield the isolatable intermediate 4-farnesyl-3,5-dihydroxy-6-methylphtalide (FDHMP) (19–22). The biosynthetic steps between FDHMP and the penultimate product demethylmycophenolic acid (DMMPA) have been speculated (14, 23) but remain uncharacterized, while the final step is known to be the *O*-methylation of DMMPA’s C5 hydroxy group by the *O*-methyltransferase MpaG' (15) to yield the final product MPA.

Our exploration of MPA biosynthesis in the present study started with our efforts to experimentally confirm that the putative prenyltransferase MpaA' can indeed add a farnesyl group to DHMP to form FDHMP. Following the recombinant-*mpaDE'-*expression and 5-MOA-feeding based generation, purification, and structural confirmation of the hypothetical MpaA' substrate DHMP (*SI Appendix*, Figs. S3-S6 and Tables S2-S5), we used the popular auxotrophic *Aspergillus oryzae* M-2-3 (*Ao*_M-2-3_) strain as the heterologous expression host to conduct *in vivo* assays of MpaA' activity (note that attempts to heterologously express this transmembrane protein (*SI Appendix*, Fig. S7) in *Escherichia coli* and *Sacchromyces cerevisiae* were unsuccessful). When purified DHMP (20 mg/L) was fed to a maltose-induced culture of the pTAex3-*mpaA'* (*SI Appendix*, Figs. S3-S4 and Table S2) harboring strain *Ao*_M-2-3_-*mpaA’* (*SI Appendix*, Table S3), the precursor DHMP was completely converted into a much more hydrophobic product within 5 d (Fig. 2*A*), and analysis of high-resolution mass spectrometry (HRMS) data suggested that the molecular formula of this product was C_24_H_32_O_4_ (*SI Appendix*, Fig. S8 and Table S4), which is consistent with that of FDHMP. NMR analyses further structurally confirmed the product as FDHMP^20-22^. (*SI Appendix*, Figs. S9-S10 and Table S6).

**Fig. 2.**
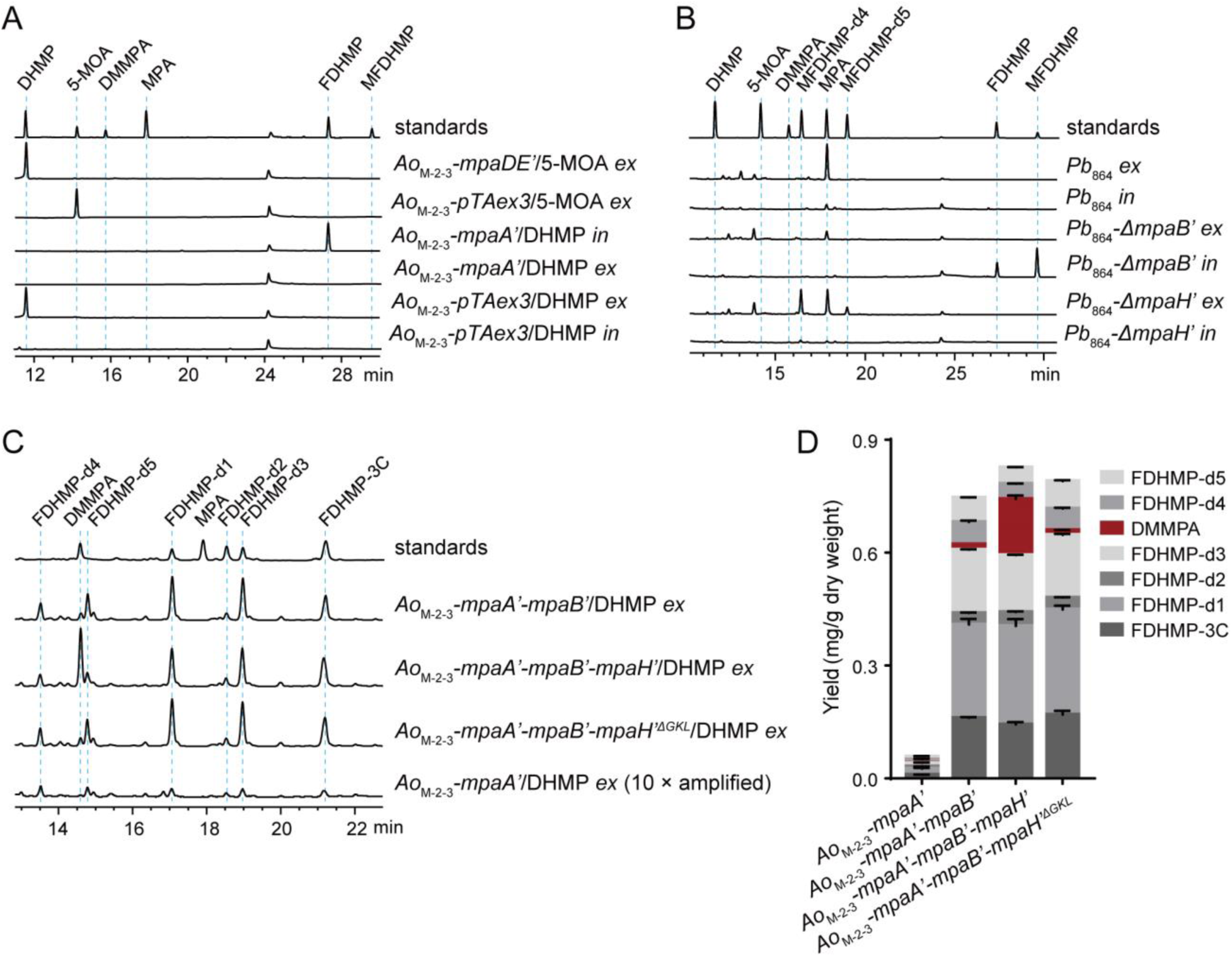
HPLC analysis (254 nm) of *Aspergillus oryzae* M-2-3 (*Ao*_M-2-3_) precursor feeding experiments and *Penicillium brevicompactum* NRRL 864 (*Pb*_864_) knockout mutants (*ex*: the extracellular extracts; *in*: the intracellular extracts). (*A*). Precursor feeding studies of the *Ao*_M-2-3_ mutant strains that express a single *mpa’* gene. (*B*). Product profiles of the wild type and mutant *Pb*_864_ strains. (*C*). Precursor feeding studies of the *Ao*_M-2-3_ mutant strains with co-expression of *mpa’* genes. (*D*). Quantitative analysis of the production of DMMPA and FDHMP derivatives.

Notably, FDHMP was only detected in the extracts prepared from mycelia, but not the fermentation broth (Fig. 2*A*), suggesting that FDHMP might have difficulty in passing through the fungal cell membrane, owing perhaps to its presumably membrane-embedded nature (like FPP) (24). Thus, MpaA' does catalyze the transfer of a farnesyl group from FPP to DHMP via C–C bond formation. However, 5-MOA was not farnesylated in a similar feeding experiment (*SI Appendix*, Fig. S11), highlighting the high substrate specificity of MpaA’.

Having experimentally confirmed the farnesyl-transfer activity of MpaA’, we next attempted to unravel the long-standing biosynthetic mystery of which biomolecule(s) are responsible for the assumed oxidative cleavage of the central double bond in the sesquiterpene chain of FDHMP (*i.e.*, the C15=C16 olefin) (1, 14, 20-23). Additional genes of the *mpa’* gene cluster include *mpaF’*, *mpaB’*, and *mpaH’*; we did not pursue MpaF’ as a candidate for oxidative cleavage functionality because it is known to be an inosine-5’-monophosphate dehydrogenase involved in the self-resistance of MPA producing strains (14,25). To investigate the unknown functions of MpaB’ and MpaH’, we used a split-marker recombination strategy (26) to singly knock out *mpaB’* or *mpaH’* in *Pb*_864_ (*SI Appendix*, Fig. S12) to produce the inactivation mutants *Pb*_864_-*ΔmpaB’* and *Pb*_864_-*ΔmpaH’* (*SI Appendix*, Table S3).

Compared to *Pb*_864_, *Pb*_864_-*ΔmpaB’* produced a dramatically decreased amounts of MPA, but this strain accumulated a significant amount of FDHMP in its mycelia during a 7 d cultivation using potato dextrose broth (Fig. 2*B*). Additionally, a new product was detected in the intracellular fraction of *Pb*_864_-*ΔmpaB’*, whose structure was determined as 5-*O*-methyl-FDHMP (MFDHMP, Fig. 1) by HRMS (*SI Appendix*, Fig. S8 and Table S4) and NMR analyses (*SI Appendix*, Figs. S13-S17 and Table S6). We reason that the inactivation of MpaB’ blocked the normal conversion of FDHMP, which was methylated to MFDHMP (likely by MpaG’, which has been reported to display considerable substrate flexibility (15)). Of note, the small amount of MPA produced by *Pb*_864_-*ΔmpaB’* (Fig. 2*B*) suggests the existence of minor compensating enzymatic activity for MpaB’ in *Pb*_864_.

To further elucidate the functionality of MpaB’, FDHMP (20 mg/L) was fed to an induction culture of *Ao*_M-2-3_-*mpaB’* (*SI Appendix*, Table S3). Surprisingly, no obvious products were detected (*SI Appendix*, Fig. S18). We reason that this negative result might be due to the difficulty for FDHMP to enter the intracellular space, which is supported by our earlier observation that FDHMP was not secreted outside of *Ao*_M-2-3_*-mpaA’* cells (Fig. 2*A*). To overcome this issue, the recombinant strain *Ao*_M-2-3_-*mpaA’-mpaB’* was generated and cultured in CD medium supplemented with maltose to induce the co-expression of MpaA’ and MpaB’ for 3 d, to which DHMP (20 mg/L) was added. Upon an additional 5 d cultivation, an intermediate with three fewer carbon atoms than FDHMP was observed (Fig. 2*C*), purified, and structurally identified as FDHMP-3C (Fig. 1; *SI Appendix*, Figs. S8, S19-S23, and Tables S4 and S7). Interestingly, FDHMP-3C was previously proposed as a putative intermediate *en route* to MPA (*SI Appendix*, Fig. S24) (12, 21).

We also found that *Ao*_M-2-3_-*mpaA’-mpaB’* produced additional derivatives with UV absorption spectra similar to those of FDHMP and FDHMP-3C (Fig. 2*C*), which presumably derived from FDHMP; these were therefore deemed FDHMP-d1–d5. Structural determination (*SI Appendix*, Figs. S8, S25-S39, and Tables S4, S7-S8) showed that FDHMP-d1–d5 appear to be chain-shortening intermediates of FDHMP-3C that also bear some additional modifications, suggesting a possible biodegradation pathway through which FDHMP-3C may undergo a *β*-oxidation process in which a C_2_/C_3_ unit can be successively lost over repeated rounds (Fig. 1). Strikingly, a small amount of DMMPA was also detected, giving an initial hint that *mpaH’* may not be a required gene for DMMPA production.

These results collectively establish that it is MpaB’ which functions as an oxygenase to mediate the oxidative cleavage of the C_19_=C_20_ double bond in FDHMP to yield FDHMP-3C. Recall that there are no reports of any known function for MpaB’; and we did not identify any obvious functional domains using BLAST or Pfam database tools. However, when using the Phyre2 program (27) to predict and compare potentially conserved three-dimensional structural features with other proteins, we noted a possible similarity in a structural fold with a distant homolog (11%/22% amino acid identity/similarity, *SI Appendix*, Figs. S40-S41)—a *b*-type heme protein latex clearing protein from *Streptomyces* sp. K30 (Lcp_k30_) that was recently biochemically and structurally characterized (28, 29).

Consideration of the proposed catalytic mechanisms from the LcpK30 study (29) guided our speculation that MpaB’ might initiate the oxidative cleavage through proton abstraction by D124 at the C_18_ allylic position. The resultant iron(IV)-oxo species could then react with the epoxide, together with a D124-mediated acid-base catalysis, ultimately leading to the cleavage of the C_19_=C_20_ double bond (*SI Appendix*, Fig. S42) (29). The expected resultant aldehyde was not observed, likely owing to instability; supporting this, chemically synthesized mycophenolic aldehyde (*SI Appendix*, Figs. S8 and S43, and Table S6) was readily oxidized to MPA by *Ao*_M-2-3_ (*SI Appendix*, Fig. S44). Thus, our results overturn the previously proposed direct cleavage of the FDHMP C_15_=C_16_ double bond (*SI Appendix*, Fig. S24) (10–12), which would otherwise lead to DMMPA but not FDHMP-3C as the dominant product when *Ao*_M-2-3_-*mpaA’-mpaB’* was fed DHMP (*SI Appendix*, Fig. S24).

Next, HPLC analysis of the fermentation culture of an aforementioned *Pb*_864_-*ΔmpaH’* strain led to the surprising finding that this *mpaH’* knockout strain retained the ability to produce MPA, albeit with a yield which was approximately 50% lower than that of *Pb*_864_. This mutant strain also produced two novel compounds (MFDHMP-d4 and MFDHMP-d5) with an even shorter isoprenyl chain than MPA (Fig. 1, *SI Appendix*, Figs. S8, S45-S51, and Tables S4 and S9); these correspond to the 5-*O*-methylated products of FDHMP-d4 and FDHMP-d5, presumably stemming from the activity of MpaG’ (Fig. 2*B*). Notably, neither compound was detected in *Pb*_864_ cultures (Fig. 2*B*). The attenuated production of MPA, together with the two over-shortening products by *Pb*_864_-*ΔmpaH’*, suggested the interesting possibility that, while MpaH’ does not catalyze the oxidative cleavage of FDHMP as previously proposed (14), this enzyme apparently does have an MPA-biosynthesis related function, likely somehow involved in the aforementioned *β*-oxidation chain-shortening process. Specifically, MpaH' may function to control the specificity and efficiency of MPA production, perhaps by acting as a “valve” to prevent the excessive *β*-oxidation-mediated shortening of MPA.

To recapitulate the MPA accumulation pattern of *Pb*_864_ in a heterologous host, we investigated the product profile of the *Ao*_M-2-3_-*mpaA’-mpaB’-mpaH’* strain (in which the three genes were co-expressed) when DHMP (20 mg/L) was fed to its induction cultures. As expected, the amount of the penultimate MPA pathway intermediate DMMPA that accumulated in *Ao*_M-2-3_-*mpaA’-mpaB’-mpaH’* was significantly higher than that of *Ao*_M-2-3_-*mpaA’-mpaB’* (Fig. 2*C* and *D*), again emphasizing the importance of MpaH’ for efficient production of either DMMPA or MPA. Notably, whereas we were expecting to only detect the accumulation of DMMPA by *Ao*_M-2-3_-*mpaA’-mpaB’-mpaH’* as the dominant MPA production by *Pb*_864_, we were surprised to observe substantial amounts of FDHMP-d1–d3 as well as low levels of FDHMP-d4, d5; note that the methylated counterparts MFDHMP-d1–d5 were only detected at negligible levels in *Pb*_864_. We speculate that these differences between *Penicillium* and *Aspergillus* species could perhaps be due to their different cellular contexts. Specifically, *Ao*_M-2-3_ may contain a non-specific acyl-CoA hydrolase with broad and highly efficient hydrolytic activities toward the CoA-esters generated from the *β*-oxidation catabolic pathway (*SI Appendix*, Fig. S52). The low-level accumulation of FDHMP-d4–d5 likely resulted from the lower activity of MpaH’ toward DMMPA-CoA than toward MPA-CoA, which could lead to the "leaking" of these two excessively chain-shortened derivatives from peroxisomes (*see below*). Nonetheless, our observation of FDHMP-d1–d5 represented important clues for our following elucidation of these unusual MPA biosynthetic pathway steps.

Interestingly, a PSORT II (30) analysis of the MpaH’ sequence identified a Type 1 peroxisomal targeting sequence like (PTS1-like) GKL tripeptide at its *C*-terminus, which strongly suggested that this protein is localized in peroxisomes—a site where *β*-oxidation metabolism can occur (31, 32). We were able to successfully confirm the peroxisomal localization of MpaH’ via confocal laser scanning microscopy (CLSM) of several *Aspergillus* strains expressing GFP fusion constructs for full-length and GKL-tripeptide-truncated MpaH’ variants alongside the recombinant expression of the peroxisome-specific RFP^SKL^ reporter (*SI Appendix*, Tables S2 and S3). As anticipated, we observed co-localization of the RFP^SKL^ reporter with the GFP-MpaH’^full-length^ but not the GFP-MpaH'^ΔGKL^ fusion proteins (Fig. 3*A*-*D* and *SI Appendix*, Fig. S53). Additionally, feeding experiments demonstrated that the peroxisomal localization of MpaH' increases the efficiency of DMMPA production; specifically, significantly more DMMPA was accumulated in the DHMP-fed *Ao*_M-2-3_-*mpaA’-mpaB’-mpaH’* cultures than in the corresponding *Ao*_M-2-3_-*mpaA’-mpaB’-mpaH’^△GKL^* cultures (Fig. 2*C* and *D*).

**Fig. 3.**
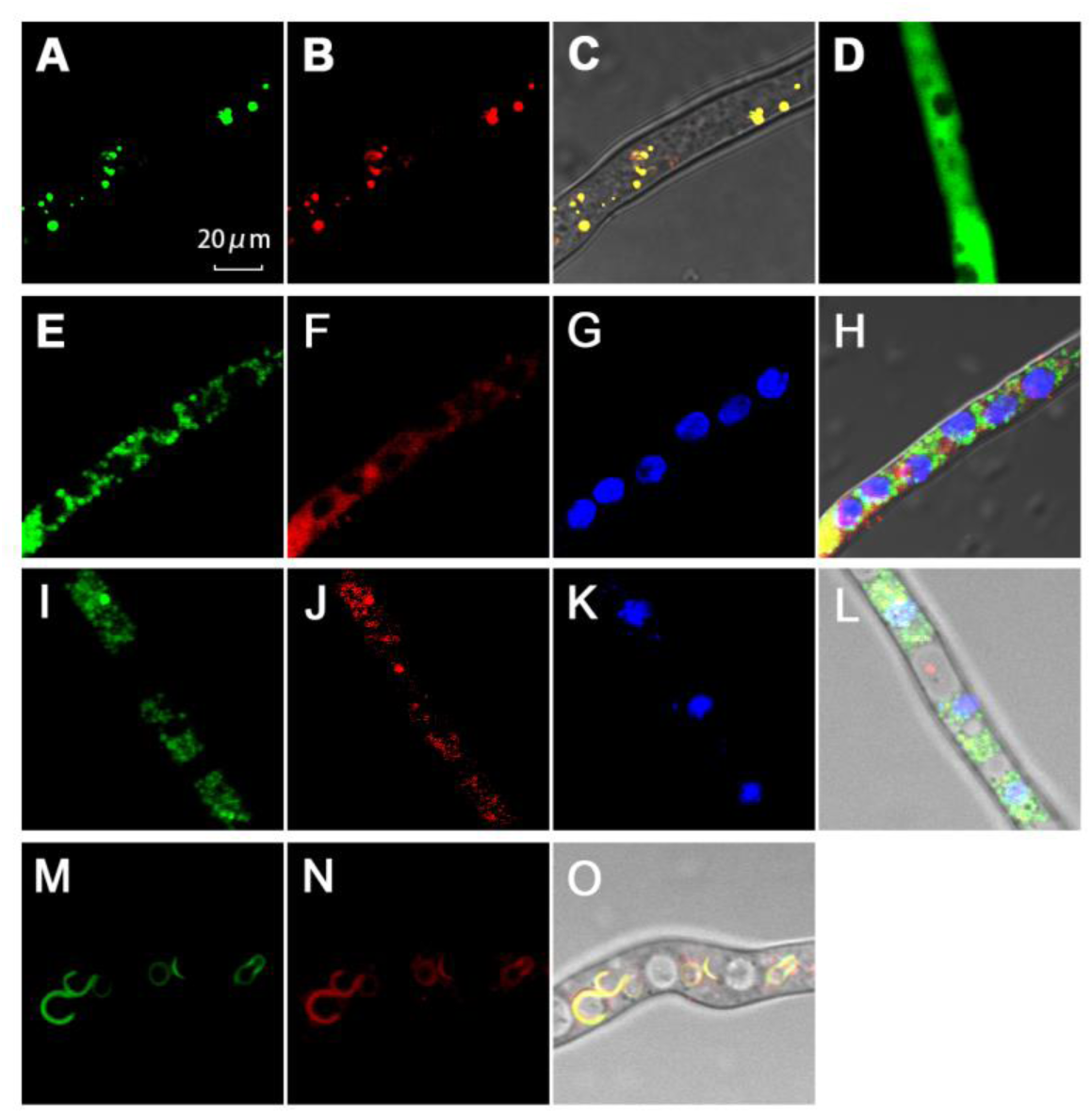
High-resolution confocal images for subcellular localization of MpaH’, MpaB’, MpaDE’, and MpaA’ in *Ao*_M-2-3_. (*A*). The GFP-MpaH’ localization; (*B*). The peroxisomal localization of RFP^SKL^; (*C*). The merged images of *A* and *B* in bright field; (*D*). The GFP-MpaH’^△GKL^ localization; (*E*). The MpaB’-GFP localization; (*F*). The localization of endoplasmic reticulum by “ER-Tracker^TM^ Red”; (*G*). The localization of multiple nuclei by DAPI; (*H*). The merged images of *E*– *G* in bright field; (*I*). The MpaDE’-GFP localization; (*J*). The localization of endoplasmic reticulum by “ER-Tracker^TM^ Red”; (*K*). The localization of multiple nuclei by DAPI; (*L*). The merged images of *I–K* in bright field; (*M*). The GFP-MpaA’ localization; (*N*). The localization of Golgi complex with CellLight™ Golgi-RFP; (*O*). The merged images of *M* and *N* in bright field.

In line with our proposed “valve” function of the peroximal protein MpaH’, the fact that fungal *β*-oxidation of long-chain acyl moieties acids can occur in peroxisomes (31, 32), together with our observation of the suspected *β*-oxidation-derived chain-shortening products in the *Pb*_864_-*ΔmpaH’* and *Ao*_M-2-3_-*mpaA’-mpaB’*/DHMP cultures (Fig. 2), we hypothesized that the *α*/*β*-hydrolase fold containing MpaH’ enzyme may be an acyl-CoA hydrolase that can specifically recognize DMMPA-CoA and/or MPA-CoA. To test this, we heterologously expressed MpaH' in *E. coli* BL21(DE3) cells and purified it to homogeneity (*SI Appendix*, Fig. S54). Indeed, when the purified MpaH’ was incubated with chemically synthesized DMMPA-CoA and MPA-CoA *in vitro*, both DMMPA and MPA were rapidly hydrolyzed from their corresponding CoA-esters (*SI Appendix*, Fig. S55). Analysis using Phyre2 revealed a likely structural relationship between MpaH’ and the peroxisomal hydrolase Lpx1 from *S. cerevisiae* (33), and careful protein sequence analysis (*SI Appendix*, Fig. S56) suggested that MpaH’ is a new member of the type I acyl-CoA thioesterase enzyme family. MpaH’ possesses a well-recognized catalytic triad of S139-D163-H365 (34). To confirm that S139 is a catalytic nucleophile, we mutated this serine into an alanine and, as expected, the hydrolytic activity of the MpaH’^S139A^ mutant for either MPA-CoA or DMMPA-CoA was completely abolished (*SI Appendix*, Fig. S55).

We subsequently analyzed the steady-state kinetics of MpaH’ *in vitro* using the 5,5-dithiobis-(2-nitrobenzoic acid) reagent (35) and found that the *k*_cat_/*K_m_* values of MpaH’ for both DMMPA-CoA (11.6 µM ^-1^ min^-1^) and MPA-CoA (81.5 µM ^-1^ min^-1^) were two orders of magnitude higher than the values for the ten other unnatural CoA-esters that we tested in similar assays (*SI Appendix*, Fig. S57 and Table S10). Thus, our results strongly suggest that MpaH’ is a dedicated MPA-CoA hydrolase with high substrate specificity, and this enzyme apparently exerts a valve-like function to prevent MPA-CoA from further peroxisomal *β*-oxidation and to avoid the hydrolysis of other CoA-esters.

The 6.2-fold higher *k*_cat_/*K_m_* value of MPA-CoA relative to DMMPA-CoA suggests (*SI Appendix*, Table S10) that the 5-*O*-methylation mediated by the methyltransferase MpaG’ likely occurs prior to FDHMP-3C’s entry into peroxisomes. Supporting this, FDHMP-3C was found to be a better substrate for MpaG’ as compared to other potential substrates including DMMPA, FDHMP, 5-MOA, and DHMP (*SI Appendix*, Fig. S58). However, we cannot exclude the possibility that DMMPA methylation could also occur *in vivo* as a minor pathway (Figs. 1 and 4). After the cytosolic methylation of FDHMP-3C, the entry of MFDHMP-3C into peroxisomes could be unidirectional: this entry likely occurs via free diffusion due to its low molecular weight of 388, which is lower than the reported 400 Da cutoff for crossing the single membrane of peroxisome via free diffusion (32). Upon a peroximal CoA ligation reaction—presumably catalyzed by the *β*-oxidation component enzyme CoA ligase—MFDHMP-3C-CoA (with a molecular weight of 1136) would then be restricted to peroxisomes for the following *β*-oxidation pathway steps (*SI Appendix*, Fig. S52).

**Fig. 4.**
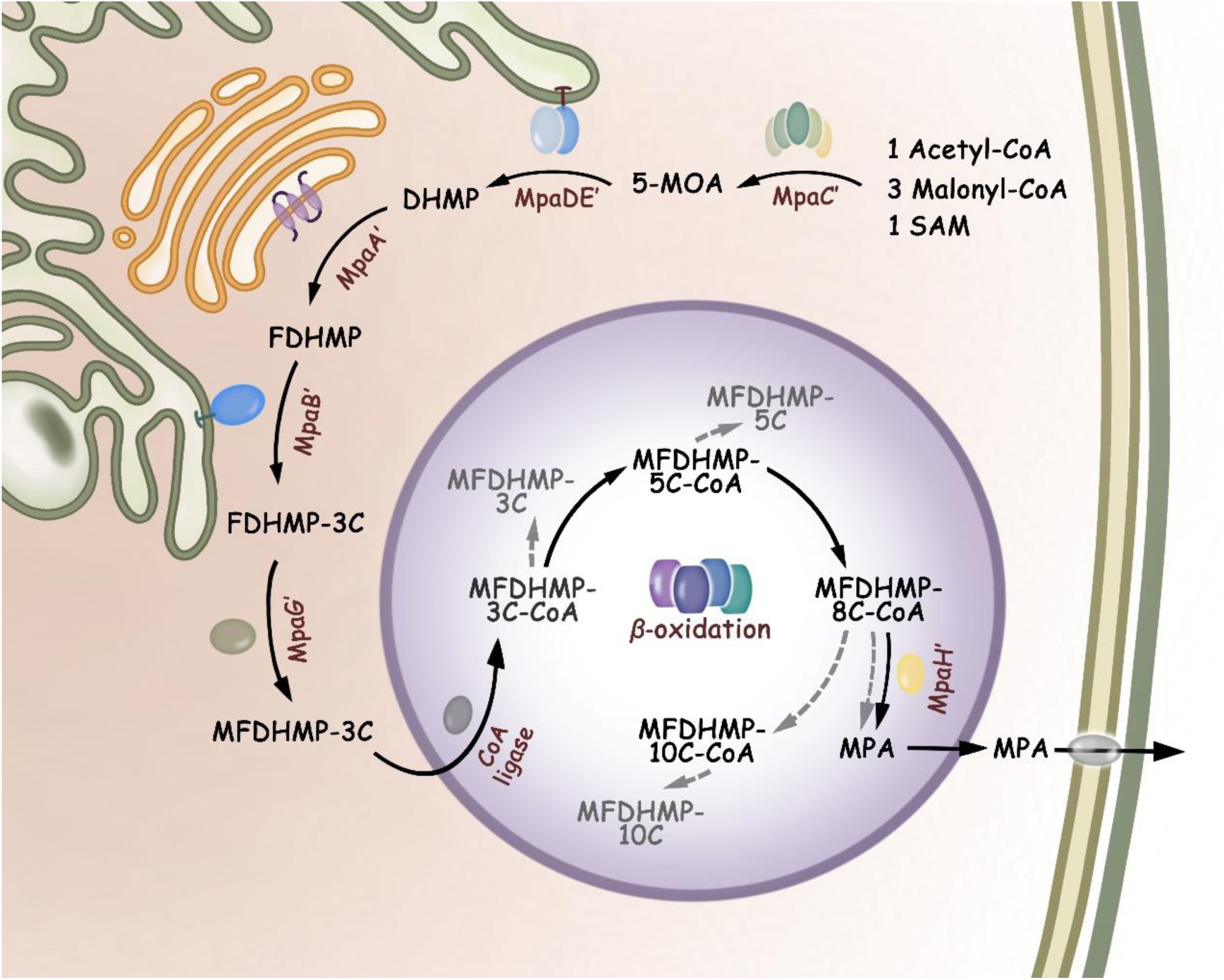
The schematic compartmentalized MPA biosynthesis (Solid arrows: the major pathway; dashed arrows: the shunt pathways), which is sequentially mediated by the cytosolic polyketide synthase MpaC’, the ER-bound P450-hydrolase fusion enzyme MpaDE’, the Golgi apparatus-associated prenyltransferase MpaA’, the ER-bound oxygenase MpaB’, the cytocolic *O*-methyltransferase MpaG’, and the *β*-oxidation machinery and the acyl-CoA hydrolase MpaH’ in peroxisomes.

The importance of the subcellular localization of MpaH’ and the fact that MpaA', MpaB’, and MpaDE’ are predicted to be membrane-associated proteins (*SI Appendix*, Figs. S7 and S59) led us to further investigate the compartmentalization of these MPA biosynthetic enzymes. Specifically, we fused GFP tags to the *N*-or *C*-termini of the transmembrane MpaA’ and the integral monotopic proteins MpaB’ and MpaDE’ (*SI Appendix*, Figs. S3-S4 and Tables S2-S3). Subsequent CLSM observations which revealed the co-localization of the green fluorescence signals of MpaDE’-GFP (or MpaB’-GFP) and the red fluorescence signals of the “ER-Tracker^TM^ Red” marker—outside of DAPI-stained nuclei—together demonstrated that both of these two proteins reside at the endoplasmic reticulum (Fig. 3*E*–*L*). The green fluorescence signals of the GFP-MpaA’ fusion protein was distributed as ring-like structures in hyphal cells that were co-localized with the red fluorescence signals of the CellLight™ Golgi-RFP BacMam 2.0 marker that specifically targets the Golgi complex (Fig. 3*M*–*O*).

The ER-bound nature of MpaDE' is unsurprising, since membrane-anchoring is a common feature of eukaryotic P450 enzymes (35). For MpaA' and MpaB’, their membrane-association is potentially functionally relevant because these enzymes must ostensibly interact with their membrane-embedded substrates including FPP and FDHMP. Finally, it is worth noting that the biotransformation activities of all of the engineered strains carrying the GFP-tagged enzymes did not differ from their non-tagged counterparts, indicating that the fusion fluorescence tags did not alter the catalytic properties of these enzymes.

In this study, we elucidate the previously unknown steps of the full MPA biosynthetic pathway. The insights gained in our work will benefit future efforts for both industrial strain improvement and novel drug development. The intriguing compartmentalization of the MPA biosynthetic enzymes (Fig. 4), including the cytosolic MpaC’ and MpaG’, the inner membrane-associated MpaA’, MpaB’, and MpaDE’, and the peroxisomal acyl-CoA hydrolase MpaH’, work together and thusly enable a unique joining of biosynthetic and *β*-oxidation catabolic machineries. These findings highlight that the underexplored organelle-associated catalytic mechanisms, as for example the final peroxisomal maturation steps of penicillin (36), can enable essential steps in natural product biosynthesis in fungi and other higher organisms. Compared to the better understanding of compartmentalization in biosynthesis of lipids (37) and plant terpenoids (38), the compartmentalized biosynthesis of fungal natural products demands much more attention in the future since only very limited knowledge about the subcellular localization of fungal biosynthetic enzymes and their involvement in product formation and intermediate trafficking has been learned so far. Finally, we suggest that studies of natural product biosynthesis should be liberated from a reductionist emphasis on enzymatic steps and would profit by adopting a more panoramic view of catalytic mechanisms, enzyme subcellular distribution, and global cellular metabolisms.

## Acknowledgments

We thank Prof. Ikuro Abe at the University of Tokyo, Prof. Shuangjiang Liu at Institute of Microbiology, Chinese Academy of Sciences, and Prof. Chunxiang Fu at Qingdao Institute of Bioenergy and Bioprocess Technology, Chinese Academy of Sciences for providing the plasmids pTAex3, pAcGFP1 and pANIC 6D, respectively. We are also grateful to Prof. Jianghua Chen at Xishuangbanna Tropical Botanical Garden, Chinese Academy of Sciences and Prof. Guochang Sun at Zhejiang Academy of Agricultural Sciences for helpful discussion.

## Funding

This work was supported by the National Natural Science Foundation of China (grants 21472204, 81741155 to S.L., 31570030 to W.Z., and 31600036 to F.Q.), the Shandong Provincial Natural Science Foundation (ZR2017ZB0207 to W.Z. and S.L.), the Qilu Youth Scholar Startup Funding of Shandong University (to W.Z.), Chinese Academy of Sciences (grants QYZDB-SSW-SMC042 to S.L., and the Youth Innovation Promotion Association of CAS Grant 2015166 to W.Z.), National Postdoctoral Innovative Talent Support Program (BX20180325 to L.D.), and China Postdoctoral Science Foundation (2016T90650 and 2015M58060 to W.Z and 6188229 to F.L.).

## Author contributions

W.Z., and S.L. conceived and designed this research; W.Z., L.D., Z.Q., X.Z., F.L., Z.L., F.Q., X.W., Y.J., P.M., J.S., S.C., C.G., and F.Q. carried out experiments; W.Z., L.D., F.L., X.W., C.L., and S.L. performed data analysis; W.Z and S.L. wrote the manuscript. All authors contributed to discussion of the research and approved the manuscript.

**Competing interests:** The authors declare no competing interests.

