## Supplementary Information for "Compartmentalized Biosynthesis of Mycophenolic Acid"

**This PDF file includes:**

Supplementary Materials and Methods
Figs. S1 to S59
Tables S1 to S11
References

**Supplementary Materials and Methods**

**General experimental procedures.** High performance liquid chromatography-mass spectrometry (HPLC-MS) analysis was carried out on an Agilent 1290-6430 spectrometer using a Waters symmetry column (4.6 mm × 150 mm, RP18) in water (0.1% formic acid) and acetonitrile (0.1% formic acid) biphasic solvent system. High resolution mass spectra were recorded on a Dionex Ultimate 3000 coupled to a Bruker Nuclear magnetic resonance (NMR) spectra were acquired on Bruker spectrometers (Bruker Avance III 600 and Bruker ASCEND$^{TM}$ 700. NMR data were processed using Topspin software and MestReNova. Reverse phase high performance liquid chromatography (RP-HPLC) was employed to purify MPA derivatives using a Waters X-Bridge 5 μm C18 column with a biphasic solvent system of acetonitrile and water on an Agilent 1260 or Hitachi Primaide system HPLC system. The UV-visible spectra were taken on a Tecan Plate Reader Infinite M200 Pro. The services of primer synthesis and DNA sequencing were provided by TsingKe Biotech (Beijing) and Sangon Biotech (Shanghai). PCR reactions were carried out with Golden Star T6 Super PCR Mix (TsingKe Biotech, Beijing) or 2 × TsingKe Master Mix (TsingKe Biotech, Beijing).

**Total RNA extraction and the preparation of cDNA.** The filter paper dried 7 d mycelia of *Penicillium brevicompactum* NRRL 864 ($Pb_{864}$) was flash frozen with liquid nitrogen and grounded into fine powder. Next, 1 mL TRIZOL solution (Thermo Fisher Scientific, Massachusetts, US) was added into ~100 mg mycelial powder and mixed well. Total RNA was extracted following the manufacturer's instruction (Thermo Fisher Scientific, Massachusetts, US). Briefly, 200 μL chloroform was added to the mycelial powder/TRIZOL mixture and vortex oscillated for 30s. Upon a centrifugation at 12,000 × *g* for 15 min, ~500 μL supernatant was transferred to a new RNase-free tube, mixed with equal volume of isopropyl alcohol, and stored at -80 °C for 30 min. After thawing on ice and a centrifugation at 12,000 × *g* for 15 min, the supernatant was discarded and the pellets were washed with chilled ethanol twice. The tube was left at room temperature for 10 min, and total RNA was re-dissolved with 100 μL RNase-free water. The cDNA library of $Pb_{864}$ was prepared by reverse transcription PCR (RT-PCR) using the purified total RNA as template. RT-PCR was performed with the kit purchased from TaKaRa following the manufacturer's protocol (6110A, TaKaRa Bio Inc., Dalian, China). All the genes from the *mpa'* gene cluster (GenBank accession number: KM595305) were PCR-amplified using the $Pb_{864}$ cDNA as template and the corresponding primer pairs (*SI Appendix*, Table S11). PCR fragments were gel-cleaned using Wizard® Genomic DNA Purification Kit (Promega, Wisconsin, US), double-digested by appropriate endonucleases, and inserted into the restriction enzymes pre-treated pET-28b to afford the expression vectors.

**Construction of *Aspergillus oryzae* M-2-3 expression vectors.** The pTAex3 plasmid was used as the heterologous expression vector in the arginine auxotrophic mutant *Aspergillus oryzae* M-2-3 ($Ao_{\text{M-2-3}}$). The *mpaA', mpaB', mpaDE', mpaF', mpaH'* genes were amplified from cDNA of $Pb_{864}$. The *gfp* (encoding green fluorescence protein, GFP) and *rfp* (encoding red fluorescence protein, RFP) were amplified from pAcGFP1 (a gift from Prof. Shuangjiang Liu at Institute of Microbiology, Chinese Academy of Sciences) and pANIC 6D (a gift from Prof. Chunxiang Fu at Qingdao Institute of Bioenergy and Bioprocess Technology, Chinese Academy of Sciences), respectively, using the corresponding primers (**Table S11**) as follows: *mpaA'*: pTAex3-MpaA'-NdeI-FP and pTAex3-MpaA'-KpnI-RP; *mpaB'*: pTAex3-MpaB'-NdeI-FP and pTAex3-MpaB'-KpnI-RP; *mpaF'*: pTAex3-MpaF'-NdeI-FP and pTAex3-MpaF'-KpnI-RP; *mpaH'*: pTAex3-MpaH'-NdeI-FP and pTAex3-MpaB'-KpnI-RP; *gfp*: pTAex3-GFP-NdeI-FP and pTAex3-GFP-KpnI-RP; $rfp^{SKL}$: pTAex3-RFP-NdeI-FP and pTAex3-RFPskl-KpnI-RP. The linear pTAex3 was prepared by *Nde*I/*Kpn*I double digestion. The PCR fragments of *mpaA', mpaB', mpaF', mpaH', gfp, and* $rfp^{SKL}$ were individually cloned into the linearized pTAex3 with ClonExpress II One Step Cloning Kit (Vazyme Biotech, Nanjing, China) to generate pTAex3-*mpaA'*, pTAex3-*mpaB',* pTAex3-*mpaDE',* pTAex3-*mpaF',* pTAex3-*mpaH'*, pTAex3-*gfp* and pTAex3-$rfp^{SKL}$, respectively. The reverse primer pTAex3-RFPskl-KpnI-RP was designed for introducing PTS1 signal SKL to the *C*-terminus of RFP, leading to pTAex3-$rfp^{\text{SKL}}$. To investigate the function of the PTS1-like signal "GKL" at the *C*-terminus of MpaH', pTAex3-$mpaH'^{\Delta GKL}$ was constructed using the primers pTAex3-MpaH'-NdeI-FP and pTAex3-MpaH'ΔGKL-KpnI-RP with One Step Cloning Kit (Vazyme Biotech, Nanjing, China). To generate the fusion proteins *N*-GFP-MpaH'-*C*, the linear sequence of pTAex3-*mpaH'* was amplified from the plasmid pTAex3-*mpaH'* using the primers nGFPMpaH'-

WL-FP/nGFPMpaH'-WL-RP. The *gfp* was amplified using the primes nGFPMpaH'-FP/nGFPMpaH'-RP, respectively. Then, the *gfp* sequence was ligated to the linear sequence of pTAex3-*mpaH'* to generate the pTAex3-*gfp-mpaH'* using One Step Cloning Kit. Following the same procedures, pTAex3-*gfp-mpaA'*, pTAex3-*gfp-mpaB'*, pTAex3-*mpaA'-gfp*, and pTAex3-*mpaB'-gfp* were constructed for expression of *N*-GFP-MpaA'-*C*, *N*-GFP-MpaB'-*C*, *N*-MpaA'-GFP-*C*, and *N*-MpaB'-GFP-*C*, respectively. All vectors were confirmed by restriction digestion and DNA sequencing (GENEWIZ, Suzhou, China). The vectors for co-expression of multiple genes, including pTAex3-*mpaA'-mpaB',* pTAex3-*mpaA'-mpaB'-mpaH',* pTAex3-*mpaA'-mpaB'-mpaH'*$^{\Delta GKL}$, pTAex3-*mpaA'-mpaH',* and pTAex3-*mpaA'-mpaH'* $^{\Delta GKL}$ were synthetized by Wuxi Qinglan Biotech. Inc. (Wuxi, China).

**Fungal transformation and cultivation.** The plasmids were used to transform $Ao_{\text{M-2-3}}$ by the protoplast–polyethylene glycol method as described previously (1). Briefly, the linear DNA containing the target gene expression cassette was PCR amplified or enzyme digested from an appropriate plasmid. The spores of $Ao_{\text{M-2-3}}$ were collected and inoculated to 50 mL DPY liquid medium (2% dextrin, 1% polypeptone, 0.5% yeast extract, 0.5% $KH_2PO_4$, and 0.05% $MgSO_4$,). After growing at 28 $^{o}$C, 180 rpm for 18 h, mycelia were collected by centrifugation at 8000 ×*g* for 5 min, and then used for preparation of protoplasts with 1% lywallzyme, 1% cellulase, and 0.5% lysozyme in 0.8 M NaCl solution by gentle shaking at 28 $^{o}$C for 1-2 h. Protoplasts were filtered and centrifuged at 2500 ×*g* for 5 min. The recovered protoplasts were diluted to $1 \times 10^8$ mL$^{-1}$ in Solution I (0.8 M NaCl, 10 mM $CaCl_2$, 50 mM Tris-HCl, pH 7.5). Subsequently, ~10 µg (in 10 µL dd $H_2O$) of linear DNA fragment was added into 100 µL each aliquot of protoplasts. The mixtures were incubated on ice for 2 min, 1 mL of Solution II (60% PEG 6000, 0.8 M NaCl, 50 mM $CaCl_2$, 50 mM Tris-HCl, pH 7.5) was then added and further incubated at room temperature for 20 min. CDS soft-top agar (7 mL, 3.5% Czapek-Dox broth, 0.7% agar, 1 M sorbitol) was added to the protoplasts mixtures and poured onto Czapek-Dox plates supplemented with 1 M sorbitol, and incubated at 28 $^{o}$C for 5-7 d. Single colonies of transformants were picked and transferred onto fresh Czapek-Dox plates for another 3 d cultivation. The genotype of each transformant was evaluated by PCR with the specific primers (*SI Appendix*, Table

S11). The confirmed transformant was inoculated into 100 mL CMP medium (Czapek-Dox supplemented with 3% maltose for induction and 1% peptone) to induce gene expression under the $\alpha$-amylase promoter in a 250 mL Erlenmeyer flask for 3 d at 28 $^oC$, 180 rpm. A certain substrate (final concentration 20 mg $L^{-1}$) was fed into the induction culture, which was further cultivated for another 3-5 d.

**Construction of pET28b-*mpaH'* for expression of MpaH' in *E. coli*.** The *mpaH'* encoding sequence was amplified from cDNA of $Pb_{864}$ using the primers pET28b-MpaH'-NdeI-FP/pET28b-MpaH'-NdeI-RP. Then, the fragment was cloned into the *Nde*I-digested pET28b to yield pET28b-*mpaH'*. The vector was confirmed by restriction digestion and DNA sequencing (GENEWIZ, Suzhou, China).

**Site-directed mutagenesis of MpaH'.** The mutation sites were introduced by site-directed PCR with the primers MpaH-S139A-FP/MpaH-S139A-RP (*SI Appendix*, Table S11). Specifically, the complimentary primer pair containing the mutated codon was used for PCR amplification (pET28b-*mpaH'* as template, 35 cycles of 95 ℃ for 10 s, 55 ℃ for 15 s, and 72 ℃ for 2 min; FailSafe PCR system, Epicentre, Wisconsin, US) to yield a linear plasmid containing the mutated sites. After gel purification with Wizard® Genomic DNA Purification Kit, the circulation of the linear plasmid DNA was performed using homologous recombinase (Hieff Clone™ One Step Cloning Kit, Yeasen, Shanghai, China) by following the vendor's manual, and then chemically transformed to *E. coli* BL21(DE3). The introduction of the mutated codon was confirmed by DNA sequencing (GENEWIZ, Suzhou, China).

**Protein expression and purification.** A single colony of a certain *E. coli* BL21 (DE3) transformant that carries a specific expression vector was inoculated into 10 mL LB medium containing 50 mg $L^{-1}$ kanamycin and shaking incubated at 37 $^oC$ overnight. The resultant seed culture (5 mL) was used to inoculate 500 mL LB medium containing 50 mg $L^{-1}$ kanamycin. The cells were grown at 37 $^oC$ for 2–3 h until $OD_{600}$ reached 0.4–0.6. Then, isopropyl-D-thiogalactopyranoside (IPTG) was added to a final concentration of 0.1-0.2 mM to induce gene expression at 18 $^oC$, 150 rpm for 16 h. The culture was centrifuged at

6,000 ×*g*, 4 °C for 10 min at 4 °C to collect cells. The freeze–thaw cell pellets were re-suspended with 50 mL lysis buffer (50 mM $NaH_2PO_4$, 300 mM NaCl, 10 mM imidazole, 10% glycerol, pH 8.0) and applied to sonication. Cell debris was removed by centrifugation at 10,000 ×*g*, 4 °C for 60 min. The supernatant was mixed with 1 mL Ni-NTA agarose (Qiagen, Venlo, Netherlands) for 1 h at 4 °C. The slurry was loaded onto an empty column, and the column was washed stepwise with 100 mL lysis buffer and 400–600 mL wash buffer (50 mM $NaH_2PO_4$, 300 mM NaCl, 20 mM imidazole, 10% glycerol, pH 8.0). The bound $His_6$-tagged proteins were eluted with 5–10 mL elution buffer (50 mM $NaH_2PO_4$, 300 mM NaCl, 250 mM imidazole, 10% glycerol, pH 8.0). The proteins were further purified and concentrated with 30 kDa size-exclusion filters (Amicon, Millipore Corporation, Bedford, MA, US). The final desalting step was attained by buffer exchange into storage buffer (50 mM $NaH_2PO_4$, 10% glycerol, pH 7.4) with a PD-10 column (GE Healthcare, Illinois, US). The concentration of proteins were determined by Bradford assay (2) using bovine serum albumin (BSA) as standard.

***In vitro* enzymatic assay of MpaG'.** Following the previously established protocol (3), the standard assay containing 1 μM MpaG', 0.5 mM substrate, and 5.0 mM SAM in 100 μL reaction buffer (50 mM $NaH_2PO_4$, pH 8.0, 10% glycerol) was carried out at 28 °C for 1h and quenched with equal volume of ethyl acetate. The two-time organic extracts were combined and dried by $N_2$ flow, and re-dissolved in 100 μL methanol for HPLC and LC-MS analysis.

***In vitro* enzymatic assay of MpaH'.** The standard assay containing 10 nM MpaH', 1 mM substrate in 100 μL reaction buffer (50 mM $NaH_2PO_4$, pH 8.0, 10% glycerol) was performed at 28 ℃ for 20 min and quenched with equal volume of ethyl acetate. The two-time organic extracts were combined and dried by $N_2$ flow, and re-dissolved in 100 μL methanol for HPLC and LC-MS analysis.

**Steady-state kinetic analysis of MpaH'.** The kinetic analysis of the acyl-CoA hydrolase MpaH' was performed by following a previous procedure (4) with minor modifications. Briefly, the free thiol group released from hydrolysis of CoA esters reacts with the reagent

5,5-dithiobis (2-nitrobenzoic acid) (DTNB) to produce a yellow product, 5-thio-2-nitrobenzoate, which can be monitored spectrophotometrically in solution at 412 nm. The MpaH' catalyzed reactions were performed in a total volume of 100 μL buffer that contains 50 mM $NaH_2PO_4$ (pH 8.0), 1% glycerol, 5 nM to 1 μM MpaH', 0.2 mM DTNB, and 0.1 to 2 mM of acyl-CoA ester in a 96-well plate. The reactions were monitored on an Infinite M200 Pro (Tecan) at 30 ℃ over a period of 30 min. The calculated velocities under different substrate concentrations were plotted and fit into Michaelis-Menton equation for the values of $k_{cat}$ and $K_m$.

**In-frame deletion of *mpaB'* and *mpaH'* in *P. brevicompactum* NRRL 864.** The split-marker recombination strategy (5) was employed to delete the gene of *mpaB'* and *mpaH'* from $Pb_{864}$. Briefly (*SI Appendix*, Fig. S12), the plasmid pRSFD-*hygB* was first constructed by inserting the hygromycin B resistance gene into the pRSFDuet backbone between *Bgl*II and *Hind*III restriction sites by ligation. Two DNA fragments corresponding to ~1.4 kb and ~1.3 kb of the 5' and 3' flanking sequences of *mpaB'* were amplified from $Pb_{864}$ genomic DNA and inserted into the *Sal*I/*Hin*dIII and *Kpn*I/*Xho*I sites of pRSFD-*hygB* vector sequentially, giving rise to pRSFD-*mpaB'up*-*hygB*-*mpaB'down*. Next, two DNA fragments were amplified from pRSFD-*mpaB'up*-*hygB*-*mpaB'down* using the primers QF121F/QF126R and QF126F/QF122R. The PCR products were purified using the Cycle-pure Kit (Omega Bio-tek, Inc., Georgia, US), and then used to transform the protoplasts of $Pb_{864}$ as previously described (6). Transformants were selected on minimal medium supplemented with 1 M sorbitol, 2% glucose, and 300 μg $mL^{-1}$ hygromycin. Anchored PCR analysis with the primers QF127F/QF128R, QF128F/QF127R, and QF129F/QF129R was performed to verify the correct integration events in $Pb_{864}$-*ΔmpaB'*. Similar experiments were performed to construct the mutant $Pb_{864}$-*ΔmpaH'*.

**Sequence alignment analysis.** Sequence alignments were performed on MAFFT 7.0 (8), and then the aligned regions with gaps or poorly aligned sites were trimmed using the TrimAl tool (9) with strict method in Phylomen 2.0 (10).

**Simulation of three-dimensional structures.** The three-dimensional structures of MpaB’ and MpaH’ were modeled using SWISS-MODEL (11) and Phyre2 (12). Figs. S41 and S56 that show protein structures were prepared using PYMOL (http://www.pymol.org).

**Confocal microscopy.** The transformant bearing GFP or RFP fusion constructs was grown for 3-5 d at 28℃ in CMP medium (Czapek-Dox supplemented with 3% maltose for induction and 1% peptone, 100 mL) to induce protein expression under the $\alpha$-amylase promoter in a 250 mL Erlenmeyer flask. The fresh mycelia were transferred to the staining regents after washing with sterilized water for three times. Specifically, 200 μL CellLight™ Golgi-RFP was added to the washed mycelia and treated at 4 $^{o}$C for 30 min, to which 1 mL sterilized water was added and the mixture was incubated at 37 $^{o}$C for another 30 min. For ER-Tracker$^{TM}$ Red staining, 200 μL regent was added to the washed mycelia and treated at 37 $^{o}$C for 45 min before washing with 1 mL sterilized water twice; after ER-Tracker$^{TM}$ Red staining, the mycelia were transferred to 150 μL DAPI solution (800 μg mL$^{-1}$) for another 2 min and washed with 1 mL sterilized water twice. Confocal laser scanning fluorescence images of fungal structures were recorded on an Olympus FluoView$^{TM}$ FV1000 laser scanning microscope (Olympus America). A krypton-argon laser was used as the source of excitation at 488 nm. The green fluorescence signals, the red fluorescence signals, and the DAPI fluorescence signals were recorded at 505 nm, 559 nm and 619 nm respectively. The images were processed with Olympus FluoViewVer. 4.0b.

**Chemical Synthesis**

**Mycophenolic aldehyde** (13-15)

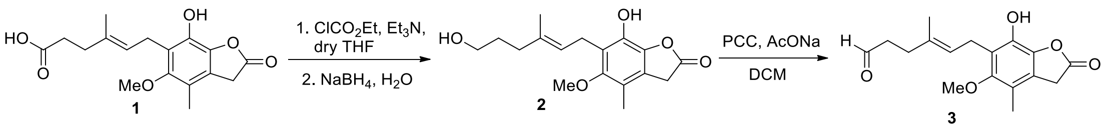


A solution of $ClCO_2Et$ (55 μL) in THF (1 mL) was added to the solution of mycophenolic acid (**1**, 96 mg) and $Et_2N$ (85 μL) in THF (10 mL) at -5 $^{o}$C and the mixture was stirred for 0.5 h. $Et_3NH^+C1^-$ was removed by filtration and the filtrate was added over 0.5 h to $NaBH_4$ (55 mg) in $H_2O$ (0.5 mL) at 10-15 $^{o}$C. The mixture was stirred at room temperature for 18 h, acidified with 3 *N* HCl and extracted with ethyl acetate. The organic extract was washed with water, dried with anhydrous $MgSO_4$, and evaporated, and the residue was heated in 3 *N* NaOH (5 mL) at 90 °C for 0.5 h. After acidification with 3 *N* HCl the reaction mixture was extracted with ethyl acetate, which was washed with aq. $NaHCO_3$ and $H_2O$ and dried with anhydrous $MgSO_4$. Evaporation of the organic phase and crystalline of the residue from ethyl acetate-petrol ether gave **2** (46 mg). Pyridimum chlorochromate (PCC, 97 mg) and sodium acetate (12 mg) were suspended in 5 mL of anhydrous $CH_2Cl_2$ and compound **2** (46 mg) in 5 mL of $CH_2C1_2$ was added in one portion to the magnetically stirred solution. After 2 h reaction at room temperature the brown precipitate was removed by filtration and the filtrate was evaporated to give rise to crude residue, which was further run column chromatography on silica gel to give mycophenolic aldehyde (**3**) (21.3 mg).

**Demethylmycophenolic acid** (**DMMPA**) (3)

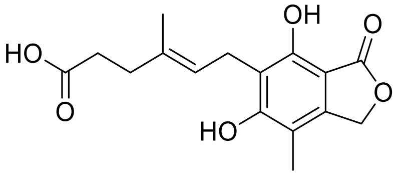


A suspension of MPA (192 mg, 0.6 mmol) and LiI (ultra-dry, 320 mg, 2.4 mmol) in 2,4,6-collidine (6 mL) was heated to 140 °C under $N_2$ for 6 h, the yellow suspension was cooled to room temperature. A clear solution was obtained by slow addition of HCl (3 *N*, 30 mL). The mixture was extracted with 20 mL ethyl acetate by 3 times. The organic phase was dried over anhydrous $Na_2SO_4$, and concentrated to give a yellow solid, which was purified by flash column chromatography (Ethyl acetate: Petroleum ether = 1:2) to afford a white solid (104 mg).

**MPA-CoA**

MPA (160 mg) was added to acetonitrile (3 mL) following by the addition of carbonyldiimidazole (120 mg) with stirring. The mixture was further stirred for 30 min at room temperature under $N_2$ and followed by dropwise addition of CoA solution (320 mg in 1 mL $H_2O$) and adjusted pH to 8 with saturated $NaHCO_3$ solution. The mixture was stirred for 6 h at room temperature, and then concentrated and purified by preparative HPLC to afford a white solid powder (94 mg).

**DMMPA-CoA**

To a suspension of DMMPA (70 mg, 0.23 mmol) in acetonitrile (2 mL), CDI (55 mg, 0.34 mmol) was added with stirring. The resulted clear mixture was stirred for 30 min under $N_2$. A solution of CoA (175 mg, 0.23 mmol) in water (0.5 mL) was added dropwise. The pH of the solution was adjusted to 8 by addition of saturated $NaHCO_3$. The mixture was stirred for 6 h at room temperature and followed by concentration and purification with preparative HPLC to give rise to a white solid powder (46 mg).

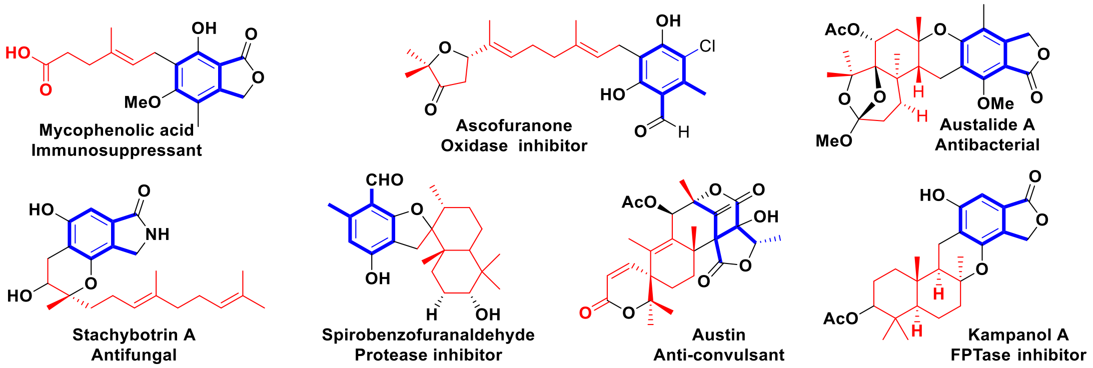


**Fig. S1.** The selected fungal tetraketide–terpenoid (TKTP) family members with various biological activities.

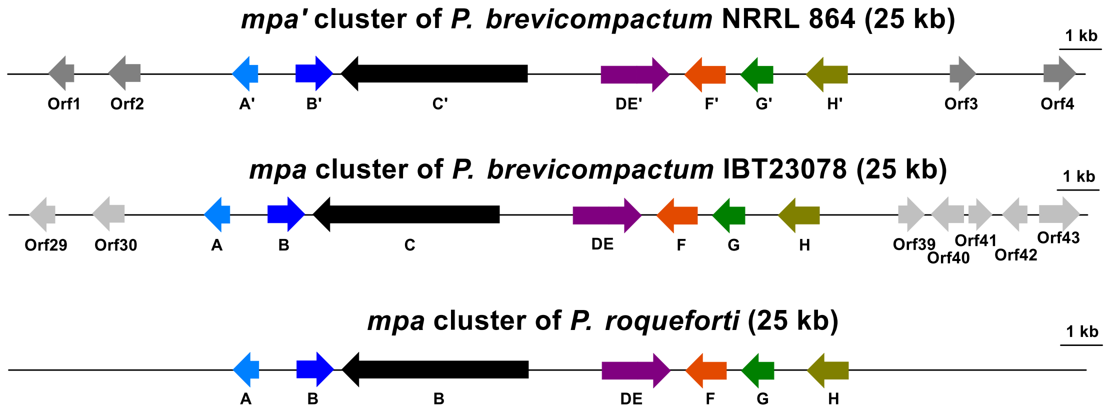


**Fig. S2.** Gene organization of the three reported MPA biosynthetic gene clusters (16-18).

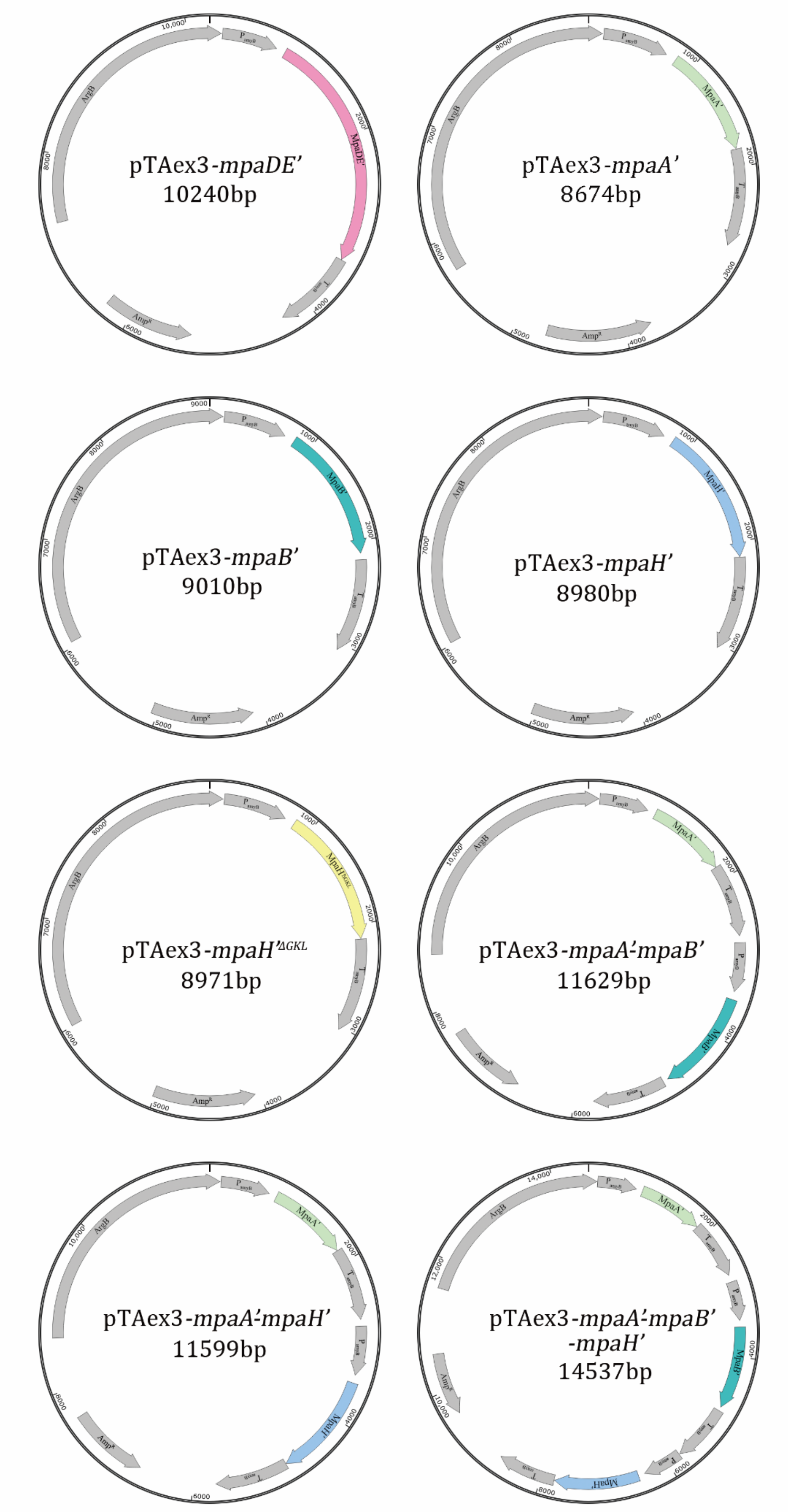
pTAex3-mpaDE’
10240bp
pTAex3-mpaA’
8674bp
pTAex3-mpaB’
9010bp
pTAex3-mpaH’
8980bp
pTAex3-mpaH’ΔGKL
8971bp
pTAex3-mpaA’-mpaB’
11629bp
pTAex3-mpaA’-mpaH’
11599bp
pTAex3-mpaA’-mpaB’
-mpaH’
14537bp

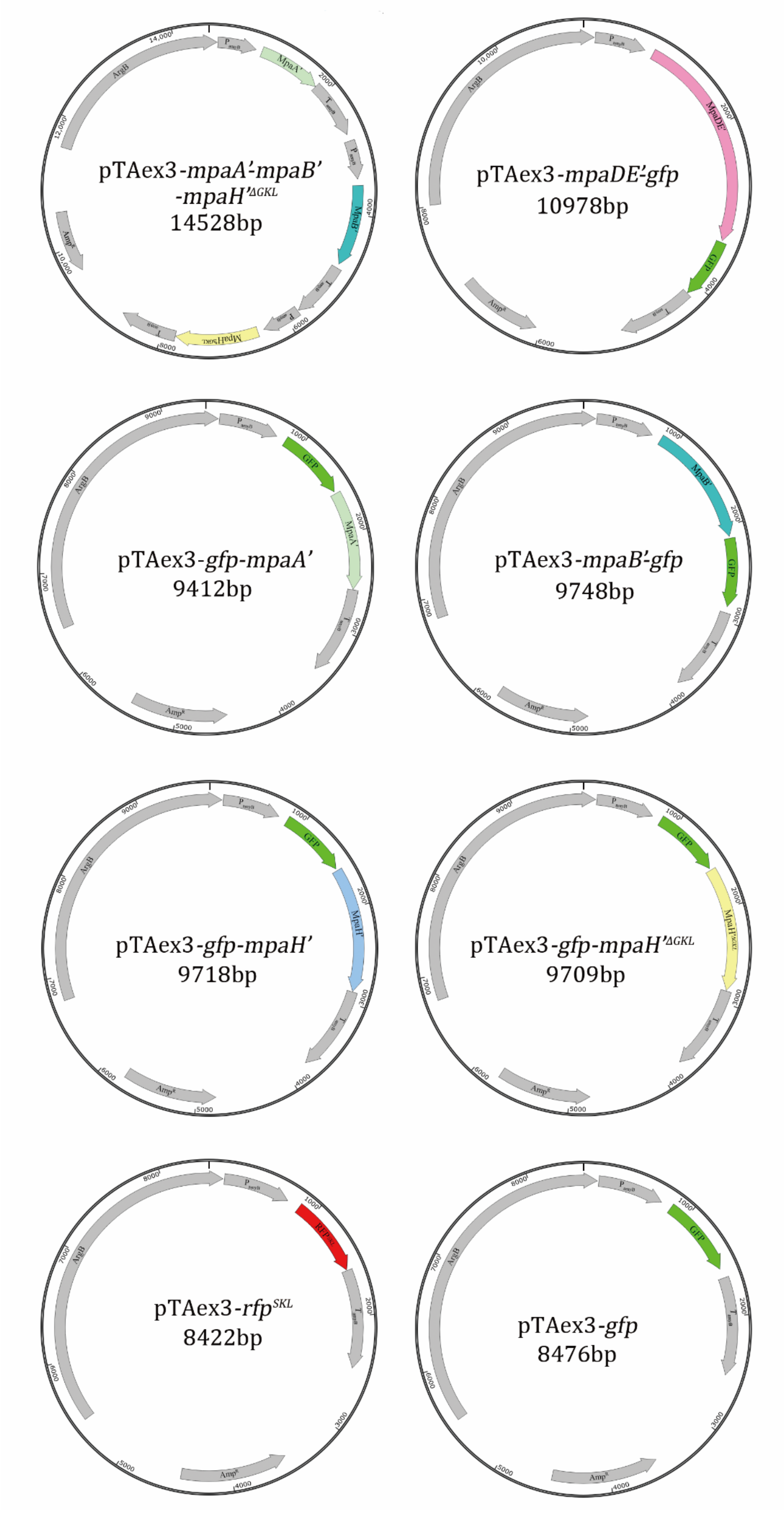
pTAex3-mpaA'-mpaB'
-mpaH'ΔGKL
14528bp
pTAex3-mpaDE'-gfp
10978bp
pTAex3-gfp-mpaA'
9412bp
pTAex3-mpaB'-gfp
9748bp
pTAex3-gfp-mpaH'
9718bp
pTAex3-gfp-mpaH'ΔGKL
9709bp
pTAex3-rfpSKL
8422bp
pTAex3-gfp
8476bp
ArgB
MpaA'
MpaB'
MpaDE'
GFP
RFP
AmpR

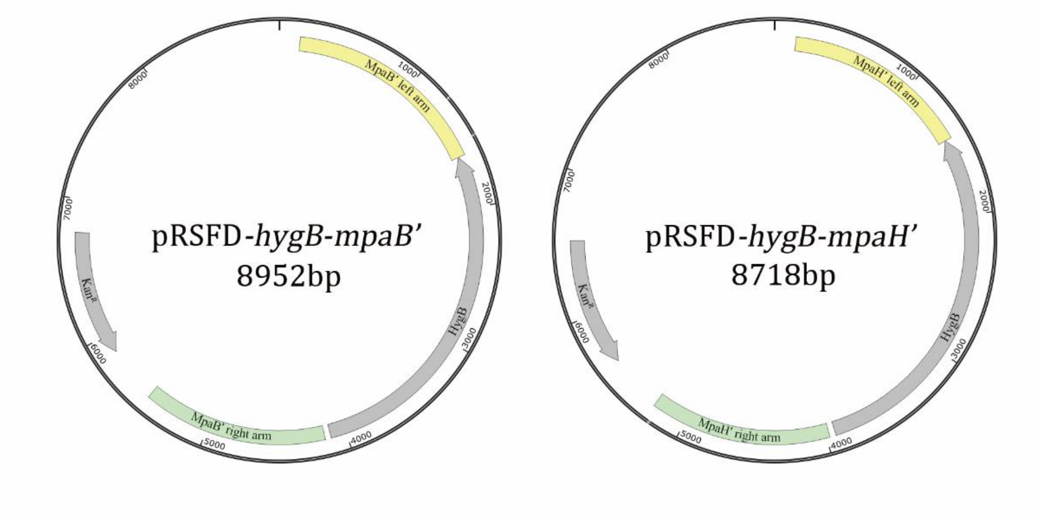


**Fig. S3.** The plasmid maps for the vectors constructed in this study.

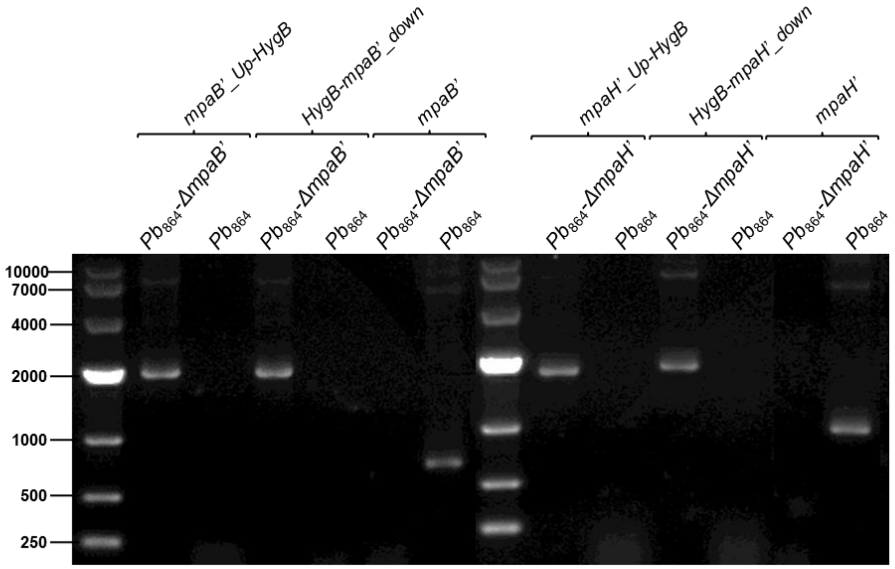
mpaB'_Up-HygB
HygB-mpaB'_down
mpaB'
mpaH'_Up-HygB
HygB-mpaH'_down
mpaH'
$Pb_{864}$-ΔmpaB'
$Pb_{864}$
$Pb_{864}$-ΔmpaH'
10000
7000
4000
2000
1000
500
250

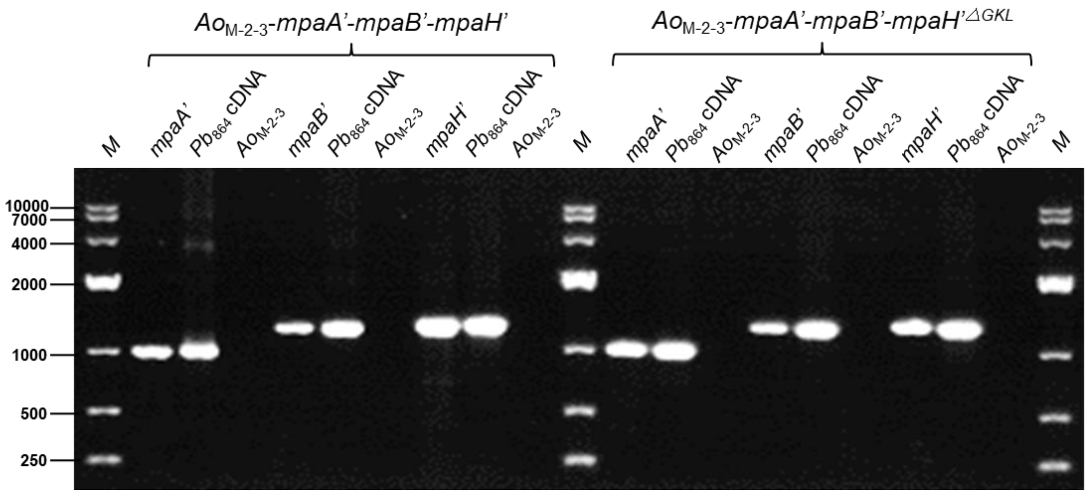
$Ao_{M-2-3}$-mpaA'-mpaB'-mpaH'
$Ao_{M-2-3}$-mpaA'-mpaB'-mpaH'$^{\triangle GKL}$
M
mpaA'
$Pb_{864}$ cDNA
$Ao_{M-2-3}$
mpaB'
mpaH'
10000
7000
4000
2000
1000
500
250

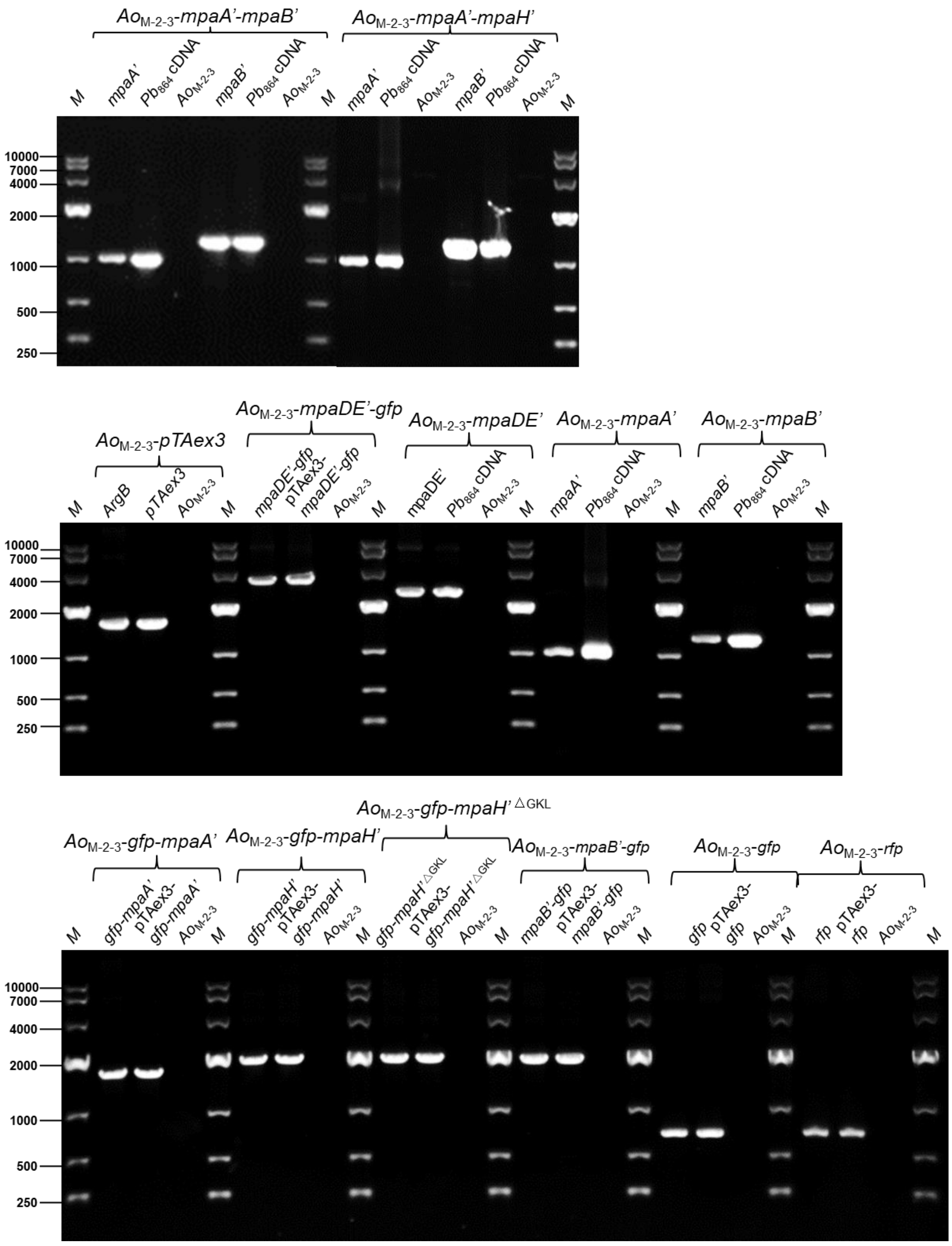


**Fig. S4.** PCR confirmation results of different *A. oryzae* M-2-3 transformants and the gene knockout mutants of *Pb*864.

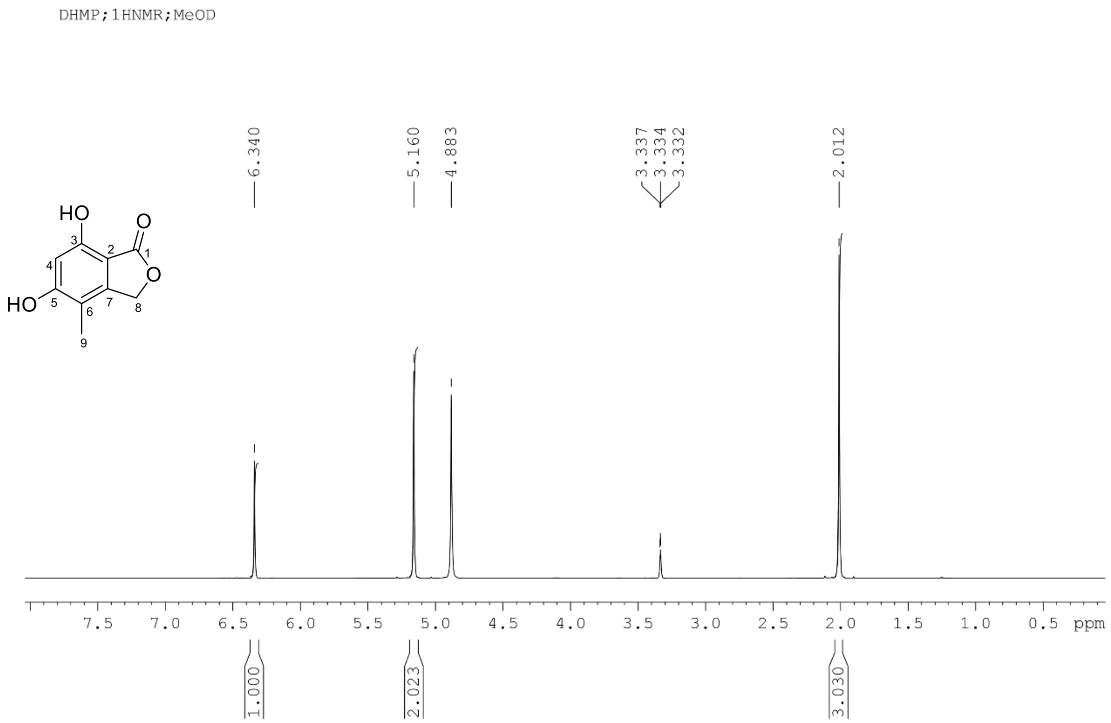


**Fig. S5.** $^{1}$H NMR spectrum of DHMP in MeOD.

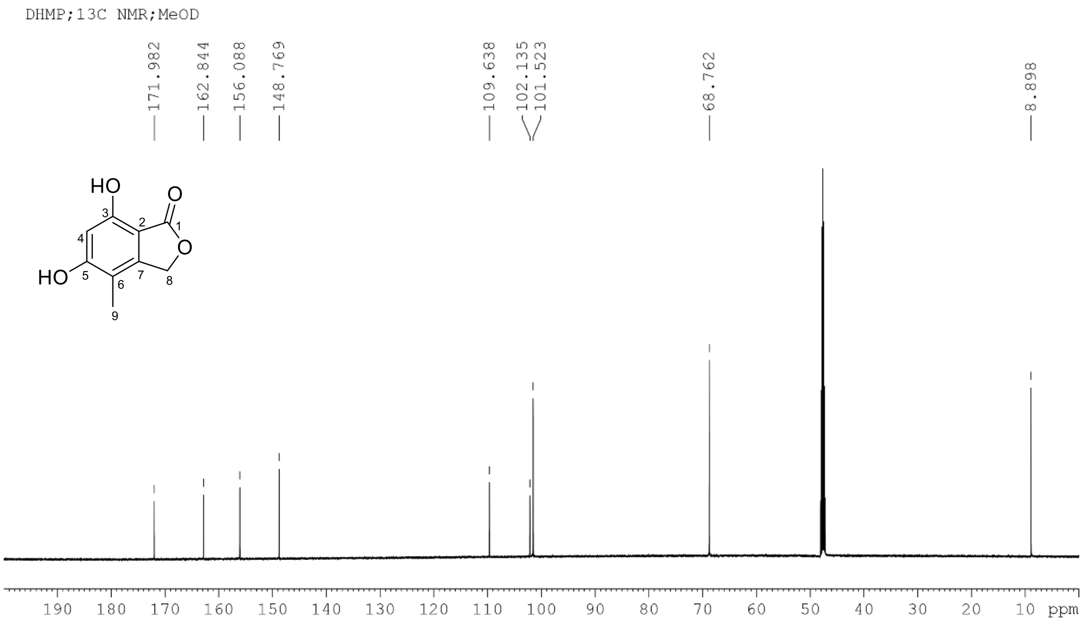


**Fig. S6.** $^{13}C$ NMR spectrum of DHMP in MeOD.

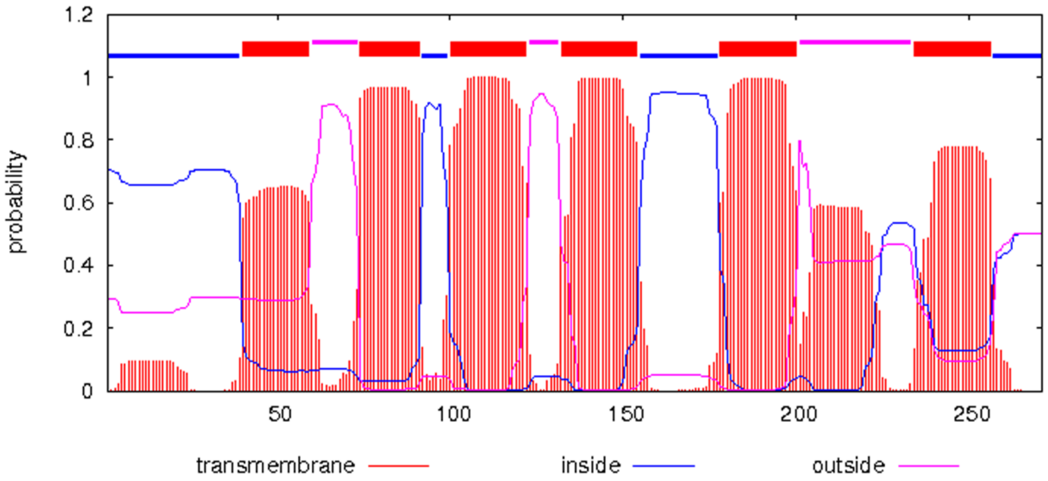


**Fig. S7.** Prediction of transmembrane regions in MpaA' by TMHMM Server v. 2.0 (http://www.cbs.dtu.dk/services/TMHMM/).

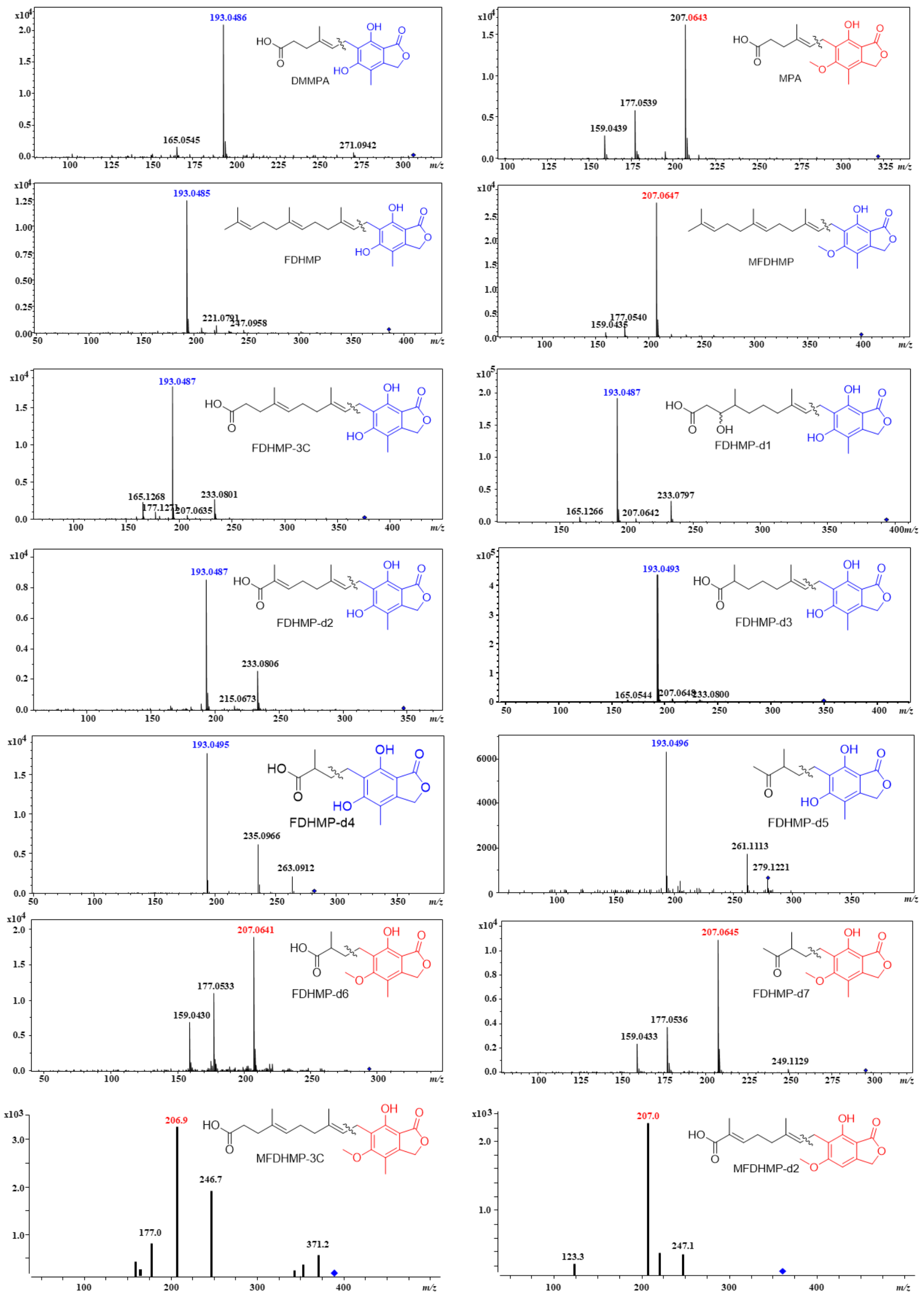


**Fig. S8.** MS/MS analysis of MPA derivatives. The molecular ions are marked by blue diamonds. The secondary fragment ion peaks and the corresponding moieties of 193 are colored in blue, while those of 207 are colored in red.

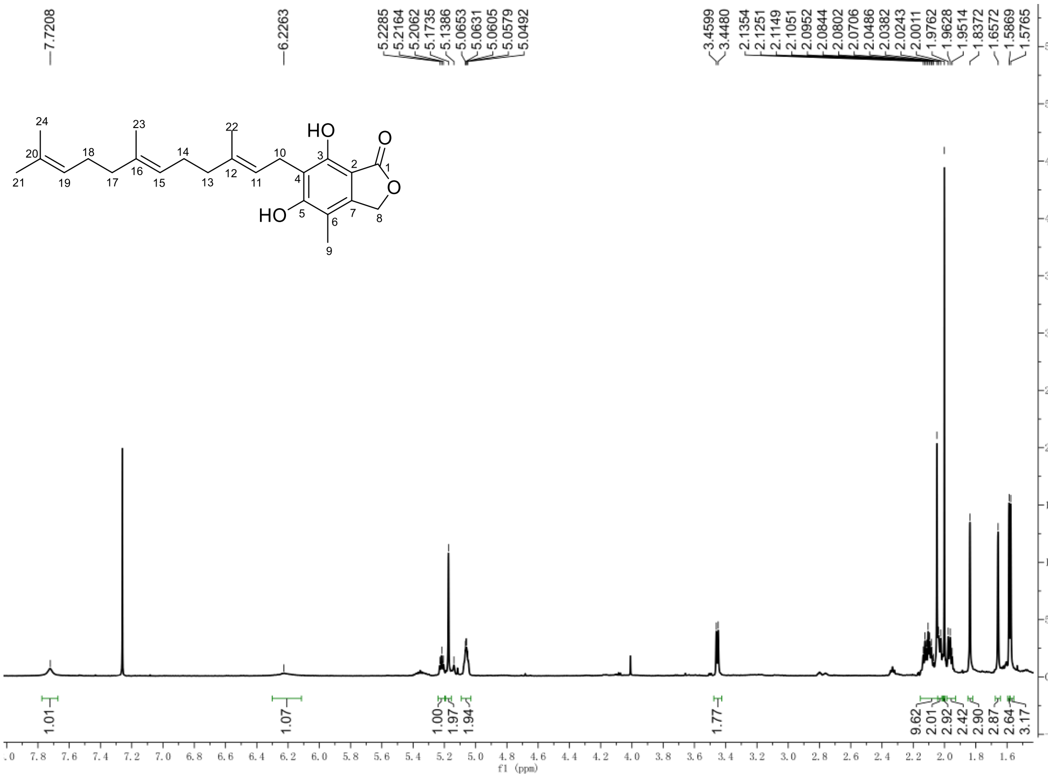


**Fig. S9.** $^{1}$H NMR spectrum of FDHMP in $CDCl_3$.

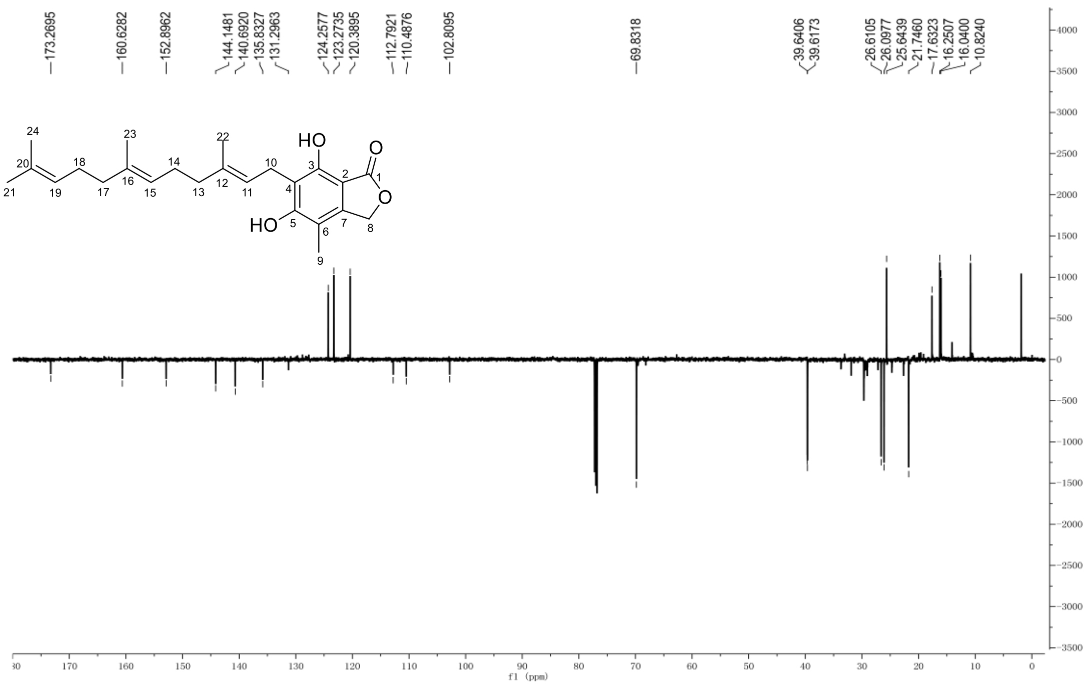


**Fig. S10.** $^{13}$C NMR spectrum of FDHMP in $CDCl_3$.

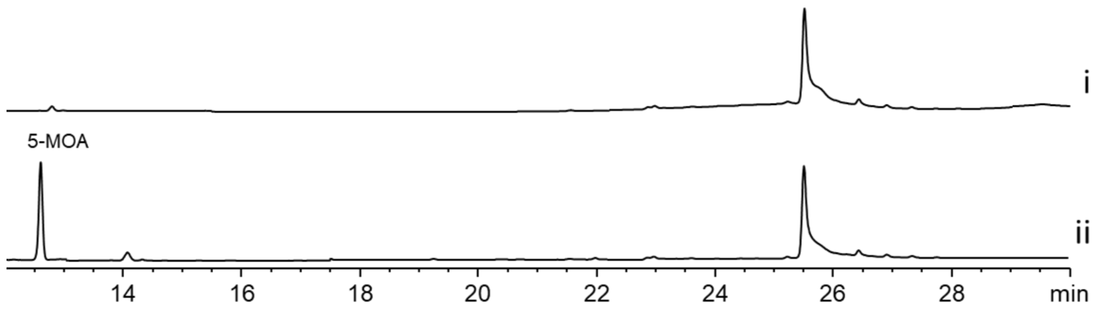


**Fig. S11.** HPLC analysis (254 nm) of *Ao*$_{M\text{-}2\text{-}3}$-*mpaA'* with the feeding of 5-MOA. i, the intracellular extract of *Ao*$_{M\text{-}2\text{-}3}$-*mpaA'*/5-MOA; ii, the extracellular extract of *Ao*$_{M\text{-}2\text{-}3}$-*mpaA'*/5-MOA.

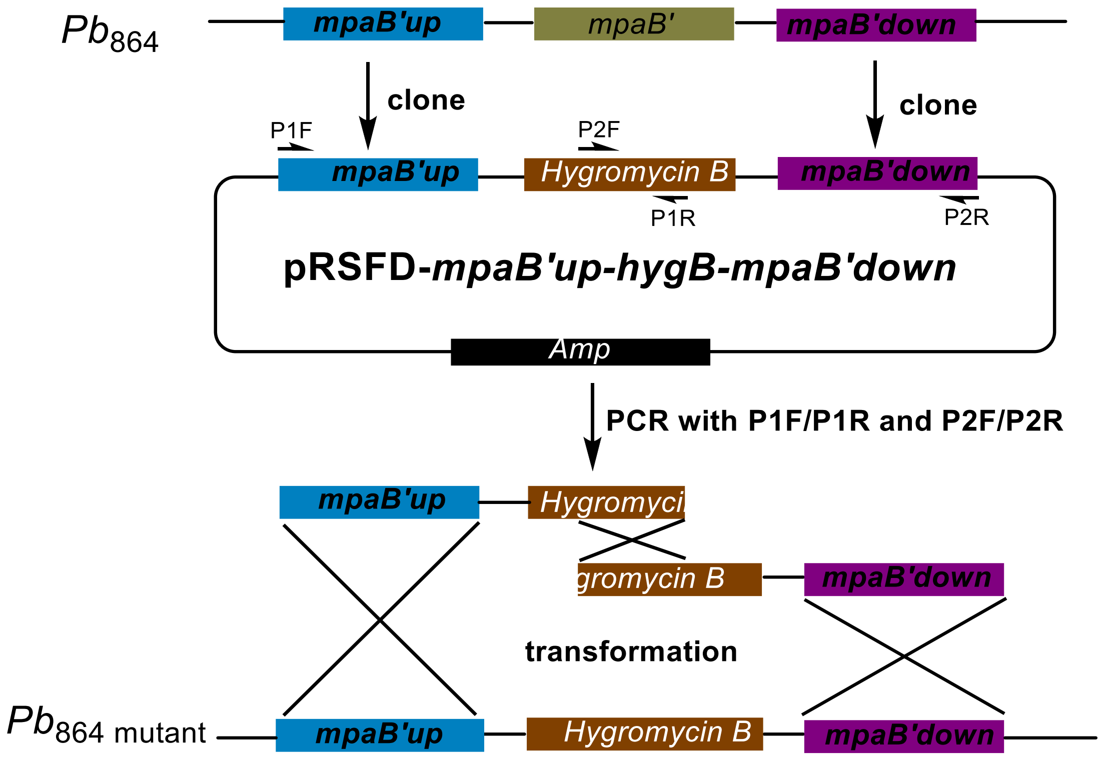


**Fig. S12.** The illustration of split-marker recombination strategy for *mpaB'* knockout in *Pb*864.

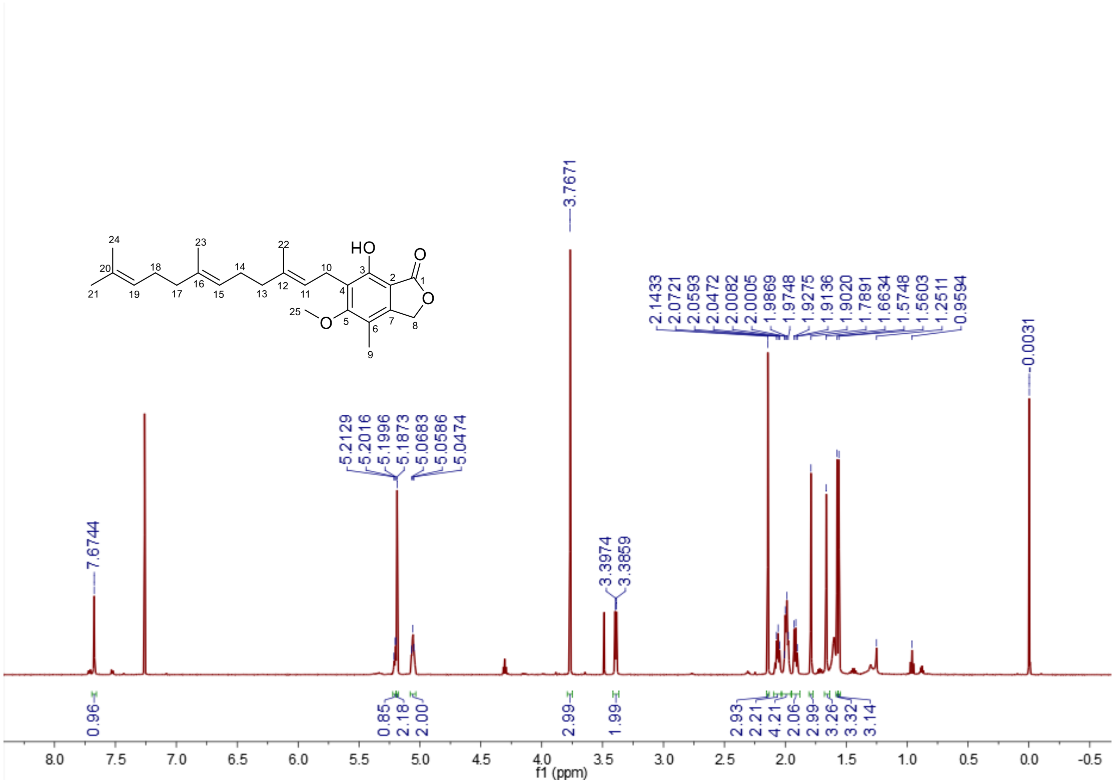

**Fig. S13.** $^{1}$H NMR spectrum of MFDHMP in $CDCl_3$ (600 MHz).

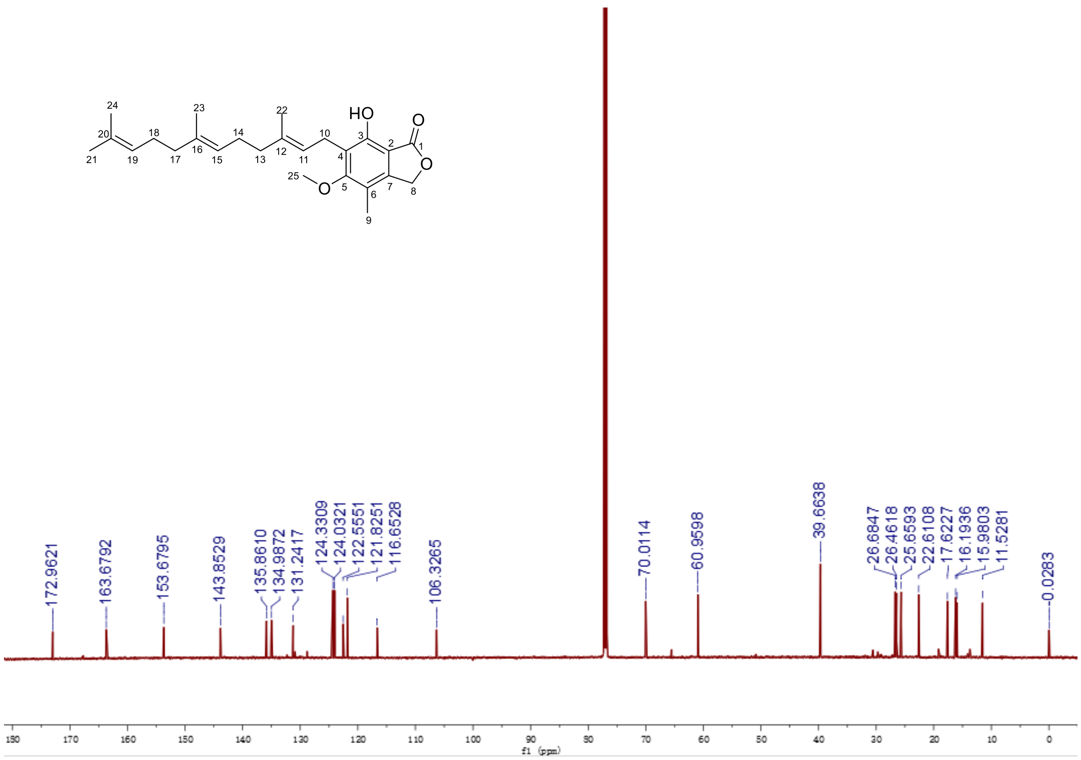


**Fig. S14.** $^{13}$C NMR spectrum of MFDHMP in $CDCl_3$ (600 MHz).

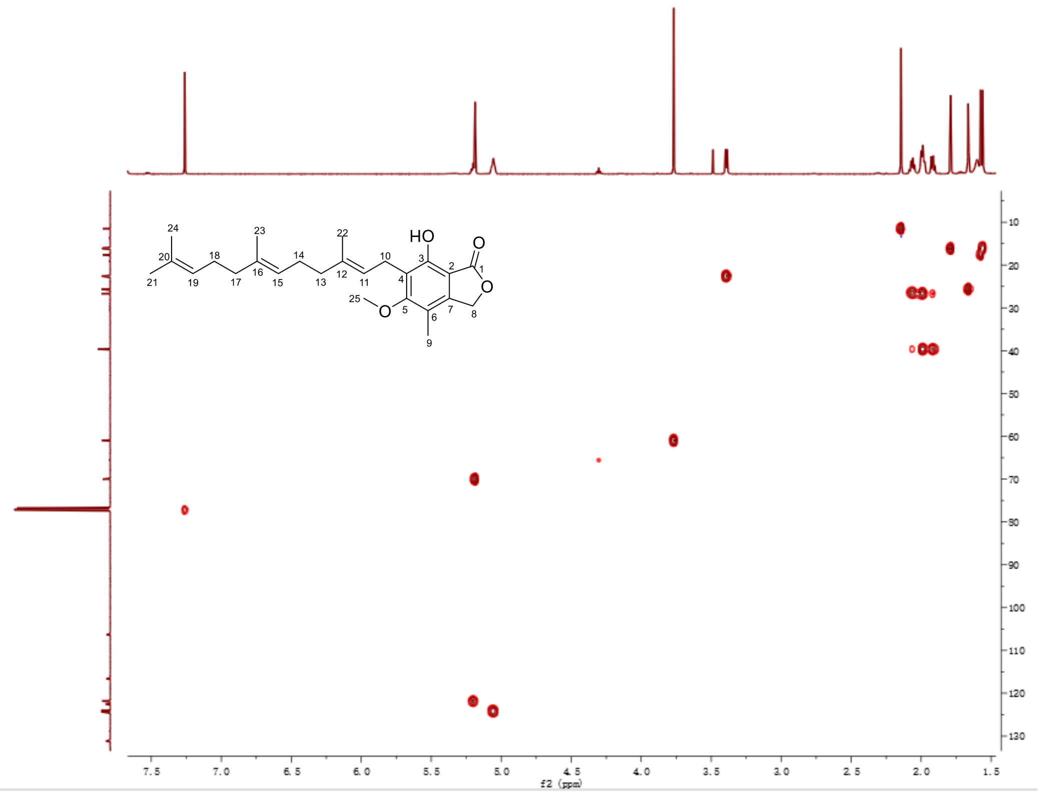


**Fig. S15.** HSQC spectrum of MFDHMP in $CDCl_3$.

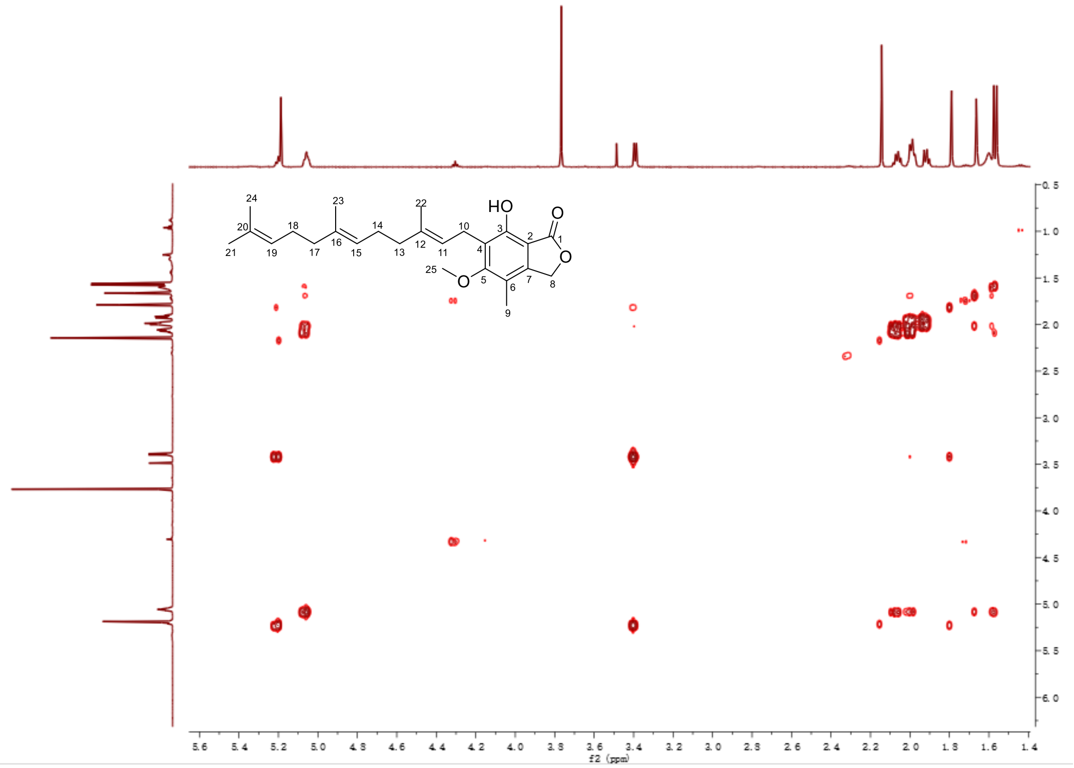


**Fig. S16.** $^{1}H$-$^{1}H$ COSY spectrum of MFDHMP in $CDCl_3$.

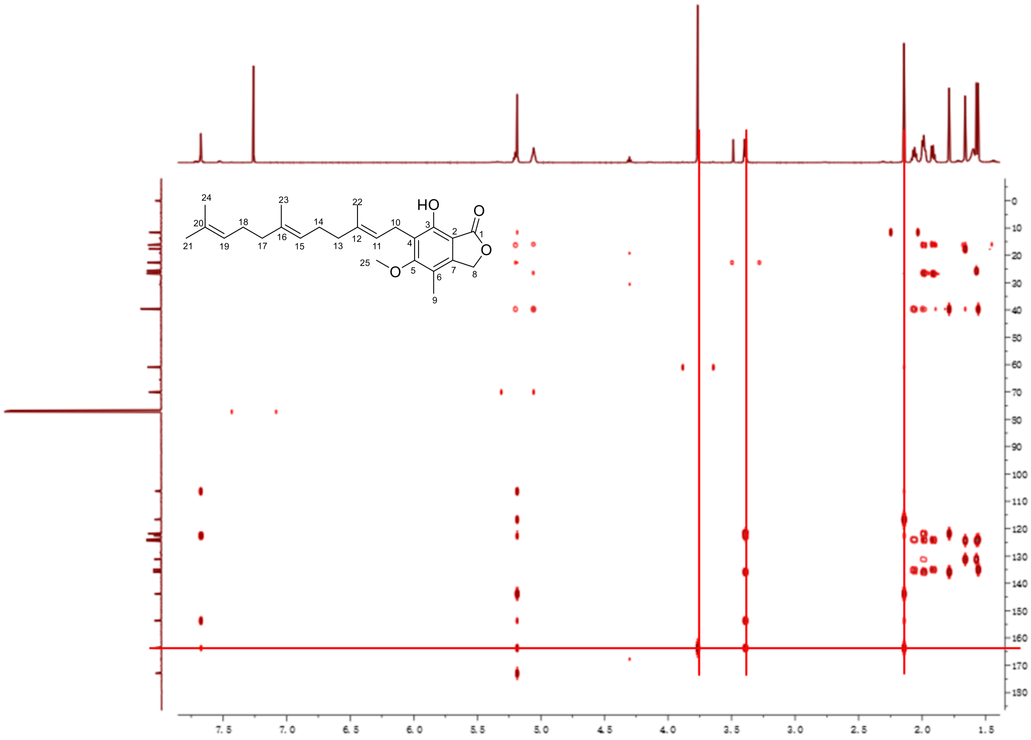

**Fig. S17.** HMBC spectrum of MFDHMP in $CDCl_3$. The crossed red lines indicate that the assignment of the methoxyl group at C5 is based on the HMBC correlations from MeO-5 ($\delta$H 3.77, *s*, 3H), $CH_2$-10 ($\delta$H 3.39, *d*, 2H), and Me-9 ($\delta$H 2.14, *s*, 3H) to C-5 ($\delta$C 163.7).

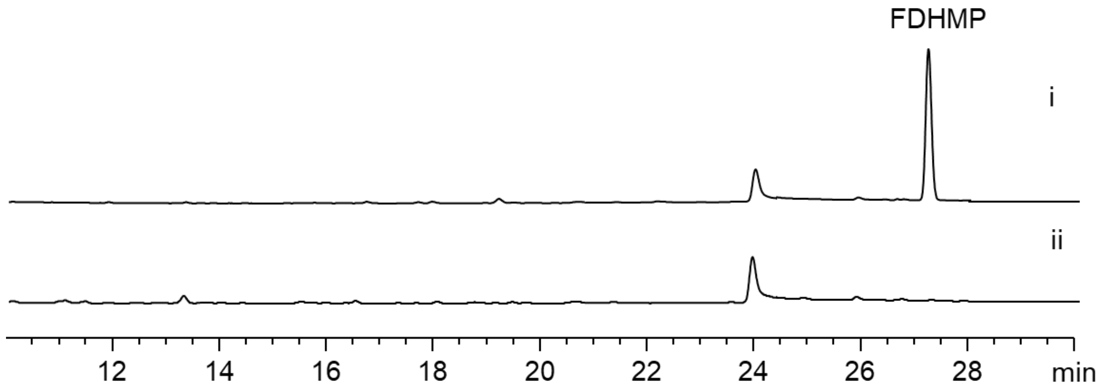


**Fig. S18.** HPLC analysis (254 nm) of *Ao*$_{M\text{-}2\text{-}3}$-*mpaB'* with the feeding of FDHMP. i, the intracellular extract of *Ao*$_{M\text{-}2\text{-}3}$-*mpaB'*/FDHMP; ii, the extracellular extract of *Ao*$_{M\text{-}2\text{-}3}$-*mpaB'*/FDHMP.

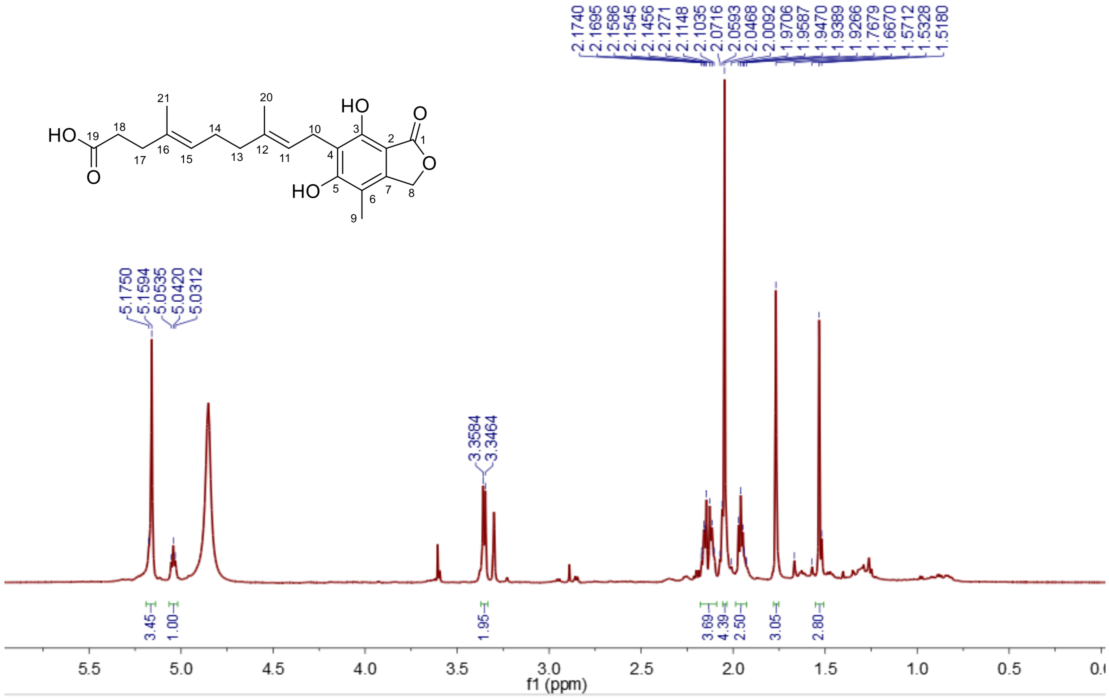

**Fig. S19.** [1]H NMR spectrum of FDHMP-3C in $CD_3OD$ (600 MHz).

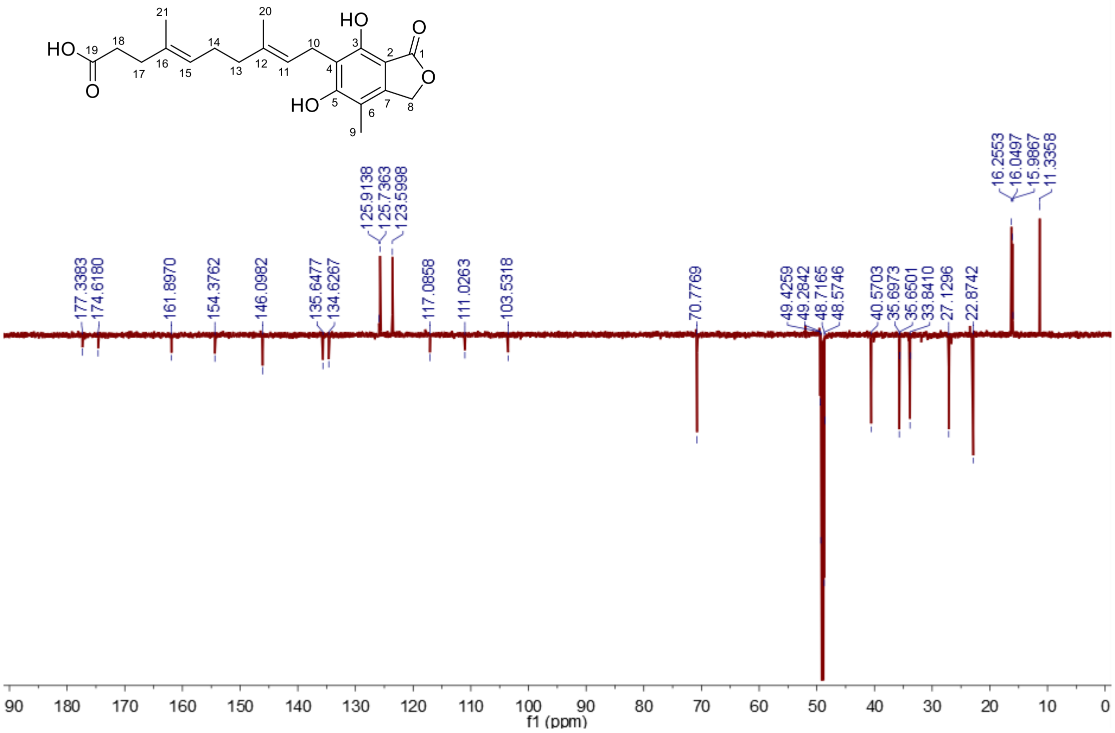


**Fig. S20.** $^{13}$C NMR spectrum of FDHMP-3C in $CD_3OD$ (150 MHz).

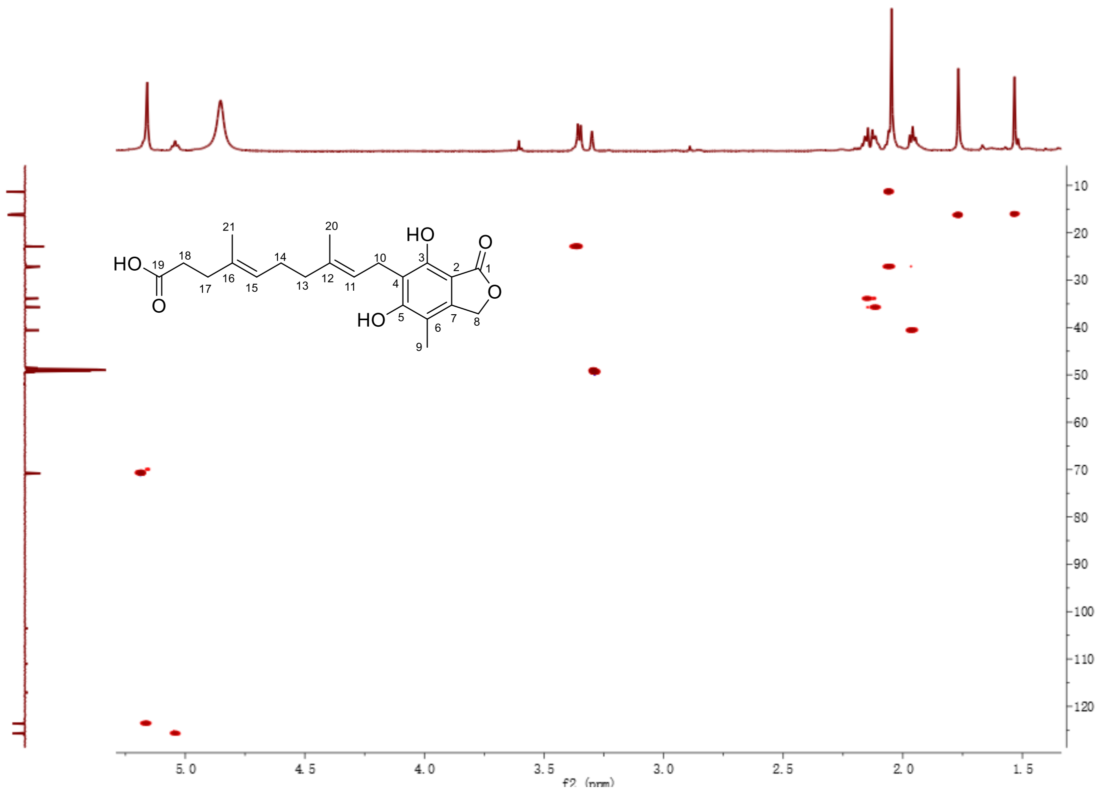


**Fig. S21.** HSQC spectrum of FDHMP-3C in $CD_3OD$.

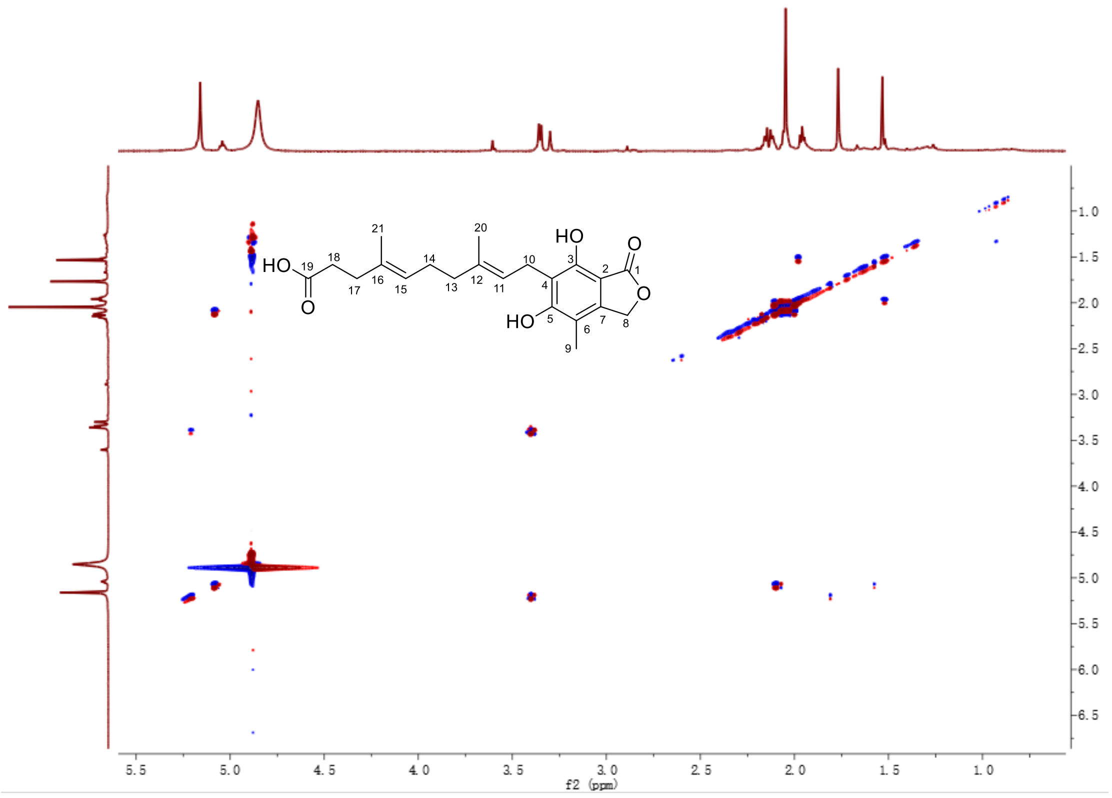


**Fig. S22.** $^{1}H$-$^{1}H$ COSY spectrum of FDHMP-3C in $CD_3OD$.

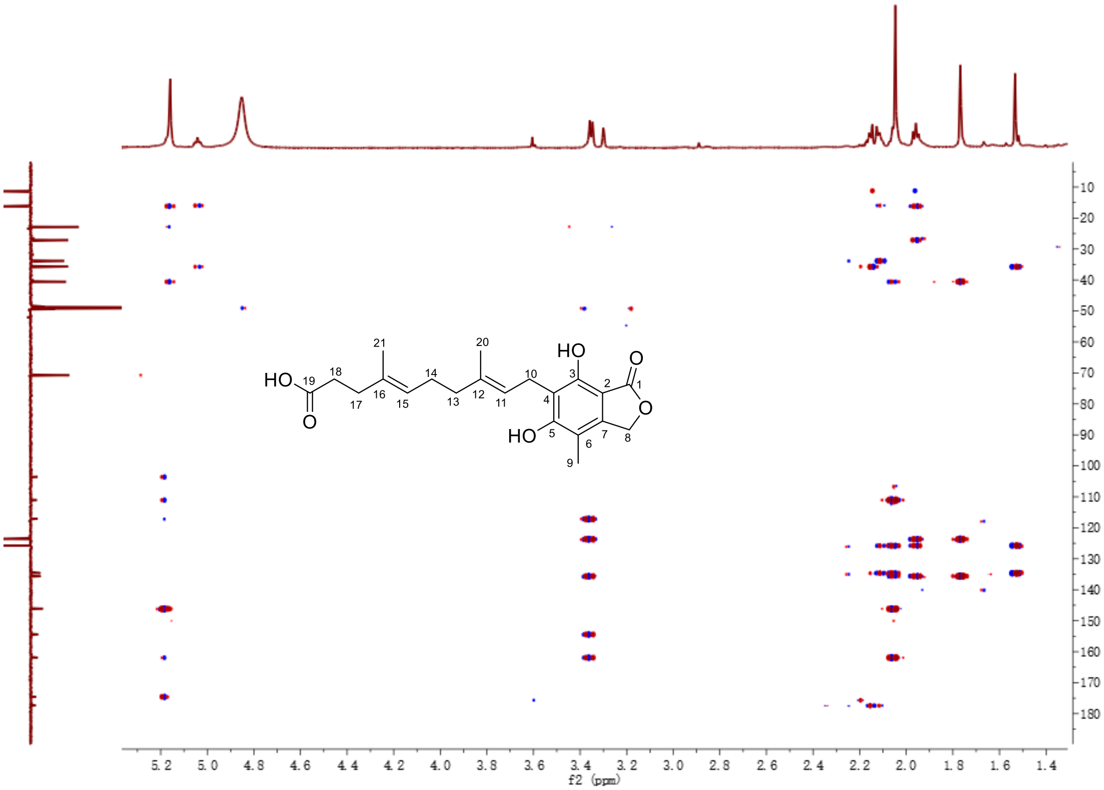


**Fig. S23.** HMBC spectrum of FDHMP-3C in $CD_3OD$.

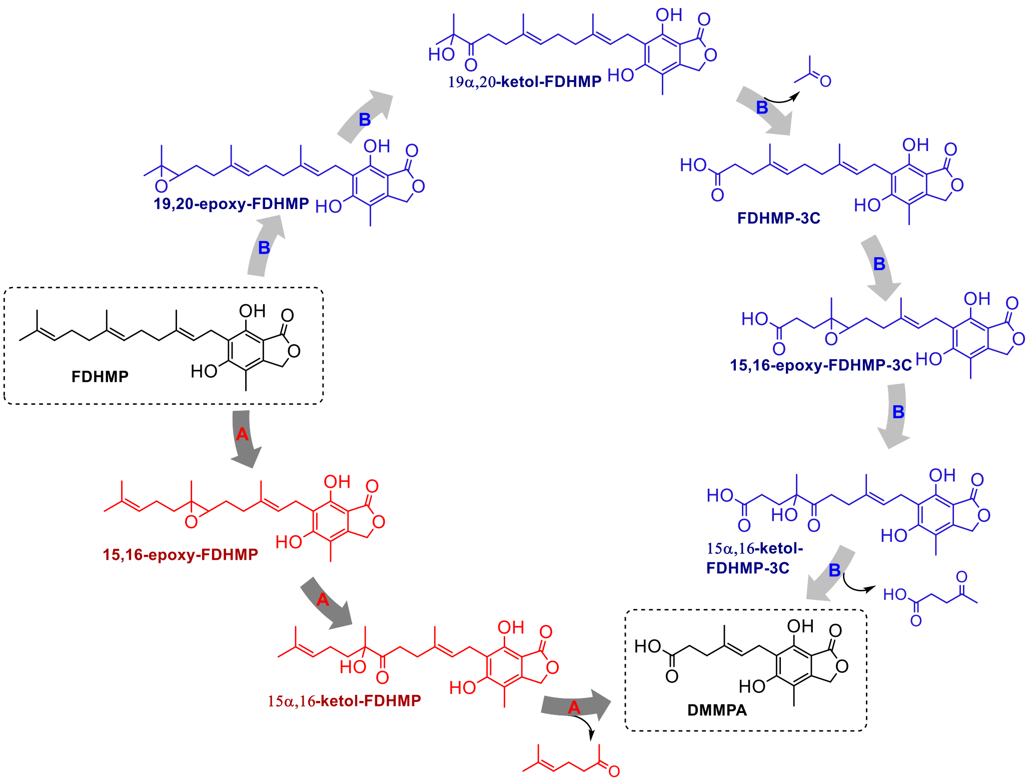


**Fig. S24.** The two putative oxidative cleavage(s) of the $C_{15}$ farnesyl side chain during MPA biosynthesis. A: direct central cleavage; B: two successive terminal cleavages.

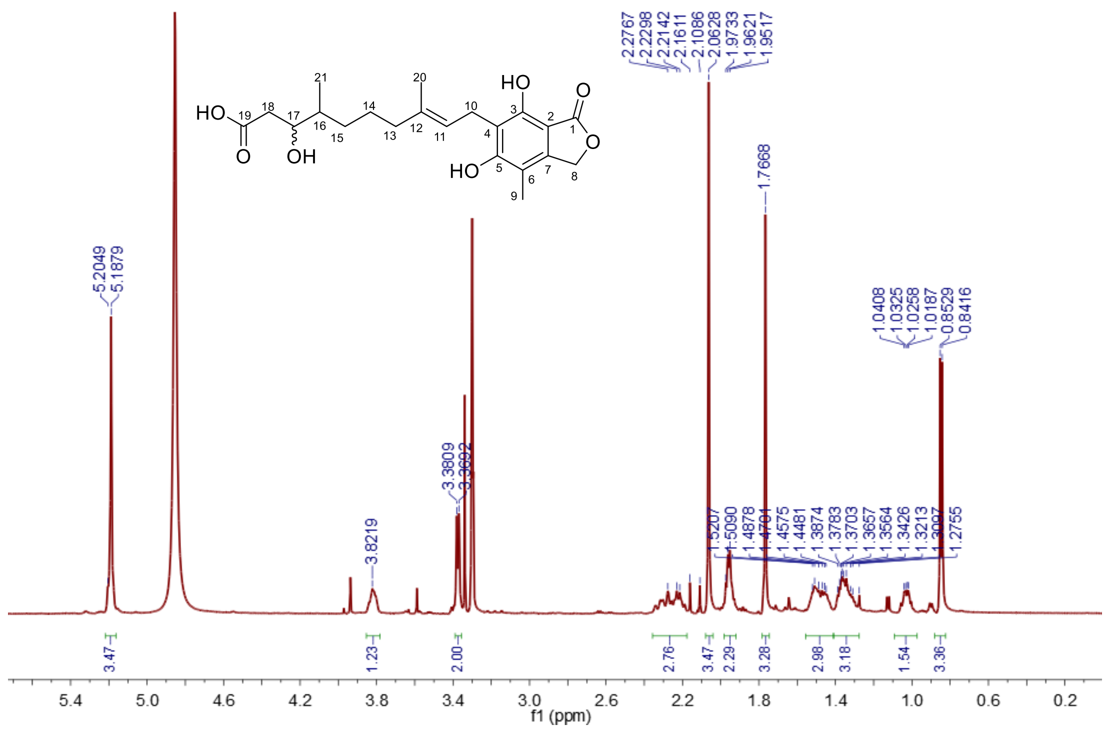


**Fig. S25.** [1]H NMR spectrum of FDHMP-d1 in $CD_3OD$ (600 MHz).

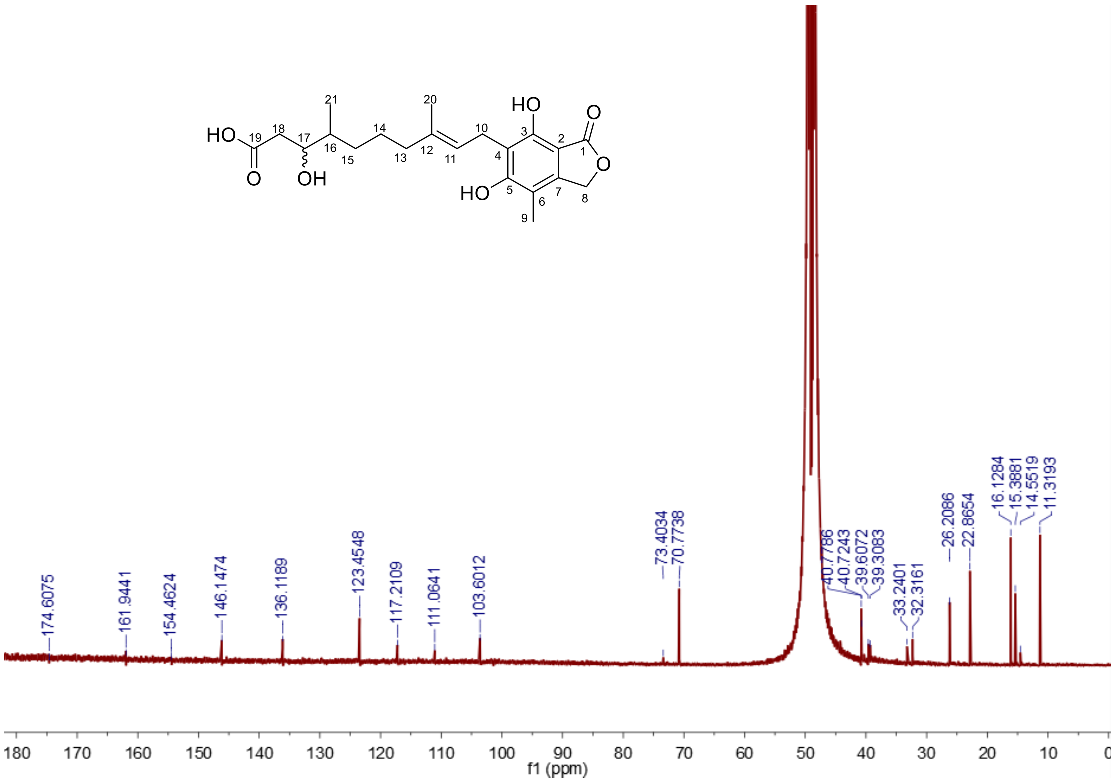


**Fig. S26.** $^{13}$C NMR spectrum of FDHMP-d1 in $CD_3OD$ (150 MHz).

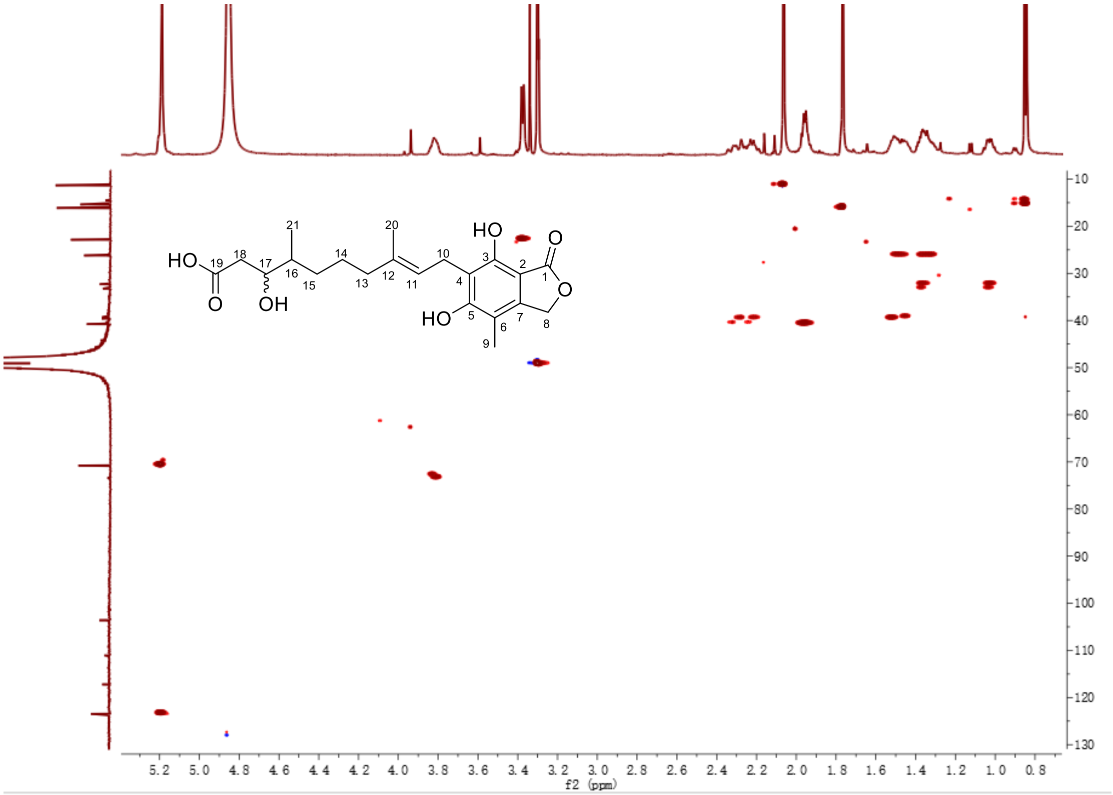


**Fig. S27.** HSQC spectrum of FDHMP-d1 in $CD_3OD$.

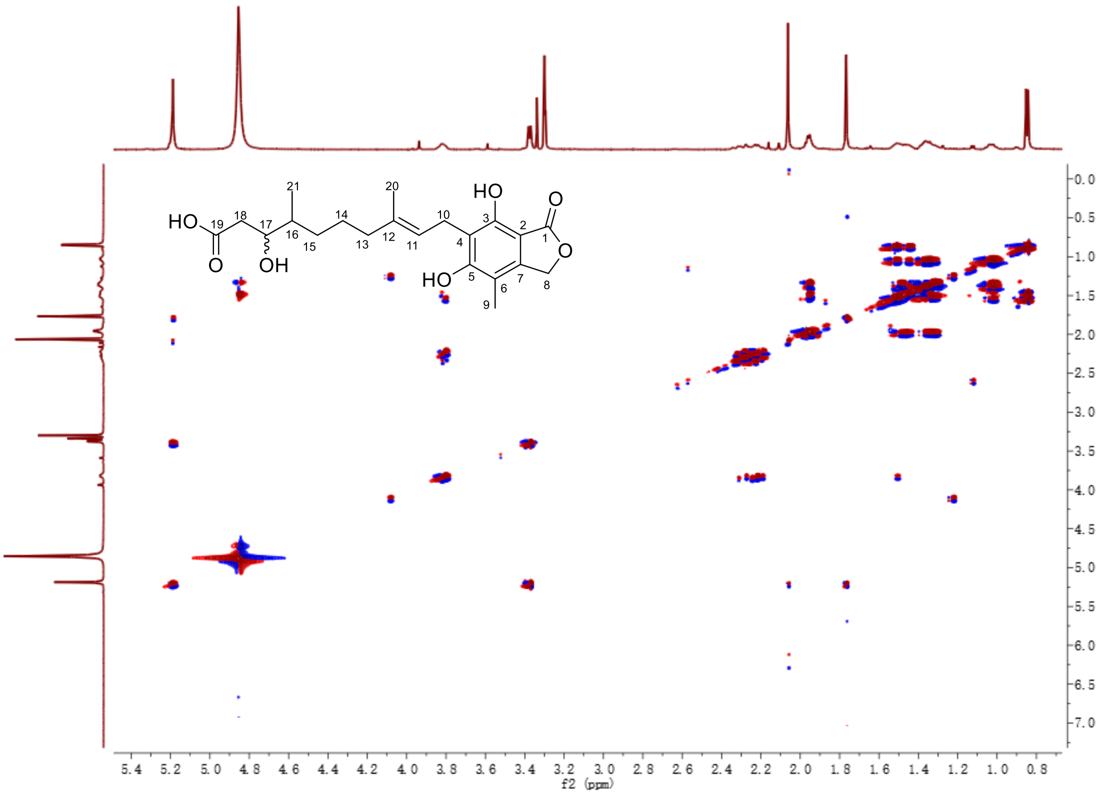


**Fig. S28.** $^{1}H$-$^{1}H$ COSY spectrum of FDHMP-d1 in $CD_3OD$.

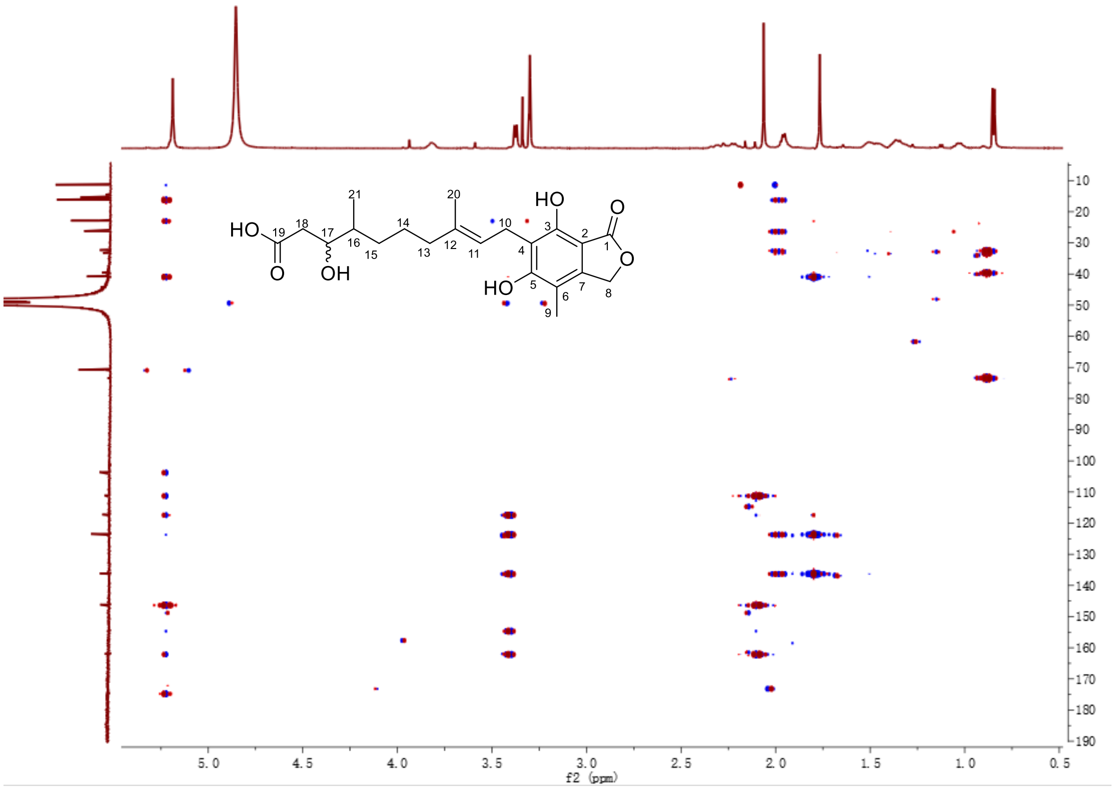


**Fig. S29.** HMBC spectrum of FDHMP-d1 in $CD_3OD$.

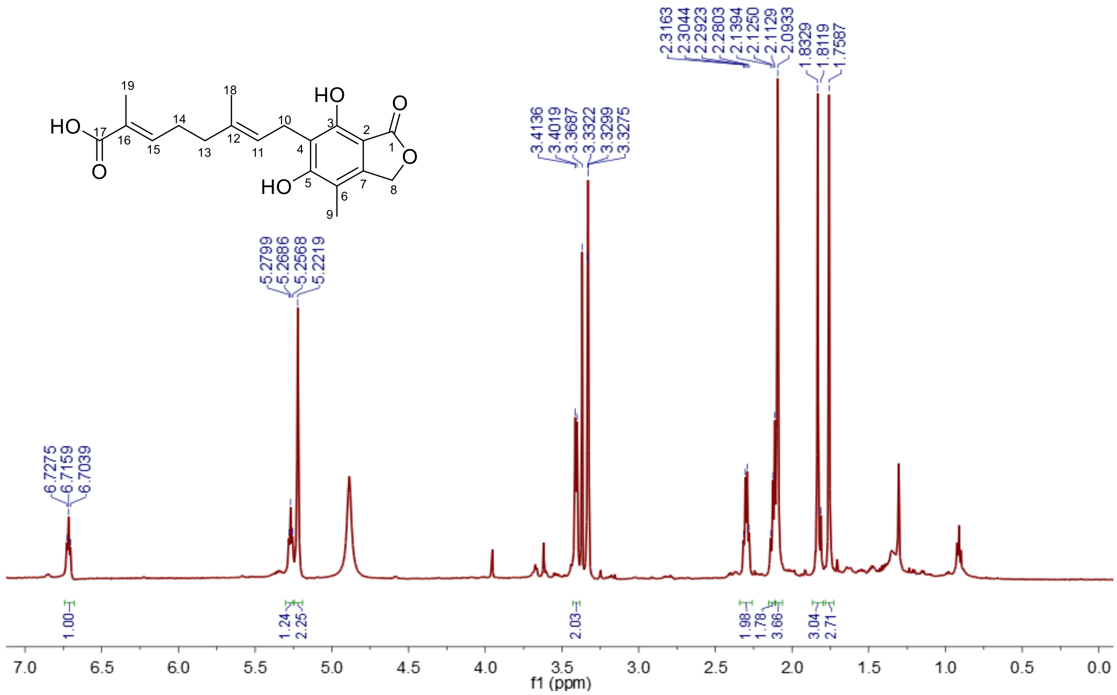


**Fig. S30.** $^1$H NMR spectrum of FDHMP-d2 in $CD_3OD$ (600 MHz).

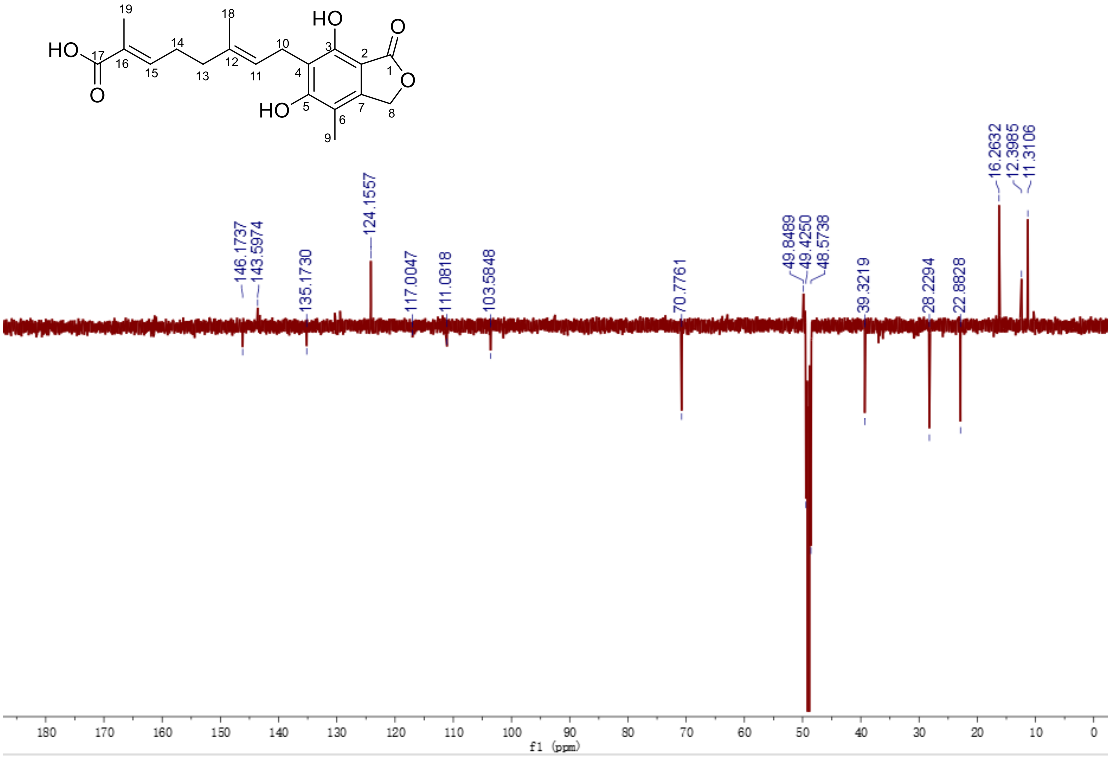


**Fig. S31.** $^{13}$C NMR spectrum of FDHMP-d2 in $CD_3OD$ (150 MHz).

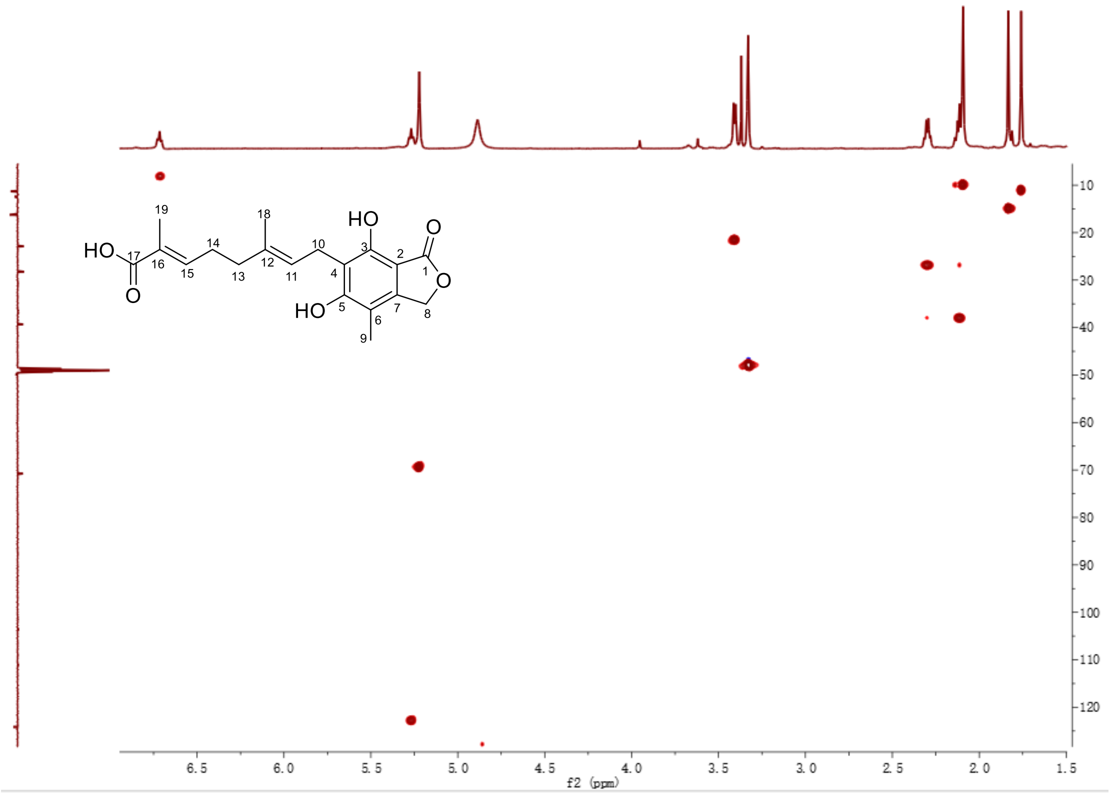


**Fig. S32.** HSQC spectrum of FDHMP-d2 in $CD_3OD$.

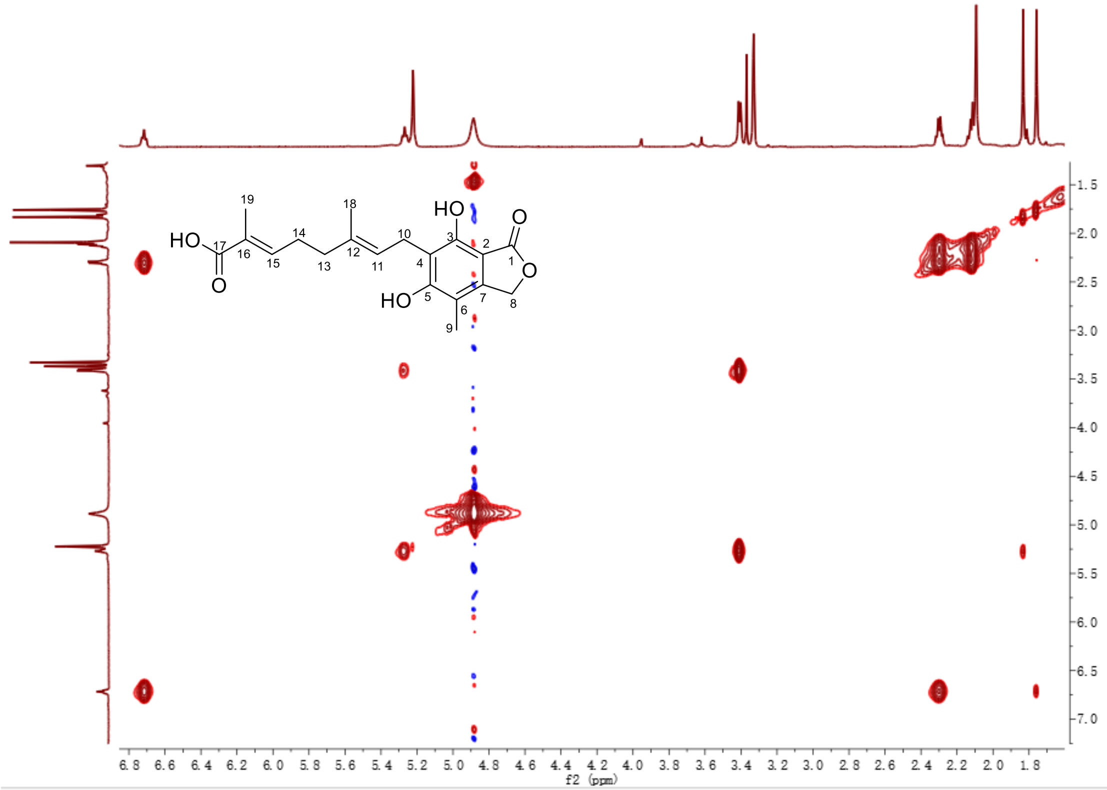


**Fig. S33.** [1]H-[1]H COSY spectrum of FDHMP-d2 in $CD_3OD$.

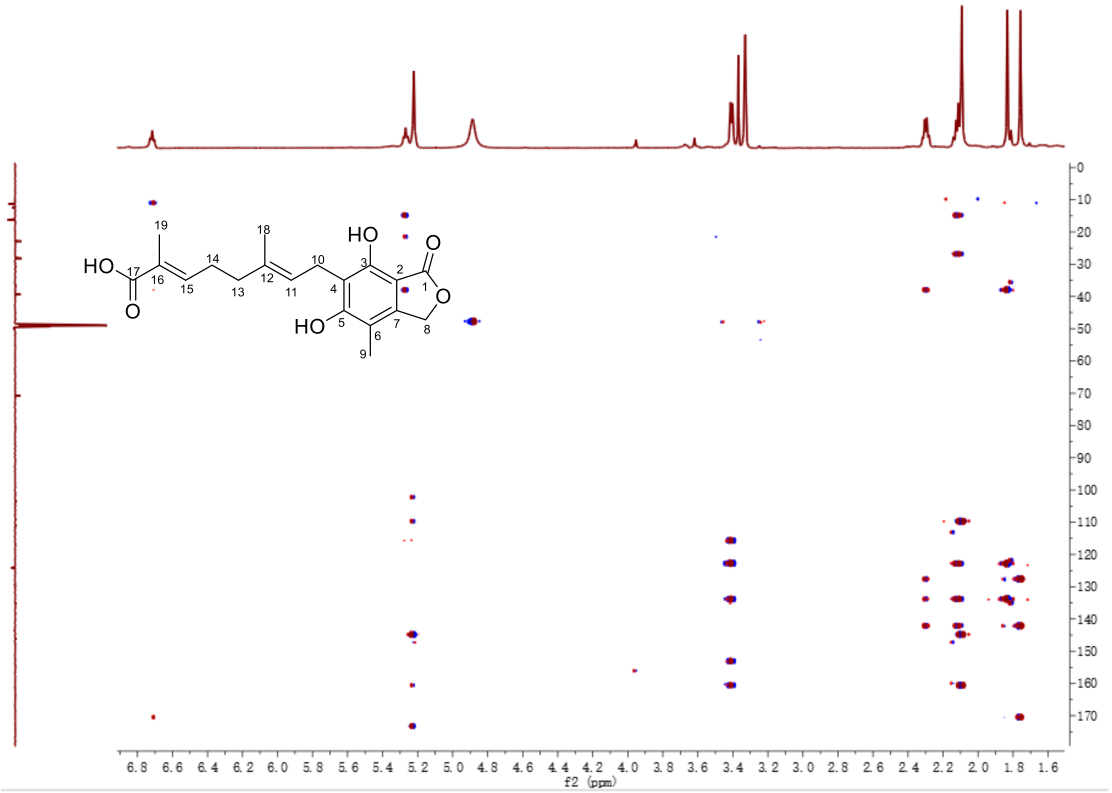


**Fig. S34.** NOESY spectrum of FDHMP-d2 in $CD_3OD$.

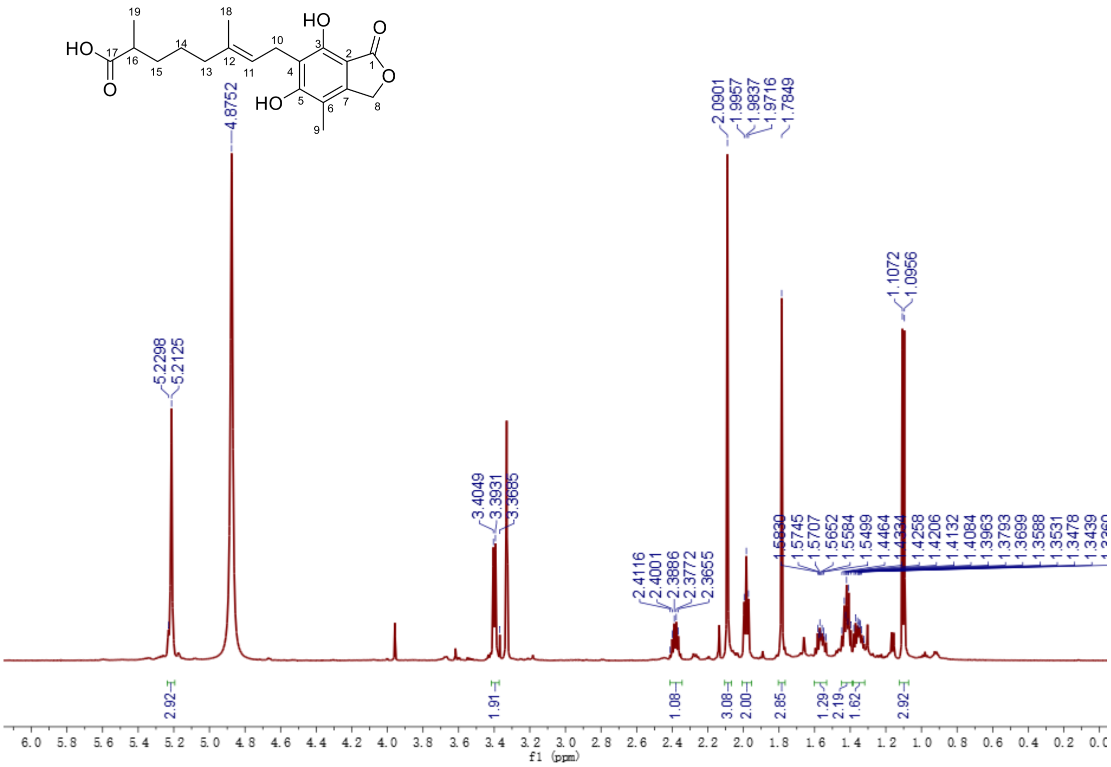

**Fig. S35.** $^{1}$H NMR spectrum of FDHMP-d3 in $CD_3OD$ (600 MHz).

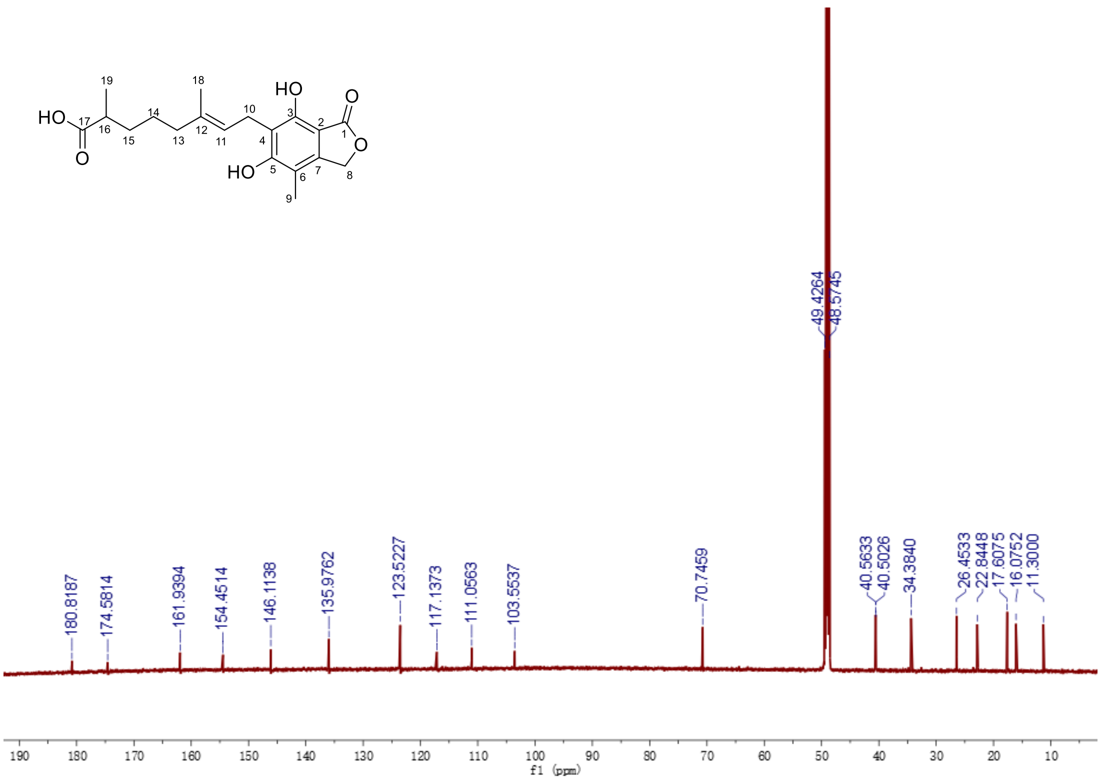


**Fig. S36.** $^{13}$C NMR spectrum of FDHMP-d3 in $CD_3OD$ (150 MHz).

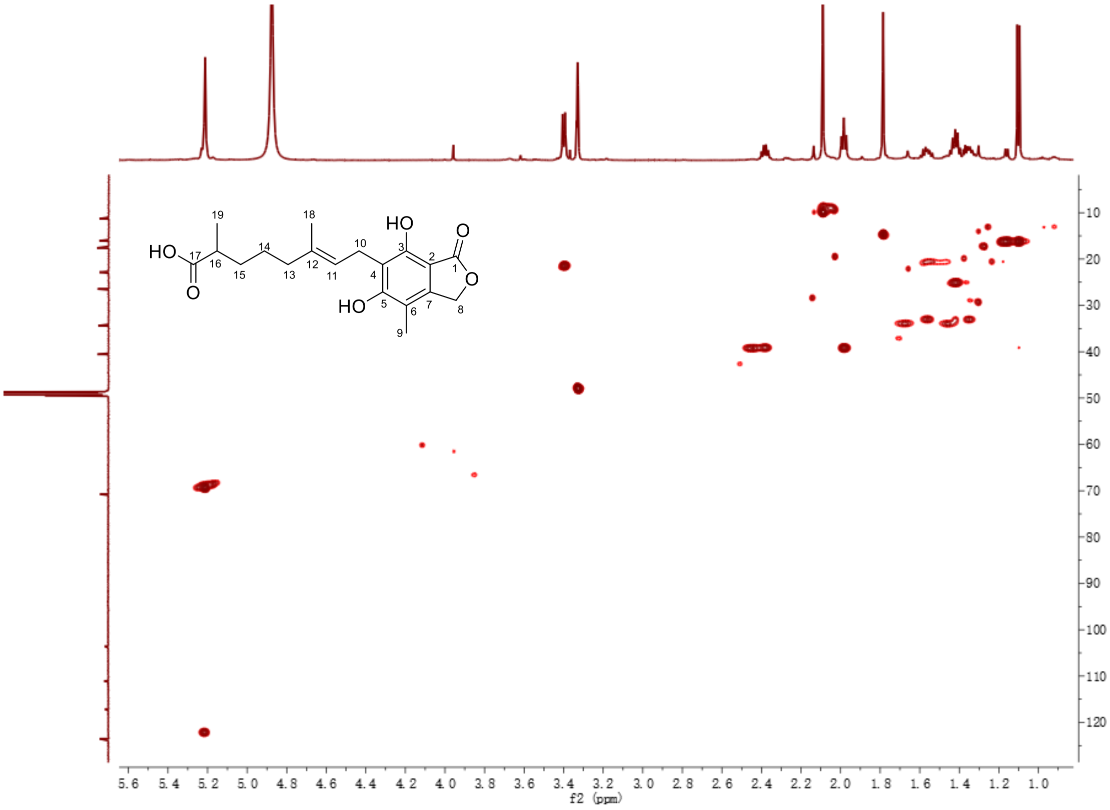


**Fig. S37.** HSQC spectrum of FDHMP-d3 in $CD_3OD$.

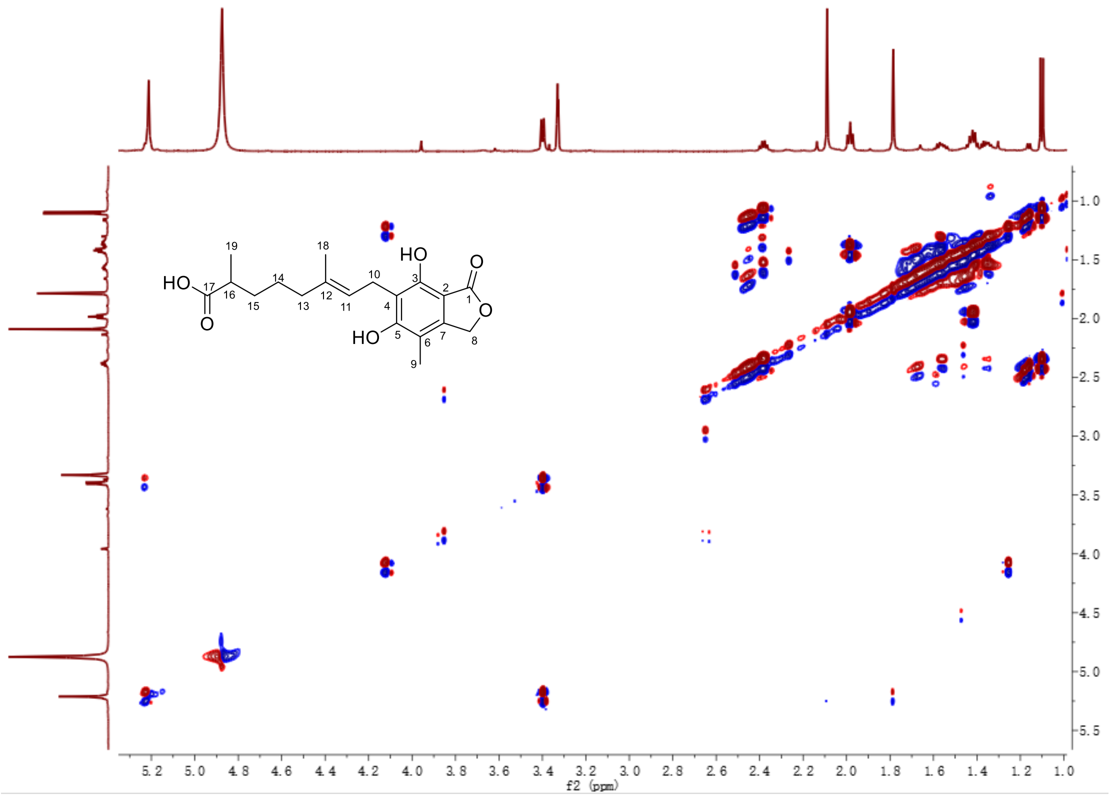


**Fig. S38.** $^1H$-$^1H$ COSY spectrum of FDHMP-d3 in $CD_3OD$.

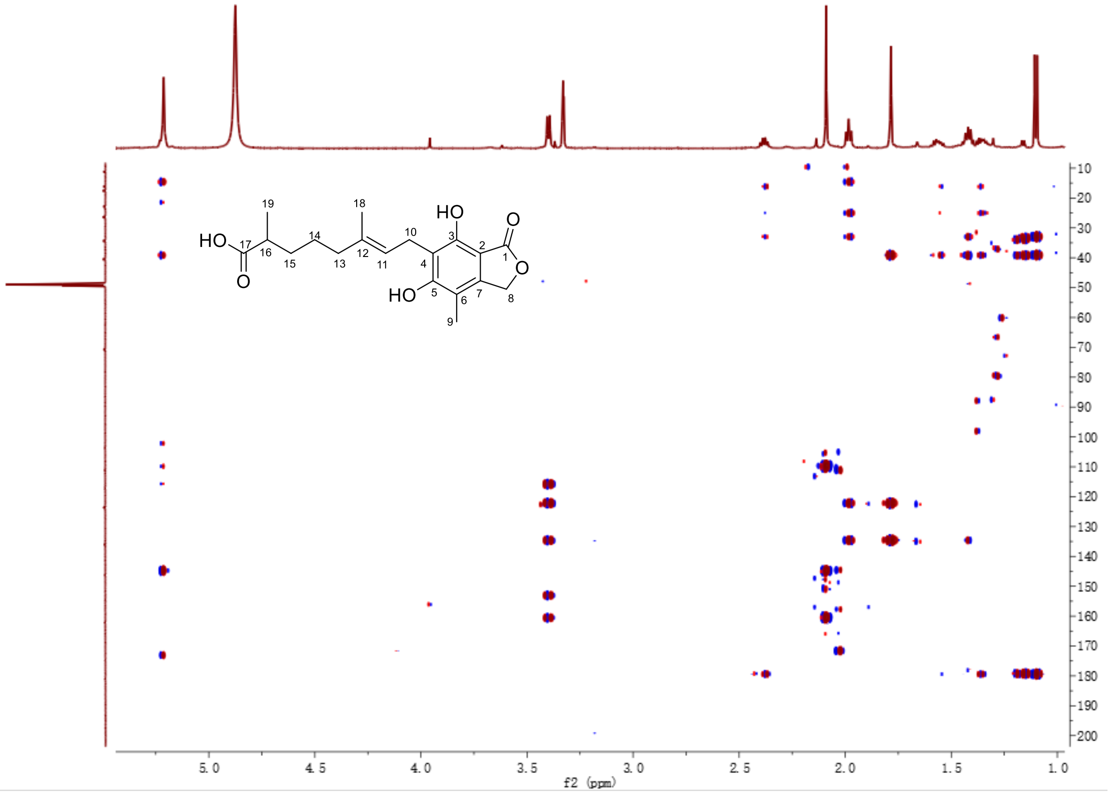


**Fig. S39.** HMBC spectrum of FDHMP-d3 in $CD_3OD$.

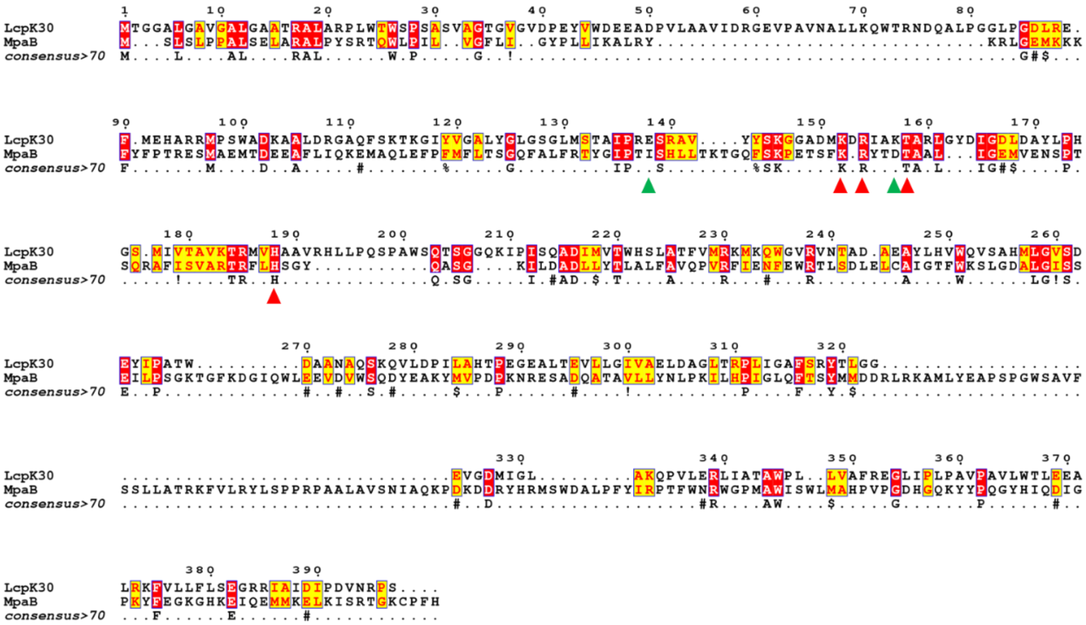

**Fig. S40.** Protein sequence alignment of MpaB' and Lcp$_{K30}$. Sequence analysis was performed using Expresso through the T-COFFEE online service, and the result was exported by ESPript 3.0 (19, 20). The red and green triangles denote the residues that are conserved and not conserved in the active site, respectively.

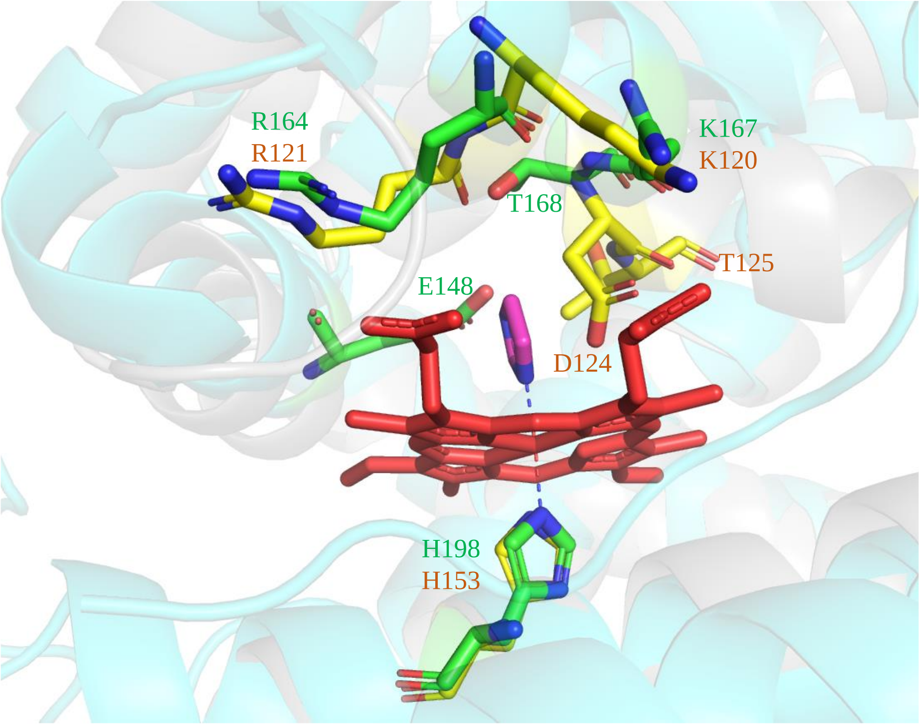


**Fig. S41.** The superposition of the modelled MpaB' (Gray) and the crystal structure of Lcp$_{K30}$ (Cyan; PDB ID: 5O1L) from *Streptomyces* sp. K30 (21). In Lcp$_{K30}$, the heme axial ligand H198, and the acitive site residues K167, T168, and R164 are shown as sticks in green. The structurally aligned residues in the modelled MpaB' are shown as sticks in yellow.

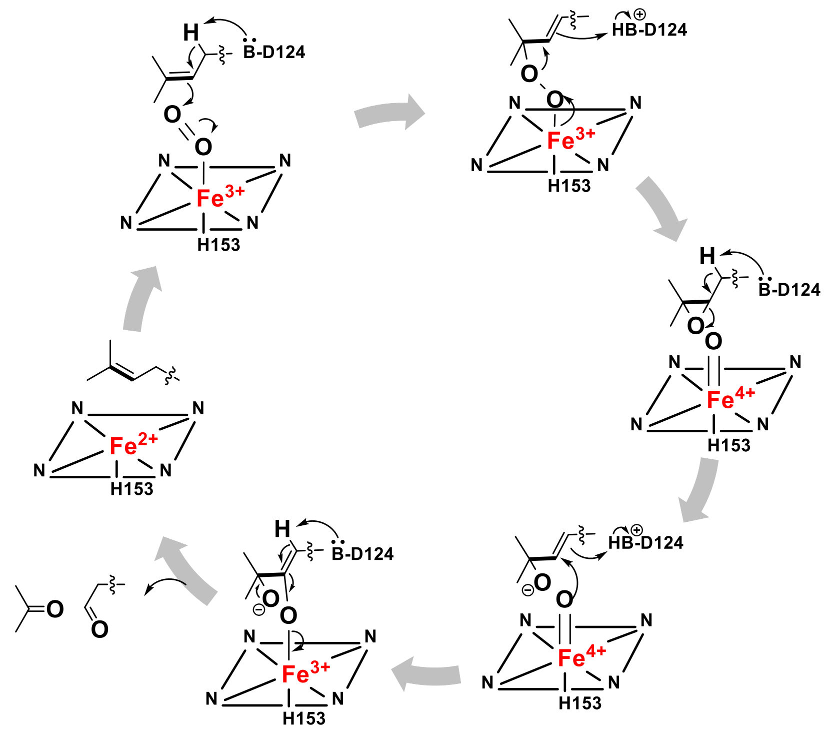


**Fig. S42.** The putative catalytic mechanism of MpaB'.

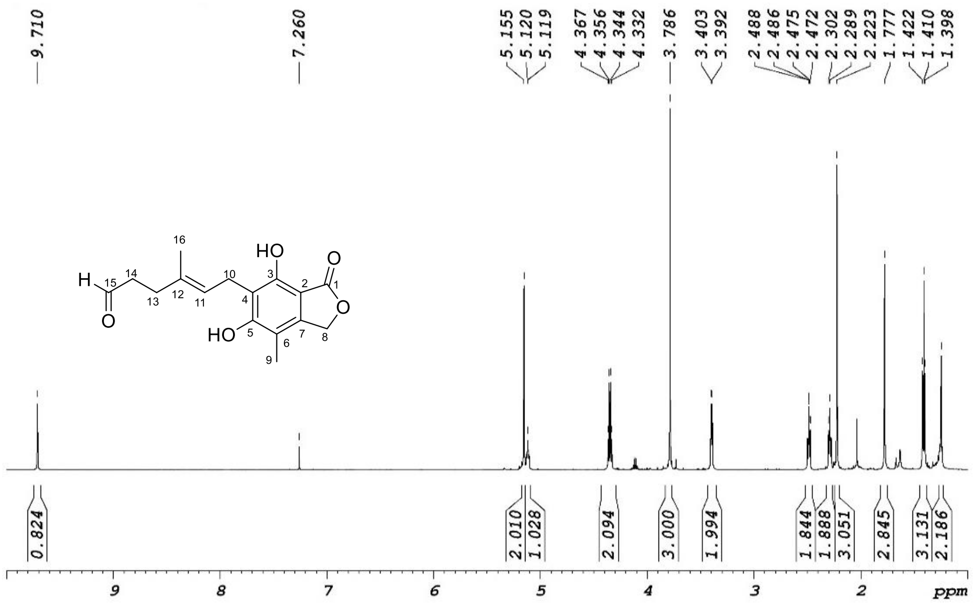


**Fig. S43.** $^{1}$H NMR spectrum of mycophenolic aldehyde in $CDCl_3$ (600 MHz).

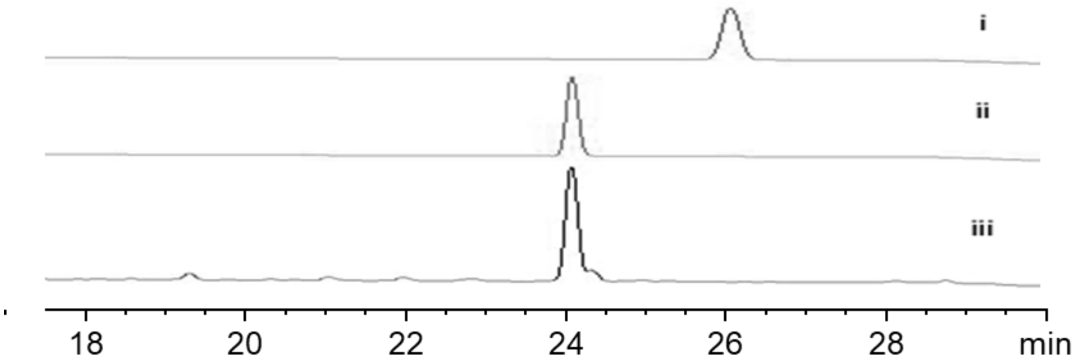


**Fig. S44.** HPLC analysis (254 nm) of *Ao*$_{M-2-3}$ with the feeding of mycophenolic aldehyde. i, synthesized mycophenolic aldehyde; ii, the extracellular extract of *Ao*$_{M-2-3}$/mycophenolic aldehyde; iii, MPA standard.

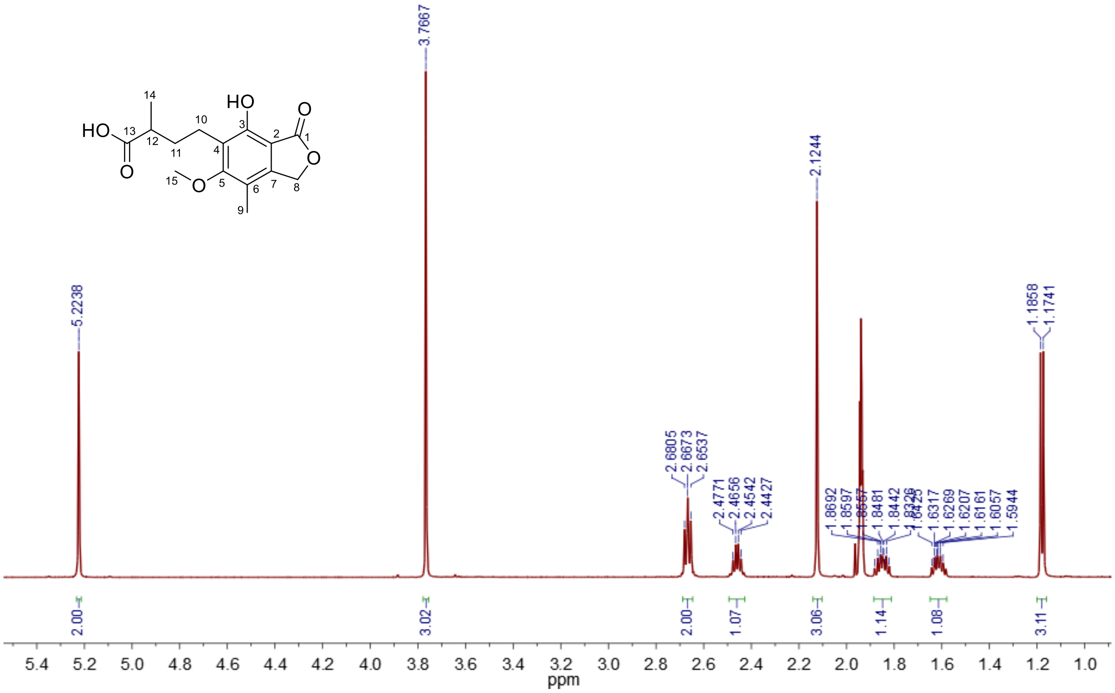

**Fig. S45.** $^{1}$H NMR spectrum of MFDHMP-d4 in $CD_3CN$ (600 MHz).

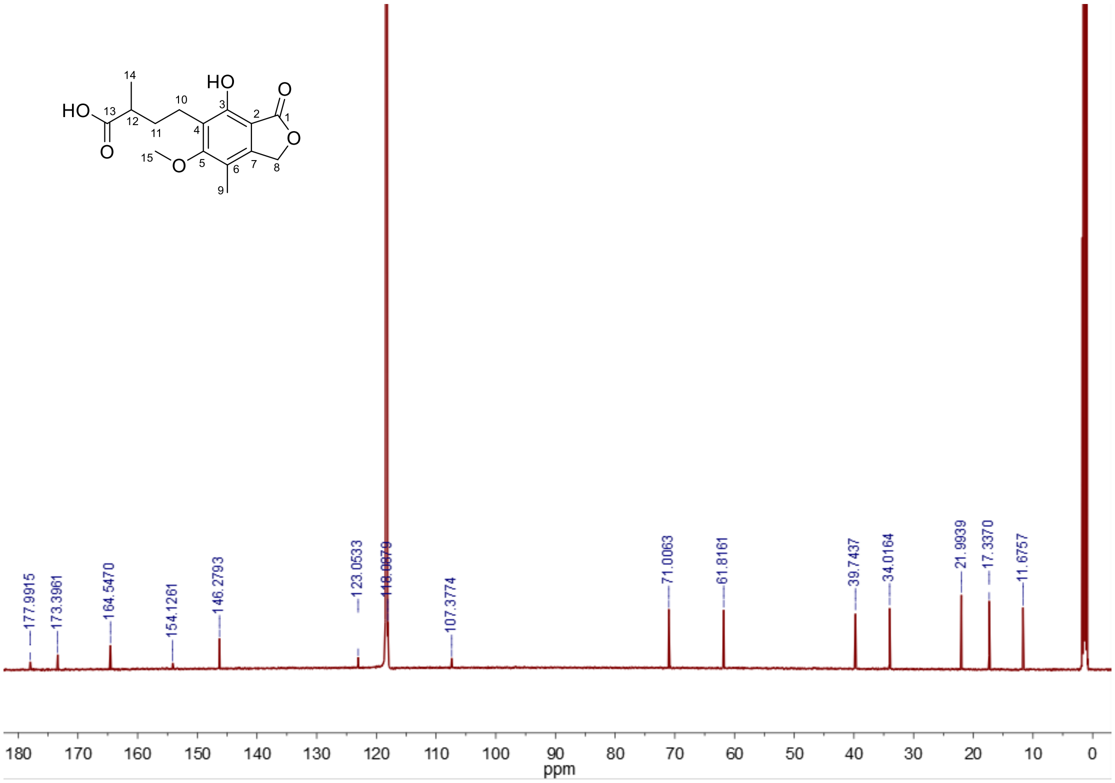


**Fig. S46.** $^{13}$C NMR spectrum of MFDHMP-d4in $CD_3CN$ (150 MHz).

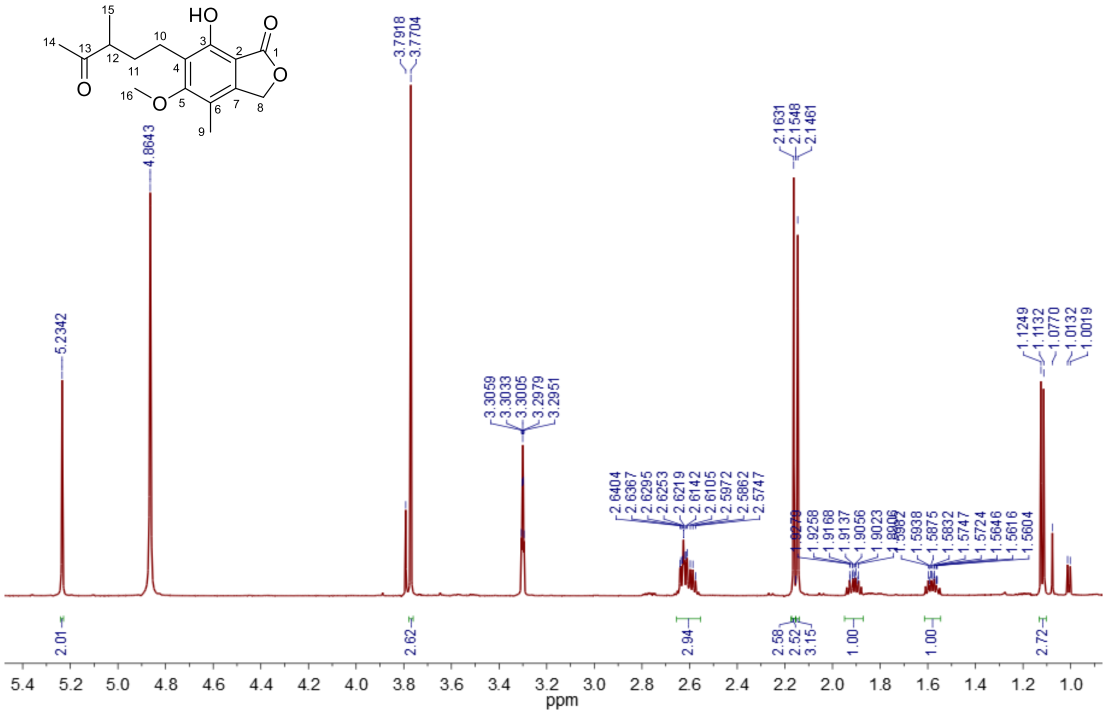

**Fig. S47.** [1]H NMR spectrum of MFDHMP-d5 in $CD_3OD$ (600 MHz).

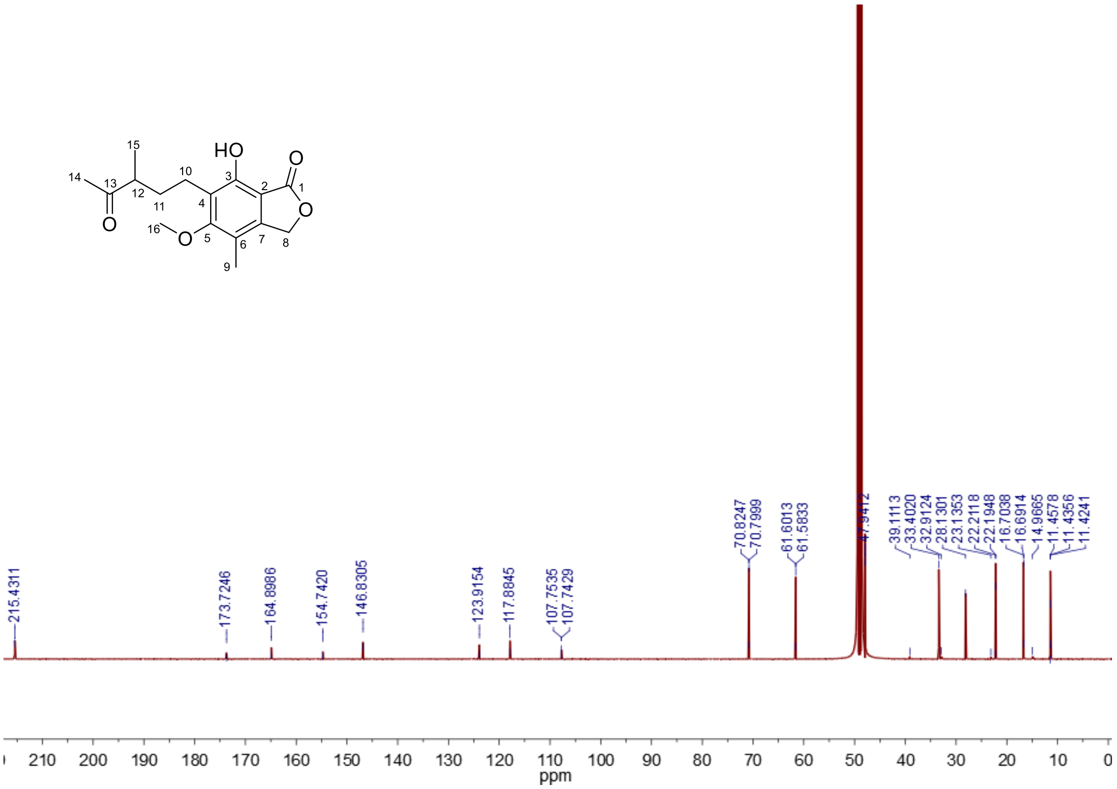


**Fig. S48.** $^{13}$C NMR spectrum of MFDHMP-d5 in $CD_3OD$ (150 MHz).

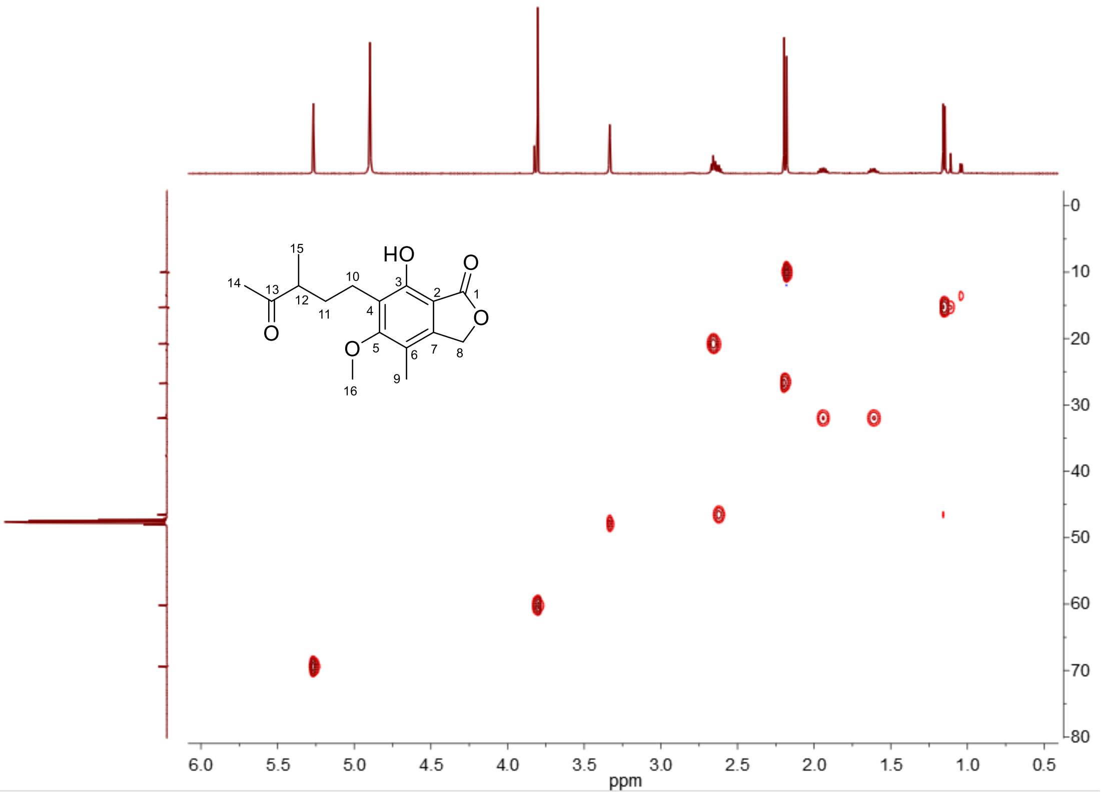


**Fig. S49.** HSQC spectrum of MFDHMP-d5 in $CD_3OD$.

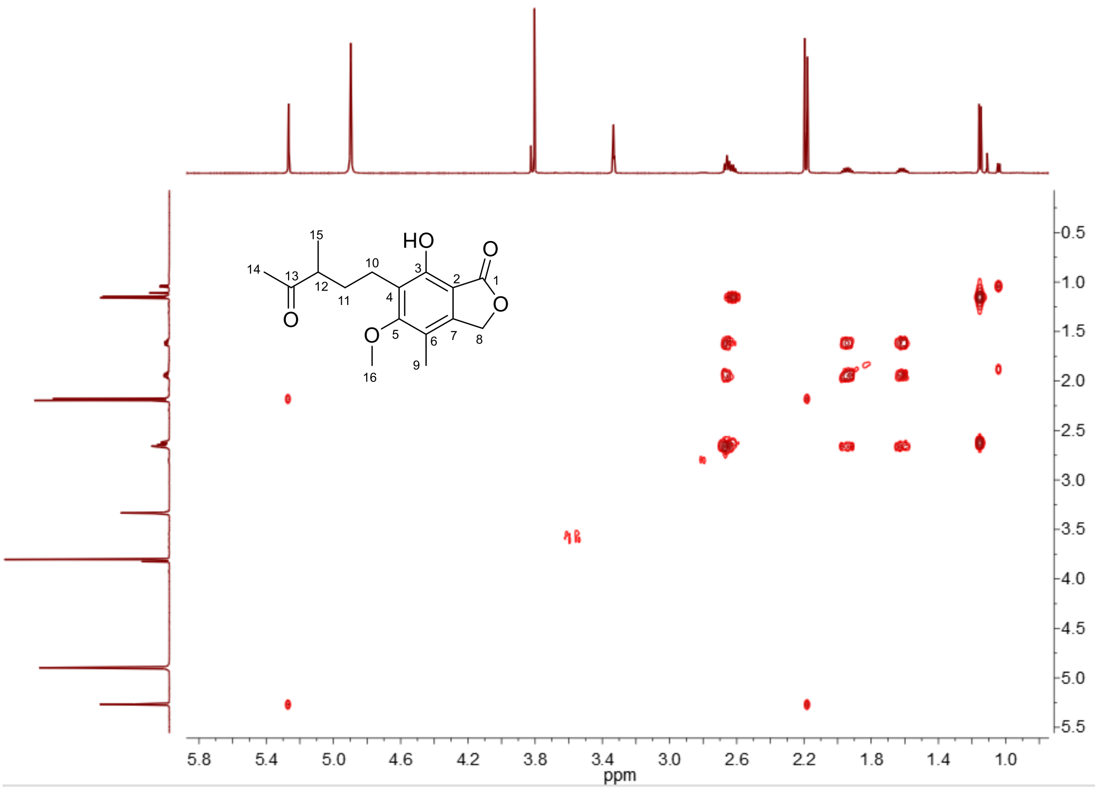


**Fig. S50.** $^{1}H$-$^{1}H$ COSY spectrum of MFDHMP-d5 in $CD_3OD$.

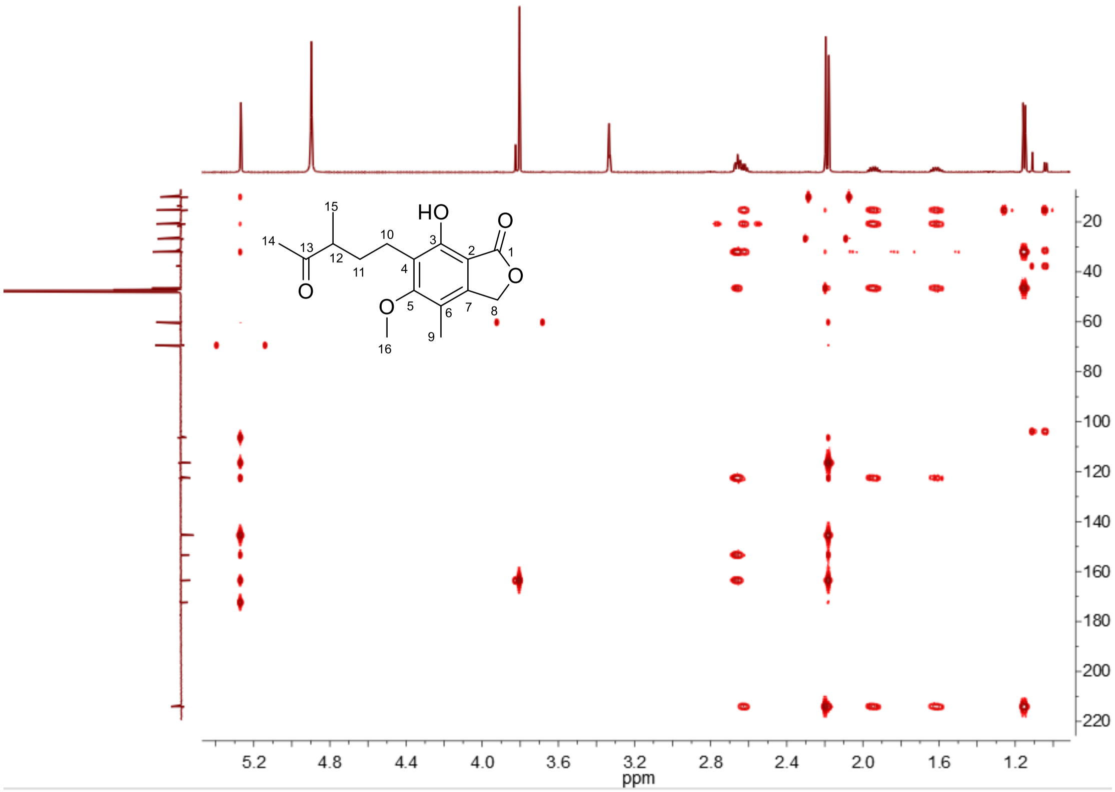


**Fig. S51.** HMBC spectrum of MFDHMP-d5 in $CD_3OD$.

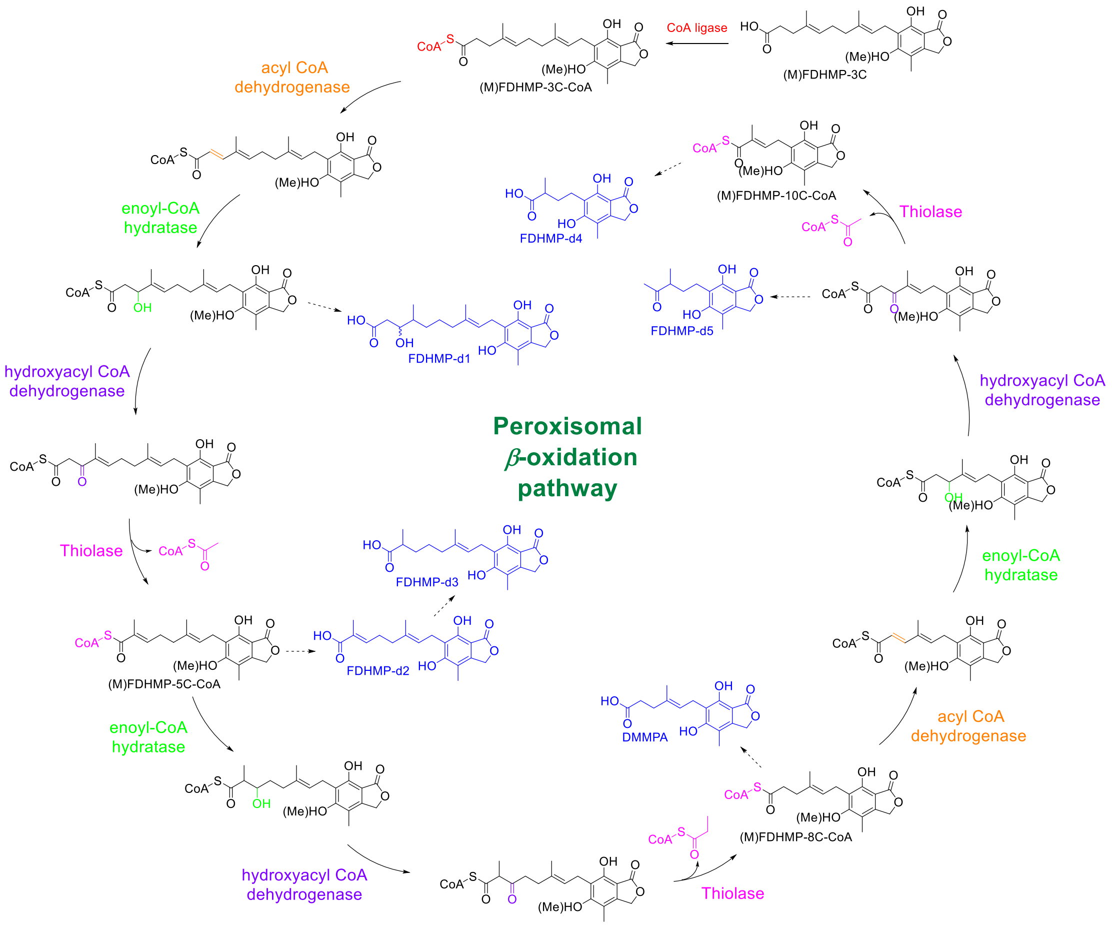


**Fig. S52.** The putative $\beta$-oxidation pathway of FDHMP-3C in fungal peroxisomes (Solid arrows: the $\beta$-oxidation pathway; dashed arrows: the shunt pathways. The newly installed functional groups are colored after the same-colored enzymes. The structures in blue are the intermediates identified from *Ao*M-2-3-*mpaA'*-*mpaB'*/DHMP and *Ao*M-2-3-*mpaA'*-*mpaB'*-*mpaH'*/DHMP).

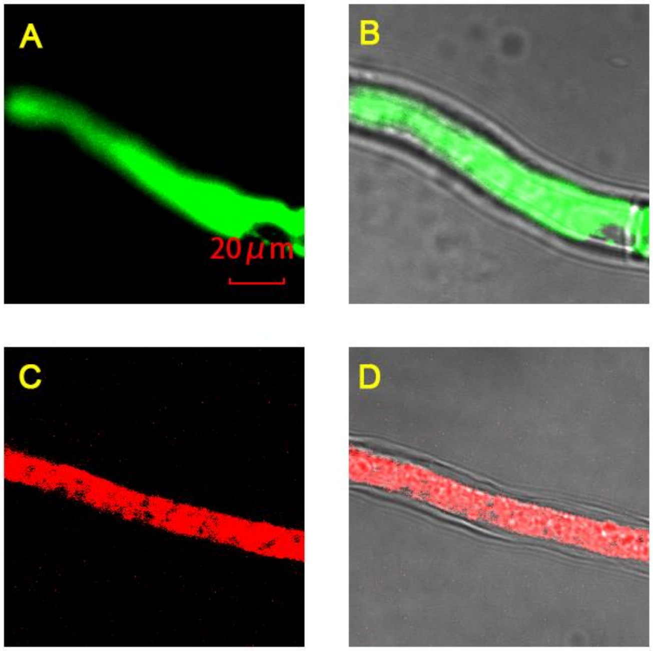


**Fig. S53.** High-resolution confocal imaging of *Ao*M-2-3-*gfp* and *Ao*M-2-3-*rfp*. **A**, The localization of GFP; **B**, The overlay of **A** with the bright field; **C**, The localization of RFP; **D**, The overlay of **C** with the bright field.

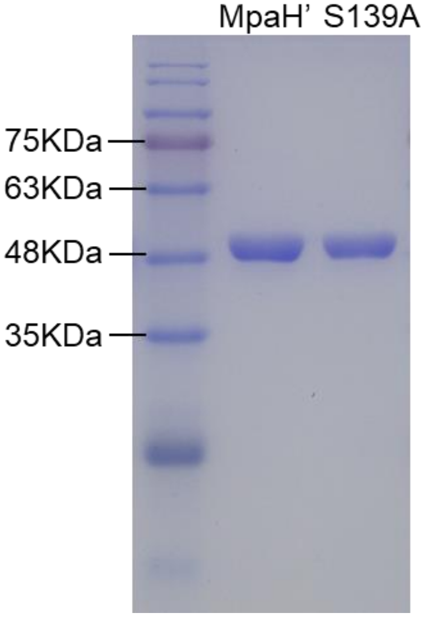


**Fig. S54.** SDS-PAGE analysis of purified MpaH’ and MpaH’$_{S139A}$.

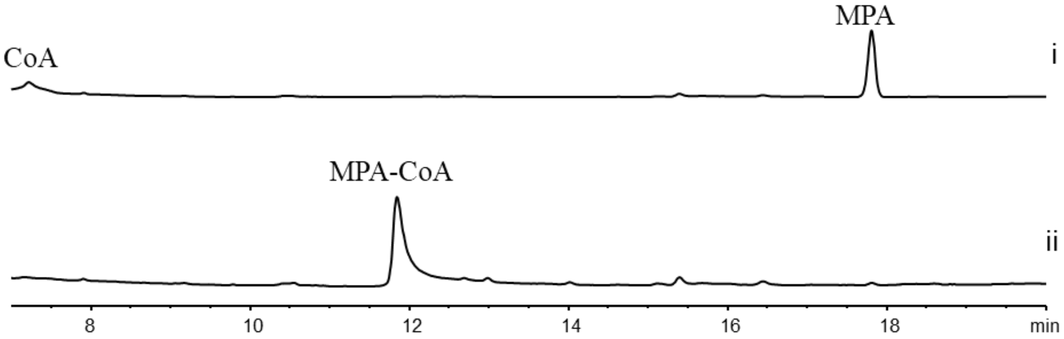


**Fig. S55.** HPLC analysis (254 nm) of *in vitro* conversions catalyzed by MpaH'. i, MpaH' /MPA-CoA; ii, MpaH'$^{S139A}$/MPA-CoA.

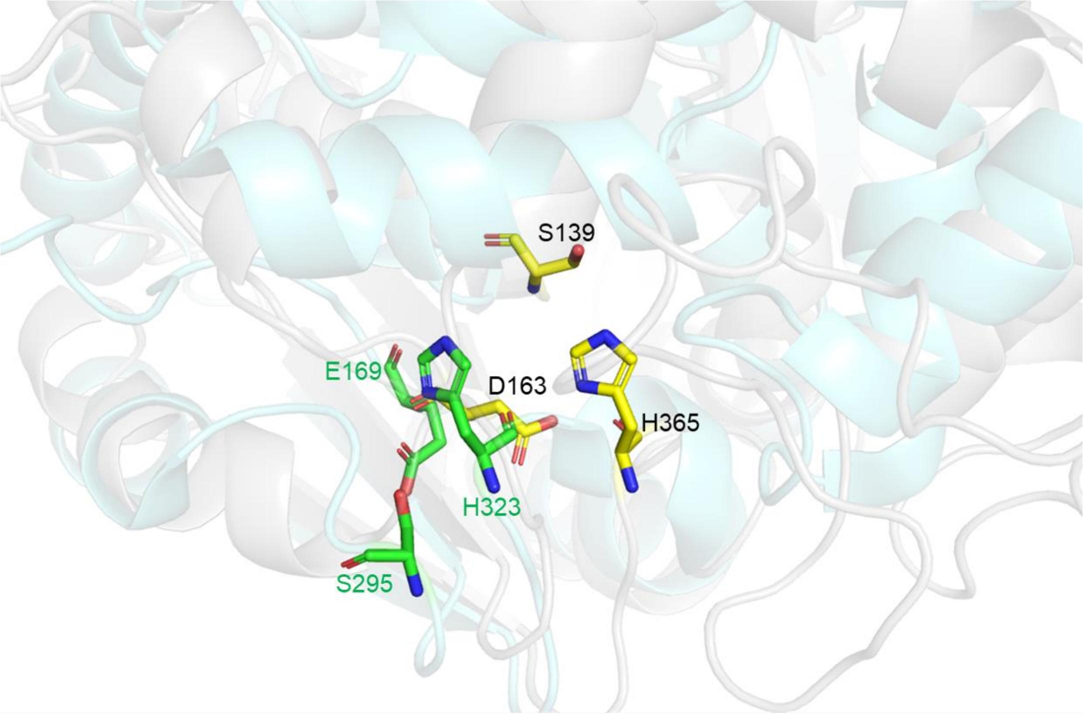


**Fig. S56.** The superposition of the modelled MpaH' (Gray) and the crystal structure of Lpx1 (Cyan, PDB ID: 2Y6V) from *Saccharomyces cerevisiae* (*33*). The catalytic triad of Lpx1 is shown as sticks in green, while the putative catalytic triad of MpaH' is shown as sticks in yellow.

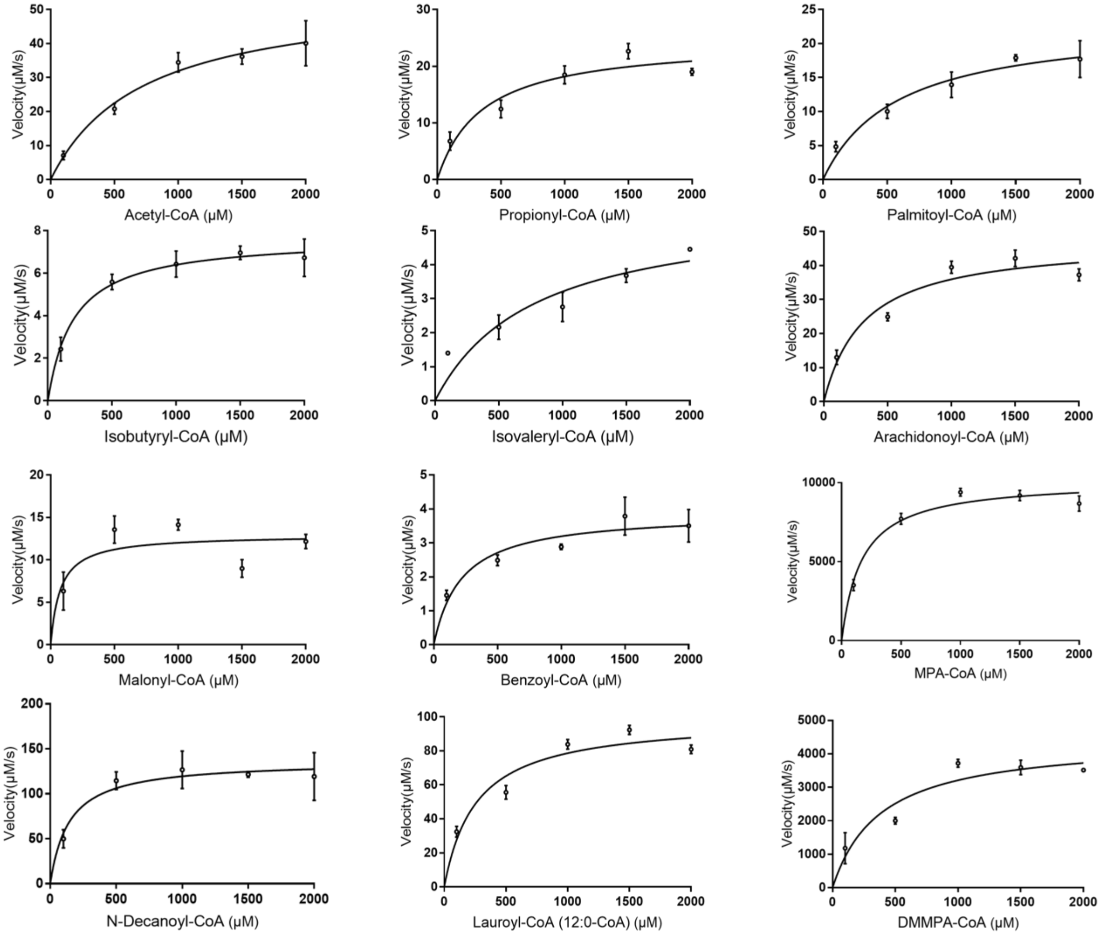


**Fig. S57.** The steady-state kinetic curves of MpaH’ toward twelve acyl-CoA-esters.

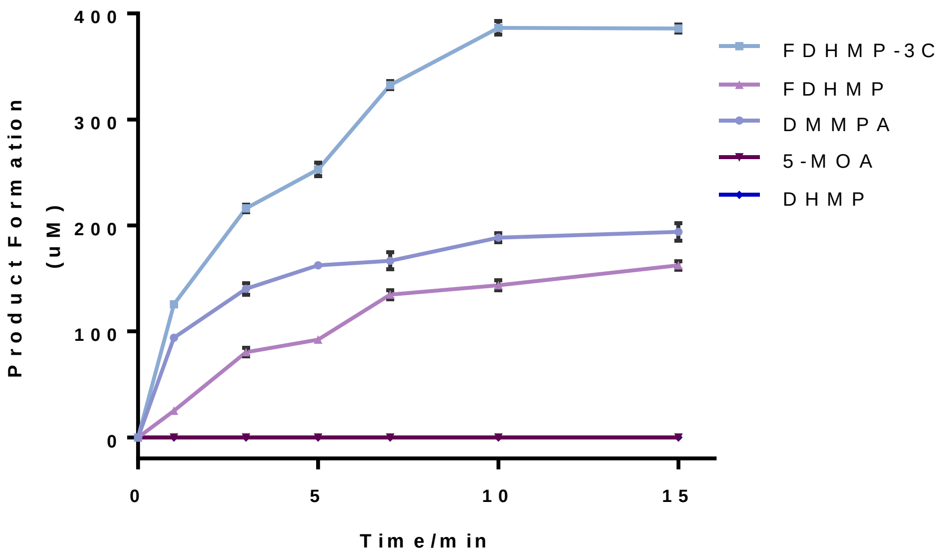


**Fig. S58.** Methylation product formation rate of *in vitro* reactions catalyzed by MpaG’ (1 μM) toward different substrates (500 μM).

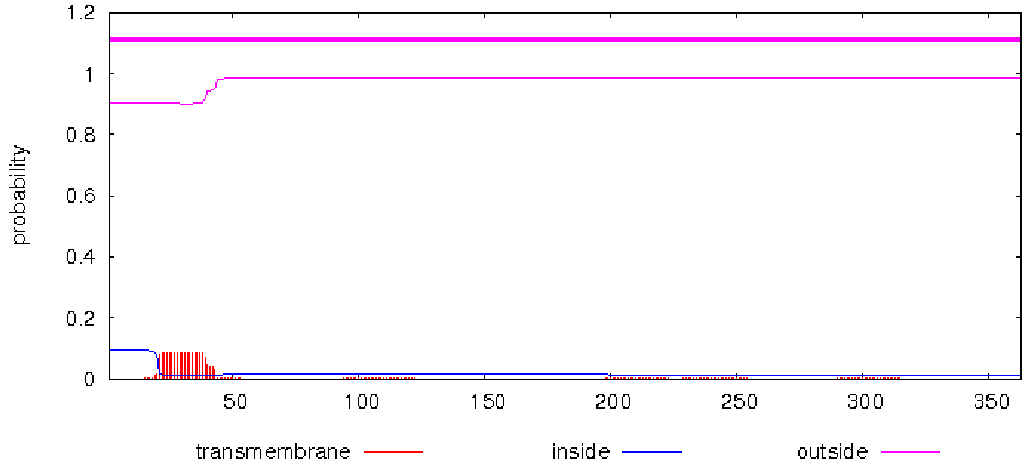


**Fig. S59.** Prediction of transmembrane regions in MpaB' by TMHMM Server v. 2.0 (http://www.cbs.dtu.dk/services/TMHMM/).

**Table S1.** Predicted functions of the open reading frames shown in *SI Appendix*, Fig. S2.

| Protein | Amino acids | Putative function | Nearest homologue (enzyme, origin) | Identity (%) | Accession number |
|---|---|---|---|---|---|
| MpaA’ | 331 | Prenyltransferase | MpaA *[P. brevicompactum]* | 94.3 | AJG44379.1 |
| MpaB’ | 423 | Dephospho-CoA kinase | MpaB *[P.brevicompactum]* | 81.8 | AJG44380.1 |
| MpaC’ | 2190 | Polyketide synthase | MpaC *[P. brevicompactum]* | 95.9 | AJG44381.1 |
| MpaDE’ | 853 | P450 monooxygenase/hydrolase fusion | MpaDE *[P. brevicompactum]* | 98.6 | AJG44382.1 |
| MpaF’ | 548 | Inosine monophosphate dehydrogenase | MpaF *[P. brevicompactum]* | 95.8 | AJG44383.1 |
| MpaG’ | 398 | *O*-methyltransferase | MpaG *[P. brevicompactum]* | 99.3 | AJG44384.1 |
| MpaH’ | 398 | Hydrolase | MpaH *[P. brevicompactum]* | 89.8 | AJG44385.1 |

**Table S2.** The plasmids used in this study

| Plasmid | Relevant characteristics | Reference |
| --- | --- | --- |
| pTAex3 | Fungal expression vector that harbors the *α*-amylase promoter/terminator ($P_{amyB}$/$T_{amyB}$) and the *argB* selective marker. | (23, 24) |
| pAcGFP1 | DNA template of *gfp* sequence. | (25) |
| pANIC6D | DNA template of *rfp* sequence. | (26) |
| pTAex3-*mpaA'* | pTAex3 harboring *mpaA'* for heterologous expression of MpaA' in *Ao*M-2-3. | This study |
| pTAex3-*mpaB'* | pTAex3 harboring *mpaB'* for heterologous expression of MpaB' in *Ao*M-2-3. | This study |
| pTAex3-*mpaDE'* | pTAex3 harboring *mpaDE'* for heterologous expression of MpaDE' in *Ao*M-2-3. | This study |
| pTAex3-*mpaA'-mpaB'* | pTAex3 harboring *mpaA'* and *mpaB'* for heterologous co-expression of MpaA' and MpaB' in *Ao*M-2-3. | This study |
| pTAex3-*mpaA'-mpaH'* | pTAex3 harboring *mpaA'* and *mpaH'* for heterologous co-expression of MpaA' and MpaH' in *Ao*M-2-3. | This study |
| pTAex3-*mpaA'-mpaB'-mpaH'* | pTAex3 harboring *mpaA', mpaB', and mpaH'* for heterologous co-expression of MpaA', MpaB', and MpaH' in *Ao*M-2-3. | This study |
| pTAex3-*mpaA'-mpaB'-mpaH'*$^{\Delta GKL}$ | pTAex3 harboring *mpaA', mpaB' and mpaH'*$^{\Delta GKL}$ for heterologous co-expression of MpaA', MpaB', and MpaH'$^{\Delta GKL}$ in *Ao*M-2-3. | This study |
| pTAex3-*gfp-mpaA'* | pTAex3 harboring the *N-gfp-mpaA'-C* fusion gene for studying the subcellular location of MpaA'. | This study |
| pTAex3-*mpaB'-gfp* | pTAex3 harboring the *N-mpaB'-gfp-C* fusion gene for studying the subcellular location of MpaB'. | This study |
| pTAex3-*mpaDE'-gfp* | pTAex3 harboring the *N-mpaDE'-gfp-C* fusion gene for studying the subcellular location of MpaDE'. | This study |
| pTAex3-*gfp-mpaH'* | pTAex3 harboring the *N-gfp-mpaH'-C* fusion gene for studying the subcellular location of MpaH'. | This study |

| | | |
|---|---|---|
| pTAex3-$rfp^{SKL}$ | pTAex3 harboring $rfp^{SKL}$ for localization of RFP to peroxisome. | This study |
| pTAex3-*gfp*-$mpaH'^{\Delta GKL}$ | pTAex3 harboring *N*-*gfp*-$mpaH'^{\Delta GKL}$-*C* for verifying the role of the *C*-terminal GKL sequence in MpaH' for its peroximal localization. | This study |
| pTAex3-*gfp* | pTAex3 harboring *gfp* for heterologous expression of GFP in $Ao_{\text{M-2-3}}$. | This study |
| pRSFD-*hygB* | pRSFduet harboring the *hygB* resistance gene cassette. | This study |
| pRSFD-*hygB*-*mpaB'* | pRSFduet harboring the *hygB* resistance gene cassette which is flanked by the upstream and downstream sequences of *mpaB'* for in-frame deletion of *mpaB'* in $Pb_{864}$ by split-marker recombination strategy | This study |
| pRSFD-*hygB*-*mpaH'* | pRSFduet harboring the *hygB* resistance gene cassette which is flanked by the upstream and downstream sequences of *mpaH'* for in-frame deletion of *mpaH'* in $Pb_{864}$ by split-marker recombination strategy | This study |
| pET28b-*mpaH'* | pET-28b harboring the codon-optimized *mpaH'* gene for heterologous expression of MpaH' in *E. coli* BL21(DE3). | This study |
| pET28b-$mpaH'_{S139A}$ | pET-28b harboring the $mpaH'_{S139A}$ mutant gene for heterologous expression of $\text{MpaH'}_{\text{S139A}}$ mutant in *E. coli* BL21(DE3). | This study |

**Table S3.** The strains used in this study

| Strain | Relevant characteristics |
|---|---|
| *Pb*$_{864}$ | *P. brevicompactum* NRRL 864, wild type MPA producer |
| *Pb*$_{864}$-*ΔmpaB'* | The *mpaB'* deletion mutant of *Pb*$_{864}$ |
| *Pb*$_{864}$-*ΔmpaH'* | The *mpaH'* deletion mutant of *Pb*$_{864}$ |
| *Ao*$_{M\text{-}2\text{-}3}$-*mpaA'* | The *Ao*$_{M\text{-}2\text{-}3}$ strain transformed by pTAex3-*mpaA'* for heterologous expression of MpaA'. |
| *Ao*$_{M\text{-}2\text{-}3}$-*mpaB'* | The *Ao*$_{M\text{-}2\text{-}3}$ strain transformed by pTAex3-*mpaB'* for heterologous expression of MpaB'. |
| *Ao*$_{M\text{-}2\text{-}3}$-*mpaH'* | The *Ao*$_{M\text{-}2\text{-}3}$ strain transformed by pTAex3-*mpaH'* for heterologous expression of MpaH'. |
| *Ao*$_{M\text{-}2\text{-}3}$-*mpaH'*$^{\Delta GKL}$ | The *Ao*$_{M\text{-}2\text{-}3}$ strain transformed by pTAex3-*mpaH'*$^{\Delta GKL}$ for heterologous expression of MpaH'$^{\Delta GKL}$. |
| *Ao*$_{M\text{-}2\text{-}3}$-*mpaA'-mpaB'* | The *Ao*$_{M\text{-}2\text{-}3}$ strain transformed by pTAex3-*mpaA'-mpaB'* for heterologous co-expression of MpaA' and MpaB'. |
| *Ao*$_{M\text{-}2\text{-}3}$-*mpaA'-mpaH'* | The *Ao*$_{M\text{-}2\text{-}3}$ strain transformed by pTAex3-*mpaA'-mpaH'* for heterologous co-expression of MpaA' and MpaH'. |
| *Ao*$_{M\text{-}2\text{-}3}$-*mpaA'-mpaB'-mpaH'* | The *Ao*$_{M\text{-}2\text{-}3}$ strain transformed by pTAex3-*mpaA'-mpaB'-mpaH'* for heterologous co-expression of MpaA', MpaB', and MpaH'. |
| *Ao*$_{M\text{-}2\text{-}3}$-*mpaA'-mpaB'-mpaH'*$^{\Delta GKL}$ | The *Ao*$_{M\text{-}2\text{-}3}$ strain transformed by pTAex3-*mpaA'-mpaB'-mpaH'*$^{\Delta GKL}$ for heterologous co-expression of MpaA', MpaB', and MpaH'$^{\Delta GKL}$. |
| *Ao*$_{M\text{-}2\text{-}3}$-*gfp-mpaA'* | The *Ao*$_{M\text{-}2\text{-}3}$ strain transformed by pTAex3-*gfp-mpaA'* for heterologous expression of the *N*-GFP-MpaA'-*C* fusion protein. |
| *Ao*$_{M\text{-}2\text{-}3}$-*mpaB'-gfp* | The *Ao*$_{M\text{-}2\text{-}3}$ strain transformed by pTAex3-*mpaB'-gfp* for heterologous expression of the *N*-MpaB'-GFP-*C* fusion protein. |
| *Ao*$_{M\text{-}2\text{-}3}$-*mpaDE'-gfp* | The *Ao*$_{M\text{-}2\text{-}3}$ strain transformed by pTAex3-*mpaDE'-gfp* for heterologous expression of the *N*-MpaDE'-GFP-*C* fusion protein. |
| *Ao*$_{M\text{-}2\text{-}3}$-*gfp-mpaH'/rfp*$^{SKL}$ | The *Ao*$_{M\text{-}2\text{-}3}$ strain co-transformed by pTAex3-*gfp-mpaH'* and pTAex3-*rfp*$^{SKL}$ for heterologous co-expression of the *N*-GFP-MpaH'-*C* fusion protein and RFP$^{SKL}$ that carries the *C*-terminal SKL sequence for localization of RFP to peroxisome. |
| *Ao*$_{M\text{-}2\text{-}3}$-*gfp-mpaH'*$^{\Delta GKL}$ | The *Ao*$_{M\text{-}2\text{-}3}$ strain transformed by pTAex3-*gfp-mpaH'*$^{\Delta GKL}$ for heterologous expression of the *N*-GFP-MpaH'$^{\Delta GKL}$-*C* fusion protein. |
| *Ao*$_{M\text{-}2\text{-}3}$-*gfp* | The *Ao*$_{M\text{-}2\text{-}3}$ strain transformed by pTAex3-*gfp* for heterologous expression GFP. |

| | |
|---|---|
| *Ao*$_{M\text{-}2\text{-}3}$-*rfp* | The *Ao*$_{M\text{-}2\text{-}3}$ strain transformed by pTAex3-*rfp* for heterologous expression GFP. |
| *Ao*$_{M\text{-}2\text{-}3}$-*pTAex3* | The *Ao*$_{M\text{-}2\text{-}3}$ strain transformed by pTAex3 as a control. |

**Table S4.** Structures and HRMS information of MPA derivatives

| Name | Structure | Formula | Calculated ([M+H]$^+$) | Observed ([M+H]$^+$) |
|---|---|---|---|---|
| DHMP | 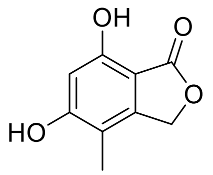 | $C_9H_8O_4$ | 181.0501 | 181.0503 |
| MPA | 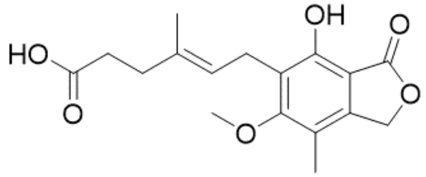 | $C_{17}H_{21}O_6$ | 321.1332 | 321.1323 |
| DMMPA | 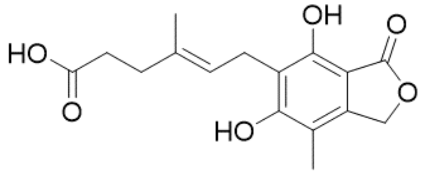 | $C_{16}H_{19}O_6$ | 307.1176 | 307.1173 |
| mycophenolic aldehyde | 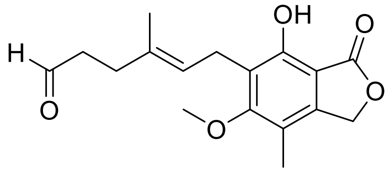 | $C_{17}H_{20}O_5$ | 305.1389 | 305.1376 |
| FDHMP | 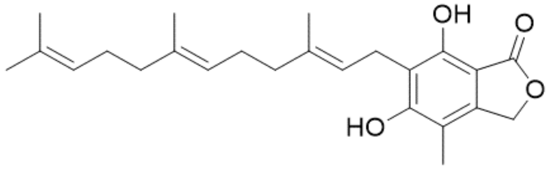 | $C_{24}H_{32}O_4$ | 385.2379 | 385.2371 |
| M-FDHMP | 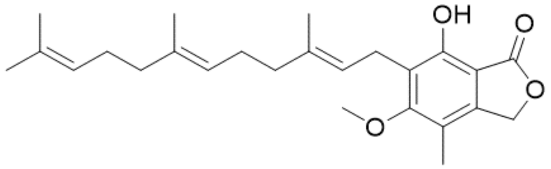 | $C_{25}H_{34}O_4$ | 399.2530 | 399.2532 |
| FDHMP-3C | 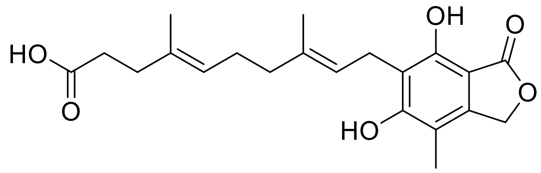 | $C_{21}H_{26}O_6$ | 375.1808 | 375.1808 |

| | | | | |
|---|---|---|---|---|
| FDHMP-d1 | 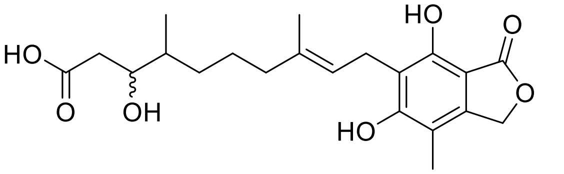 | $C_{21}H_{28}O_7$ | 393.1913 | 393.1914 |
| FDHMP-d2 | 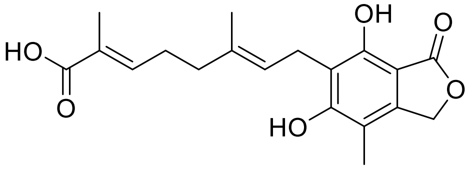 | $C_{19}H_{22}O_6$ | 347.1495 | 347.1499 |
| FDHMP-d3 | 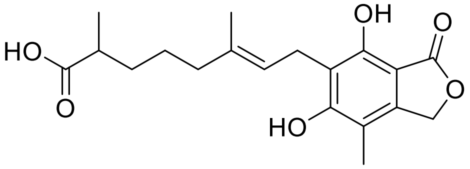 | $C_{19}H_{24}O_6$ | 349.1651 | 349.1650 |
| FDHMP-d4 | 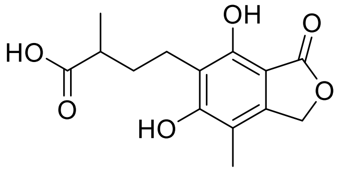 | $C_{14}H_{16}O_6$ | 281.1019 | 281.1007 |
| FDHMP-d5 | 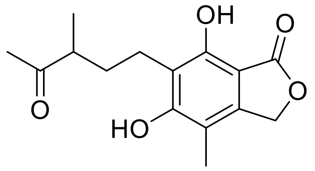 | $C_{15}H_{18}O_5$ | 279.1227 | 279.1221 |
| MFDHMP-d4 | 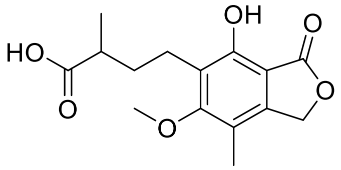 | $C_{15}H_{18}O_6$ | 295.1168 | 295.1176 |
| MFDHMP-d5 | 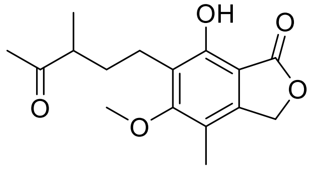 | $C_{16}H_{21}O_5$ | 293.1384 | 293.1387 |

**Table S5.** $^{1}H$ (600 MHz) and $^{13}C$ (150 MHz) NMR data of DHMP ($CD_3OD$) and mycophenolic aldehyde ($CDCl_3$)

| | **DHMP** | | **mycophenolic aldehyde** |
|---|---|---|---|
| Position | $\delta_C$, Type | $\delta_H$, mult., (*J* in Hz) | $\delta_H$, mult., (*J* in Hz) |
| 1 | 172.0, s | | |
| 2 | 102.1, s | | |
| 3 | 156.1, s | | |
| 4 | 101.5, d | 6.34, s | |
| 5 | 162.8, s | | |
| 6 | 109.6, s | | |
| 7 | 148.8, s | | |
| 8 | 68.8, t | 5.16, s | 5.16, s |
| 9 | 8.90, q | 2.01, s | 2.22, s |
| 10 | | | 3.40, d, (6.9) |
| 11 | | | 5.12, t, (6.4) |
| 12 | | | |
| 13 | | | 2.29, t, (7.5) |
| 14 | | | 2.49, td, (7.6, 1.5) |
| 15 | | | 9.71, t, (1.4) |
| 16 | | | 1.78, s |

**Table S6.** $^1$H (600 MHz) and $^{13}$C (150 MHz) NMR data of FDHMP ($CDCl_3$) and MFDHMP ($CDCl_3$)

| | **FDHMP** | | **MFDHMP** | |
|---|---|---|---|---|
| Position | $\delta_C$, type | $\delta_H$, mult., (*J* in Hz) | $\delta_C$, type | $\delta_H$, mult., (*J* in Hz) |
| 1 | 173.3, s | | 173.0, s | |
| 2 | 102.8, s | | 106.3, s | |
| 3 | 152.9, s | | 153.7, s | |
| 4 | 112.8, s | | 122.6, s | |
| 5 | 160.6, s | | 163.7, s | |
| 6 | 110.5, s | | 116.7, s | |
| 7 | 144.1, s | | 143.9, s | |
| 8 | 69.8, t | 5.19, s | 70.0, t | 5.19, s |
| 9 | 10.8, q | 2.00, s | 11.5, q | 2.14, s |
| 10 | 21.7, t | 3.45, d, (7.1) | 22.6, t | 3.39, d, (6.9) |
| 11 | 120.4, d | 5.22, t, (6.7) | 121.8, d | 5.20, t, (overlapped) |
| 12 | 140.7, s | | 135.9, s | |
| 13 | 39.6, t | 2.10, m; 1.96, m | 39.7, t | 1.92, m; 1.99, m |
| 14 | 26.6, t | 2.02, m | 26.7, t | 1.99, m |
| 15 | 123.3, d | 5.06, t, (overlapped) | 124.0, d | 5.06, t, (6.3) |
| 16 | 135.8, s | | 134.6, s | |
| 17 | 39.6, t | 2.13, m; 2.08, m | 39.7, t | 1.92, m; 1.99, m |
| 18 | 26.0, t | 2.09, m, (overlapped) | 26.5, t | 2.06, m |
| 19 | 124.3, d | 5.06, t, (overlapped) | 124.3, d | 5.06, t, (6.3) |
| 20 | 131.3, s | | 131.2, s | |
| 21 | 25.6, q | 1.66, s | 25.7, q | 1.66, s |
| 22 | 16.3, q | 1.84, s | 16.2, q | 1.79, s |
| 23 | 16.0, q | 1.58, s | 16.0, q | 1.56, s |
| 24 | 17.6, q | 1.59, s | 17.6, q | 1.57, s |
| 25 | | | 61.0, q | 3.77, s |

**Table S7.** $^{1}H$ (600 MHz) and $^{13}C$ (150 MHz) NMR data of FDHMP-3C ($CD_3OD$) and FDHMP-d1 ($CD_3OD$)

| | **FDHMP-3C** | | **FDHMP-d1** | |
|---|---|---|---|---|
| Position | $\delta_C$, type | $\delta_H$, mult., (*J* in Hz) | $\delta_C$, type | $\delta_H$, mult., (*J* in Hz) |
| 1 | 174.6, s | | 174.6, s | |
| 2 | 103.5, s | | 103.6, s | |
| 3 | 154.4, s | | 154.5, s | |
| 4 | 117.1, s | | 117.2, s | |
| 5 | 161.9, s | | 161.9, s | |
| 6 | 111.0, s | | 111.1, s | |
| 7 | 146.1, s | | 146.1, s | |
| 8 | 70.8, t | 5.16, s | 70.8, t | 5.19, s |
| 9 | 11.3, q | 2.05, s | 11.3, q | 2.06, s |
| 10 | 22.9, t | 3.35, d, (7.2) | 22.9, t | 3.38 d, (7.0) |
| 11 | 123.6, d | 5.16, t, (overlapped) | 123.5, d | 5.19, t, (overlapped) |
| 12 | 135.6, s | | 136.1, s | |
| 13 | 40.6, t | 1.96, t, (7.1) | 40.8, t | 1.96, m |
| 14 | 27.1, t | 2.05, m | 26.2, t | 1.37, m; 1.51, m |
| 15 | 125.7, d | 5.04, t, (6.7) | 33.2, t | 1.03, m; 1.37, m |
| 16 | 134.6, s | | 39.3, d | 1.50, m |
| 17 | 35.7, t | 2.12, m | 73.4, d | 3.82, m |
| 18 | 33.8, t | 2.16, m | 39.3, t | 2.23, m; 2.28, m |
| 19 | 177.3, s | | 177.4, s (from HMBC) | |
| 20 | 16.3, q | 1.77, s | 16.1, q | 1.77, s |
| 21 | 16.0, q | 1.53, s | 16.0, q | 0.85, d, (6.8) |

**Table S8.** $^{1}H$ (600 MHz) and $^{13}C$ (150 MHz) NMR data of FDHMP-d2 ($CD_3OD$) and FDHMP-d3 ($CD_3OD$)

| | **FDHMP-d2** | | **FDHMP-d3** | |
|---|---|---|---|---|
| Position | $\delta_C$, Type | $\delta_H$, mult., (*J* in Hz) | $\delta_C$, Type | $\delta_H$, mult., (*J* in Hz) |
| 1 | 174.7, s (from HMBC) | | 174.6, s | |
| 2 | 103.6, s | | 103.6,s | |
| 3 | 154.6, s (from HMBC) | | 154.4, s | |
| 4 | 117.0, s | | 117.1, s | |
| 5 | 160.7, s | | 161.9, s | |
| 6 | 111.1, s | | 111.1, s | |
| 7 | 146.2, s | | 146.1, s | |
| 8 | 70.8, t | 5.22, s | 70.7, t | 5.21, s |
| 9 | 11.3, q | 2.09, s | 11.3, q | 2.09, s |
| 10 | 22.9, t | 3.41, d, (7.0) | 22.8, t | 3.40, d, (7.1) |
| 11 | 124.2, d | 5.27, t, (6.9) | 123.5, d | 5.22, t, (overlapped) |
| 12 | 135.2, s | | 136.0, s | |
| 13 | 39.3, t | 2.11, m | 40.6, t | 1.98, t, (7.3) |
| 14 | 28.2, t | 2.30, dd, (7.2, 14.4) | 26.5, t | 1.42, m |
| 15 | 143.6, d | 6.72, t, (7.1) | 34.4, t | 1.35, m; 1.56, m |
| 16 | 129.0, s (from HMBC) | | 40.5, d | 2.39, m |
| 17 | 171.5, s | | 171.5, s | |
| 18 | 16.3, q | 1.83, s | 16.1, q | 1.78, s |
| 19 | 12.4, q | 1.76, s | 17.6, q | 1.10, d, (7.0) |

**Table S9.** $^{1}H$ (600 MHz) and $^{13}C$ (150 MHz) NMR data of MFDHMP-d4 ($CD_3CN$) and MFDHMP-d5 ($CD_3OD$)

| | **MFDHMP-d4** | | **MFDHMP-d5** | |
|---|---|---|---|---|
| Position | $\delta_C$, Type | $\delta_H$, mult., (*J* in Hz) | $\delta_C$, Type | $\delta_H$, mult., (*J* in Hz) |
| 1 | 173.4, s | | 173.7, s | |
| 2 | 107.4, s | | 107.7, s | |
| 3 | 154.1, s | | 154.7, s | |
| 4 | 118.1, s | | 117.9, s | |
| 5 | 164.6, s | | 164.9, s | |
| 6 | 123.1, s | | 117.9, s | |
| 7 | 146.3, s | | 146.8, s | |
| 8 | 71.0, t | 5.22, s | 70.8, t | 5.23, s |
| 9 | 11.7, q | 2.12, s | 11.4, q | 2.15, s |
| 10 | 22.0, t | 2.67, t, (8.0) | 22.2, t | 2.63, ddd, (8.9, 6.7, 2.2) |
| 11 | 34.0, t | 1.86, m; 1.62, m | 33.4, t | 1.91, m; 1.57, m |
| 12 | 39.7, d | 2.46, m | 47.9, d | 2.59, m |
| 13 | 178.0, s | | 215.4, s | |
| 14 | 17.3, q | 1.18, d, (7.0) | 28.1, q | 2.16, s |
| 15 | 61.8, q | 3.77, s | 16.7, q | 1.12, d, (7.0) |
| 16 | | | 61.6, q | 3.77, s |

**Table S10.** The steady-state kinetic parameters of twelve acyl-CoA esters

| Acyl-CoA esters | $k_{cat}$ ($min^{-1}$) | $K_m$ (μM) | $k_{cat}/K_m$ ($\mu M^{-1}$ $min^{-1}$) |
|---|---|---|---|
| Acetyl-CoA | 54.98 ± 6.139 | 722.3 ± 213.9 | 0.076 |
| Propionyl-CoA | 24.58 ± 2.443 | 351.5 ± 130.9 | 0.070 |
| Malonyl-CoA | 12.94 ± 1.295 | 76.37 ± 52.11 | 0.169 |
| Isobutyryl-CoA | 7.688 ± 0.3602 | 201.9 ± 45.03 | 0.038 |
| Isovaleryl-CoA | 5.718 ± 1.068 | 780.8 ± 372.8 | 0.007 |
| Benzoyl-CoA | 3.891 ± 0.3079 | 217.9 ± 79.63 | 0.018 |
| N-Decanoyl-CoA | 136.5 ± 9.288 | 143 ± 52.66 | 0.955 |
| Lauroyl-CoA | 99.11 ± 7.066 | 260.2 ± 79.56 | 0.381 |
| Palmitoyl-CoA | 23.19 ± 2.574 | 577 ± 189.20 | 0.040 |
| Arachidonoyl-CoA | 47.35 ± 3.873 | 316.6 ± 101.8 | 0.150 |
| DMMPA-CoA | 4438 ± 494.3 | 382.6 ± 153.4 | 11.6 |
| MPA-CoA | 9578 ± 389.2 | 117.5 ± 32.93 | 81.5 |

**Table S11.** Primers used in this study

| Primers | Sequence (5'-3') |
|---|---|
| pET28b-MpaH'-*Nde*I-FP | CCTGGTGCCGCGCGGCAGC*CATATG*TCAACTGAGAAATTTAC |
| pET28b-MpaH'-*Nde*I-RP | GTCCACCAGTCATGCTAGC*CATATG*CTAAAGTTTTCCCTTATCAT |
| MpaH-S139A-FP | GGCCACGCTTTTGGTGGCAATATTATTACCAACCT |
| MpaH-S139A-RP | ACCAAAAGCGTGGCCGATTCCAACTAGAGGACGTG |
| pTAex3-GFP-*Nde*I-FP | CGCGCGGCAGCGAGCTC*CATATG*GTGAGCAAGGGCGCCG |
| pTAex3-GFP-*Kpn*I-RP | GCTACTACAGATCCCCG*GGTACC*CTACTTGTACAGCTCATCC |
| pTAex3-RFP-*Nde*I-FP | CGCGCGGCAGCGAGCTC*CATATG* GCTCTTTCAAAGCAAA |
| pTAex3-RFP-*Kpn*I-RP | GCTACTACAGATCCCCG*GGTACC*TTAGTGATGGTGATGGTGA |
| pTAex3-RFPskl-*Kpn*I-RP | GCTACTACAGATCCCCG*GGTACC*TTACAGCTTCGAGGCTACCGGT<br>AAGTTAGAACTACAGATCCCCGGGTACCCTAATGGAAGGGACATT<br>TCC |
| pTAex3-MpaA'-NdeI-FP | GCGGCAGCGAGCTC*CATATG*ATGACCAACGCAGTGGAG |
| pTAex3-MpaA'-KpnI-RP | ACTACAGATCCCCG*GGTACC*TCAAAGCTTAATGTACCCCTC |
| pTAex3-MpaB'-NdeI-FP | GCGGCAGCGAGCTC*CATATG*ATGTCTTTGTCTTTGCCTCC |
| pTAex3-MpaB'-KpnI-RP | ACTACAGATCCCCG*GGTACC*CTAATGGAAGGGACATTTCC |
| pTAex3-MpaH'-NdeI-FP | CGCGCGGCAGCGAGCTC*CATATG*TCAACTGAGAAATTTAC |
| pTAex3-MpaH'-KpnI-RP | GCTACTACAGATCCCCG*GGTACC*CTAAAGTTTTCCCTTATCAT |
| pTAex3-MpaH'ΔGKL -KpnI-RP | GCTACTACAGATCCCCG*GGTACC*CTACTTATCATTCCTCTTC |
| nGFPMpaH'-WL-FP | GGTGGTGGTGGTGGCTGGATCATGTCAACTGAGAAATTTAC |
| nGFPMpaH'-WL-RP | ATGGAGCTCGCTGCCGCGC |
| nGFPMpaH'-FP | GTGCCGCGCGGCAGCGAGCTC*CATATG*GTGAGCAAGGGCGCCG |
| nGFPMpaH'-RP | GATCCAGCCACCACCACCACCCTTGTACAGCTCATCCATG |
| nRFP MpaH'-FP | GTGCCGCGCGGCAGCGAGCTCCATATGGCTCTTTCAAAGCAAA |
| nRFP MpaH'-RP | GATCCAGCCACCACCACCACCGTGATGGTGATGGTGATGG |
| nGFPMpaB'-WL-FP | GGTGGTGGTGGTGGCTGGATCATGTCTTTGTCTTTGC CTCC |
| MpaB'cGFP-FP | GGTGGTGGTGGTGGCTGGATCATGGTGAGCAAGGGCGCCG |
| MpaB'cGFP-RP | ACTACAGATCCCCGGGTACCCTACTTGTACAGCTCATCCATG |
| MpaB'cGFP-WL-FP | GATCCAGCCA CCACCACCACCATGGAAGGGACATTTCCC |
| MpaB'cGFP-WL-RP | *GGTACC*CGGGGATCTG TAGT |
| MpaA'nGFP-WL-FP | GGTGGTGGTGGTGGCTGGATCATGACCAACGCAGTGGAG |
| MpaA'cGFP-WL-FP | GATCCAGCCA CCACCACCACCAAAGCTTAAT GTACCCCT |
| pTAex3-agrB-FP | CAGTGCCAAGCTTGCATGCCTGCAGGTC |
| pTAex3-Tamy-RP | CTGGAAAGCGGGCAGTGAGCGCAACG |
| pTAex3-YZ-FP | CTCTTCTGC GAATCGCTTG |
| pTAex3-YZ-RP | GACAGCAGTAACGACTCCAA |
| PRSF -H-B-UP-FP | ACGCGTCGACAATCACGAAGTGGCAGGACTGGAAA |
| PRSF -H-B-UP-RP | CCCAAGCTTACTTGGACGTTATGTCTTGTAGAGC |
| PRSF -H-B-DN-FP | CGGGGTACCAGTTTGGGCTTCTGAGATTACCTGC |
| PRSF -H-B-FN-RP | CCGCTCGAGGTCCAGCCCAGACTCAGACCCTTCT |
| QF121F | ACGCGTCGACAATCACGAAGTGGCAGGACTGGAAA |
| QF121R | CCCAAGCTTACTTGGACGTTATGTCTTGTAGAGC |
| QF122F | CGGGGTACCAGTTTGGGCTTCTGAGATTACCTGC |
| QF122R | CCGCTCGAGGTCCAGCCCAGACTCAGACCCTTCT |
| QF126F | CTTCTACACAGCCATCGGTCCA |
| QF126R | TGAGCGCCCCTACAGATACCAC |
| QF127F | TGGTGGCTGTATGCGGTGTTGA |
| QF128R | TACGCTTGGTGGCACAGTTCTCG |
| QF128F | GCTTGCCGTCTGTGGTATCTGT |
| QF127R | ATCTTGGATTCTGTCGGCTTGG |
| QF129F | ATGTTCTTGACATCTGGGCAGTTTG |
| QF129R | TATCACATACAACATCGCCTTCCTC |
| MPAH-UP-F | ACGCGTCGACCATGTCTCCAGTTTCGTAAGATTTA |
| MPAH-UP-R | CCCAAGCTTTTGCACGTTACAAGCTGAAAGGAGT |
| MPAH-DOWN-F | GGAAGATCTGACAAATCAGGCACCAAGGCTC |
| MPAH-DOWN-R | CGGGGTACCGCTCAGCAGCCCAGCCAGTCAG |
| H-ANCHOR-UP-F | CGACCTACAGTGACCCAACAAG |
| PRSF-ANCHOR-R | TACGCTTGGTGGCACAGTTCTCG |
| PRSF-ANCHOR-F | GCTTGCCGTCTGTGGTATCTGT |
| H-ANCHOR-DOWN-R | ACAACCCGAATCTCAATACAGC |

| | |
|---|---|
| H-INACHOR-F | GGAGTCGGTCTGCCAAAGGTGA |
| H-NACHOR-R | ACCTCCGCTCCCACCAATACCC |
| MpaA-FP | CTACTGATCAGCACCTCGCG |
| MpaA-RP | TCCACCTTCTGCAATCCGAC |
| MpaB-FP | CACTTTCTGAGCTTGCGAGG |
| MpaB-RP | TCCTGGATTTCCTTGTGGCC |
| MpaH-FP | TACCATCACTGAGCACCTGG |
| MpaH-RP | CCCGGGTACCCTAAAGTTTTCC |
| MpaHCT-RP | TGCGTCCACTAGGAATTGGC |
| MpaAopt-FP | ACTAACGCCGTCGAGGATTC |
| MpaAopt-RP | AGCTTGATGTAGCCCTCGAC |
| MpaBopt-FP | CTCTCTCTCTCCCTCCTGCC |
| MpaBopt-RP | CGGTACGGGAGATCTTGAGC |
| MpaHopt-FP | CCACCGAGAAGTTCACCATC |
| MpaHCTopt-RP | CTTATCGTTACGCTTCTTGCGG |
| gfp-FP | ATGGTGAGCAAGGGCGC |
| gfp-RP | CTTGTACAGCTCATCCATG |
| rfp-FP | ATGGCTCTTTCAAAGCAAAG |
| rfp-RP | TTAGTGATGGTGATGGTGAT |
| gfp-mpaA’-RP | TCAAAGCTTAATGTACCCC |
| mpaB'-gfp-FP | ATGTCTTTGTCTTTGCCTCC |
| gfp-mpaH'-RP | CTAAAGTTTTCCCTTATC |
| gfp-mpaH'-RP | CTACTTATCATTCCTCTTC |
| mpaDE'-FP | GAGTCTTTGTCGCTAACATG |
| mpaDE'-RP | TTACTTCTGTCCTTCTATGG |
| PamyB-FP | TCATGGTGTTTTGATCATTT |
| TamyB-RP | CTGGAAAGCGGGCAGTGAGC |

*Note*: 1. Restriction sites are shown in italic; 2. FP and RP stand for forward primer and reverse primer, respectively.
